# Fab-dimerized glycan-reactive antibodies neutralize HIV and are prevalent in humans and rhesus macaques

**DOI:** 10.1101/2020.06.30.178921

**Authors:** Wilton B. Williams, R. Ryan Meyerhoff, RJ Edwards, Hui Li, Nathan I. Nicely, Rory Henderson, Ye Zhou, Katarzyna Janowska, Katayoun Mansouri, Kartik Manne, Victoria Stalls, Allen L. Hsu, Mario J. Borgnia, Guillaume Stewart-Jones, Matthew S. Lee, Naomi Bronkema, John Perfect, M. Anthony Moody, Kevin Wiehe, Todd Bradley, Thomas B. Kepler, S. Munir Alam, Robert J. Parks, Andrew Foulger, Mattia Bonsignori, Celia C. LaBranche, David C. Montefiori, Michael Seaman, Sampa Santra, Joseph R. Francica, Geoffrey M. Lynn, Baptiste Aussedat, William E. Walkowicz, Richard Laga, Garnett Kelsoe, Kevin O. Saunders, Daniela Fera, Peter D. Kwong, Robert A. Seder, Alberto Bartesaghi, George M. Shaw, Priyamvada Acharya, Barton F. Haynes

## Abstract

The HIV-1 envelope (Env) is comprised by mass of over 50% glycans. A goal of HIV-1 vaccine development is the induction of Env glycan-reactive broadly neutralizing antibodies (bnAbs). The 2G12 bnAb recognizes an Env glycan cluster using a unique variable heavy (V_H_) domain-swapped conformation that results in fragment antigen-binding (Fab) dimerization. Here we describe Fab-dimerized glycan (FDG)-reactive antibodies without V_H_-swapped domains from simian-human immunodeficiency virus (SHIV)-infected macaques that neutralized heterologous HIV-1 isolates. FDG precursors were boosted by vaccination in macaques, and were present in HIV-1-naïve humans with an average estimated frequency of one per 340,000 B cells. These data demonstrate frequent HIV-1 Env glycan-reactive bnAb B cell precursors in macaques and humans and reveal a novel strategy for their induction by vaccination.

**Highlights:**

- Discovery of Fab-dimerized HIV-1 glycan-reactive antibodies with a non-domain-swapped architecture
- Fab-dimerized antibodies neutralize heterologous HIV-1 isolates.
- Antibodies with this architecture can be elicited by vaccination in macaques.
- Fab-dimerized antibodies are found in HIV-1 naïve humans.

## Introduction

A glycan shield covers the HIV-1 envelope (Env) and limits antibody access to broadly reactive neutralizing antibody (bnAb) epitopes. Thus, a major goal of HIV-1 vaccine development is to target the Env glycan shield with an easily-induced glycan-targeted neutralizing antibody response (*1*). HIV-1 Env glycans are predominantly high mannose in character (*2–7*). Since HIV-1 Env glycans are host-derived, they mimic glycans on host cells; thus HIV-1 Env immunogenicity may be limited by host tolerance controls (*1, 8–11*).

There are multiple examples of HIV-1 Env glycopeptide-reactive HIV-1 bnAbs that use long complementary-determining region (CDR) loops to penetrate the glycan shield and contact protein residues in Env, while simultaneously making stabilizing contacts with the surrounding high mannose glycans (*12–21*). In contrast, antibody 2G12, one of the first bnAbs isolated, has been the only example to date of an HIV-1 Env glycan-reactive bnAb that interacts solely with high mannose glycans (*22*). The 2G12 bnAb has a short variable heavy (V_H_) domain CDR3 loop, but has a unique domain-swapped architecture in which the V_H_ domains of the two fragment antigen-binding (Fab) arms of the 2G12 IgG swap to create a Fab-dimerized multivalent glycan binding surface (*22–26*).

Carbohydrate antibodies typically bind glycans weakly with micromolar affinities (*27*). In contrast, 2G12 binds HIV-1 Env glycans with nanomolar affinity, primarily due to the unique glycan cluster architecture of the 2G12 epitope on HIV-1 Env (*28*), presented in a non-self-arrangement on HIV (*1*). However, glycan recognition by 2G12 is not limited to HIV-1 Env; 2G12 also binds to heat-killed *Candida albicans* yeast that present a similar Man(α1-2)Man motif as HIV-1 Env glycans (*23, 28, 29*). Moreover, yeast have been implicated in the induction of 2G12-like B cell responses in humans (*1*). However, since the 2G12 bnAb was isolated from a chronically HIV-1-infected individual (*30*), the ontogeny of 2G12 bnAb lineage is unknown.

Here we report non-V_H_ domain-swapped, Fab-dimerized, glycan-reactive (FDG) HIV-1 bnAbs that target the Env glycan shield. Precursors of FDG B cell lineages were common and resided in a mutated IgM B cell pool before virus infection or vaccination in macaques as well as in HIV-1 seronegative humans. Genetic, functional and structural characteristics of FDG antibodies demonstrated that they represent a new category of HIV-1-reactive antibodies that can undergo affinity maturation to neutralize heterologous HIV-1 strains.

## Results

### SHIV-induced DH851 FDG bnAb B cell lineage

The CH848 transmitted-founder (TF) virus initiated infection in an HIV-1-infected individual from whom a V3-glycan bnAb lineage was isolated (*20*). Four rhesus macaques were infected with CH848TF Env SHIV (*31*). In one macaque (RM6163), plasma antibodies developed after 20 weeks that neutralized the autologous tier 2 (difficult-to-neutralize) HIV-1 strain (**Figure S1A**). Heterologous tier 2 neutralization breadth developed after 80 weeks **(Figure S1B**). Tetramerized fluorophore-labeled HIV-1 CH848 soluble stabilized Env trimers (SOSIPs) (*32*), or synthetic HIV-1 Env glycopeptide (Man_9_-V3) (*33*) that mimics a V3-glycan bnAb epitope, were used to flow sort SHIV CH848TF-infected macaque lymph node memory B cells at week 52 after infection, as well as blood memory B cells at weeks 52 and 104 after infection. We isolated 173 CH848-Env reactive antibodies (**Table S1**), including two glycan-dependent neutralizing B cell lineages (DH851 and DH898) from lymph node and blood memory B cells at 52 and 104 weeks post-SHIV infection (**Table S2**).

A four-member clonal lineage comprised of one IgM and three IgG antibodies, termed DH851, was isolated from week 52 lymph node (IgG DH851.1-DH851.3) and week 104 blood (IgM DH851.4) memory B cells using tetramerized fluorophore-labeled Man_9_-V3 glycopeptide (*33*) or CH848TF SOSIPv4.1 (**Figure 1A**). DH851 antibodies used macaque Vλ2 gene paired with a V_H_2 gene encoding an HCDR3 of 15 amino acids (**Table S2**). When expressed as recombinant IgGs, DH851 monoclonal antibodies (mAbs) potently neutralized (Kifunensine [Kif]-treated) HIV-1 isolates bearing high mannose glycans (**Figure S2A**). However, only DH851 mAbs that were originally class-switched IgGs <u>in vivo</u> (DH851.1, DH851.2 and DH851.3) broadly neutralized non-Kif-treated wild-type autologous or heterologous tier 2 HIV-1 isolates, whereas DH851.4 (originally an IgM) did not (**Figure 1B**). DH851.1, DH851.2 and DH851.3 mAbs showed up to 26% neutralization breadth against a panel of 119 multiclade heterologous tier 2 HIV-1 strains (**Table S3**) and were dependent on N339, N386 and N392 glycans in the V3-glycan bnAb epitope for neutralization (**Figure S2B**).

**Figure 1.**
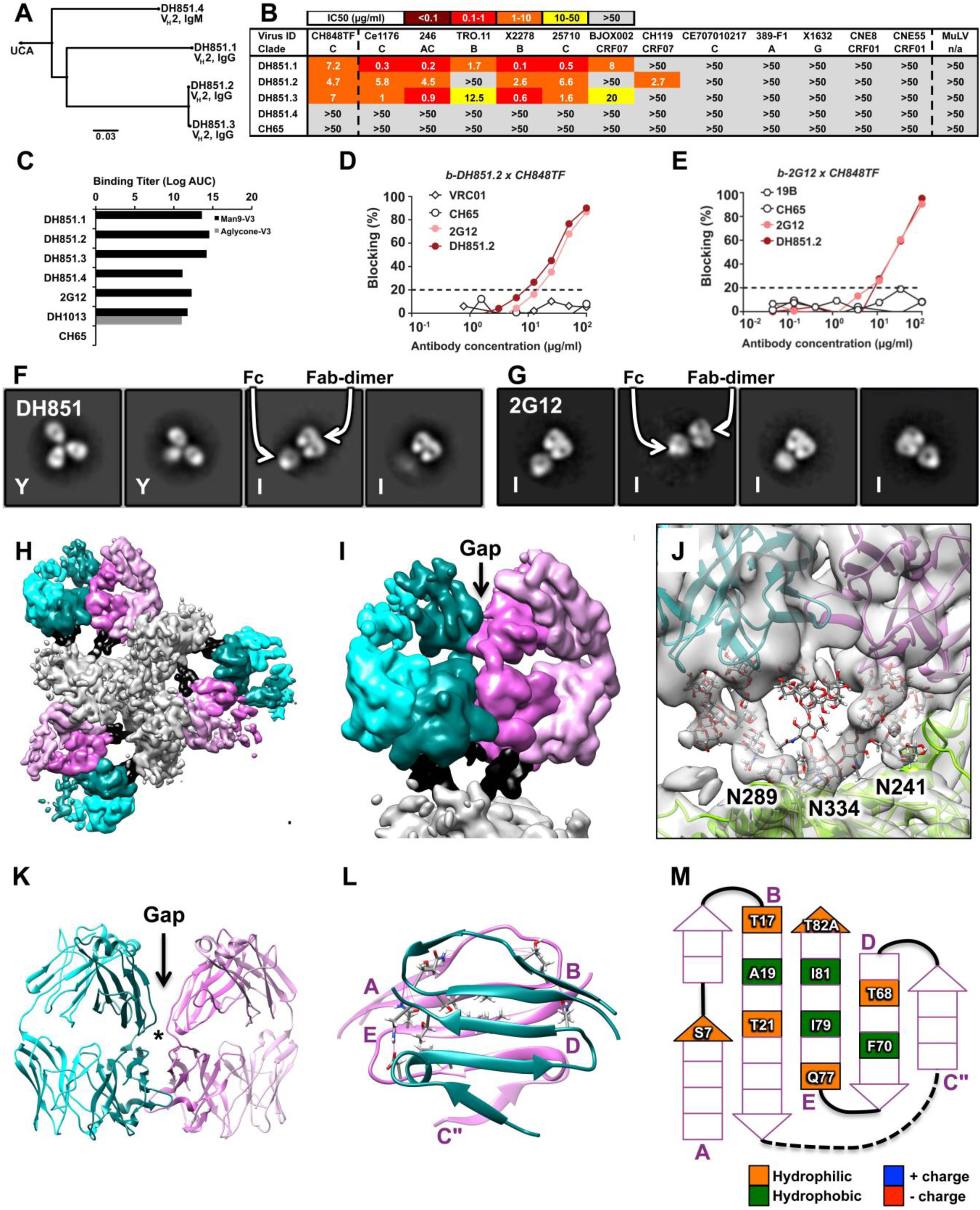
Characteristics of macaque CH848TF SHIV-induced Env glycan-reactive bnAb B cell lineage, DH851. **(A)** Phylogram and immunogenetics of DH851 bnAbs. **(B)** Neutralization of DH851 mAbs against autologous tier 2 CH848TF and the global panel of heterologous tier 2 HIV-1 strains. Data shown are representative of multiple neutralization assays tested in TZM-bl cells with different antibody lots, and neutralization titers were reported as IC50 in µg/ml. **(C)** ELISA binding titers (Log AUC) of DH851 mAbs to synthetic Man_9_-V3 glycopeptide that mimics a V3 glycan bnAb epitope (black bars) and the corresponding non-glycosylated aglycone V3 peptide (gray bars) (*33*). Reference antibodies tested included peptide-reactive DH1013, Env glycan-reactive 2G12, and negative control anti-influenza CH65 mAbs. Data shown are representative of four assays. Cross-blocking DH851 **(D)** and 2G12 **(E)** in ELISA. Data show percent blocking of biotinylated (b) DH851 and 2G12 binding to CH848TF gp120 monomeric protein by competing non-biotinylated mAbs at varying concentrations. 2G12 blocking on CH848TF gp120 was representative of three experiments. DH851 blocking on CH848TF was from a single experiment, but these results were similarly observed for blocking on CH848 10.17 gp120 in three separate experiments (not shown). **(F)** NSEM analysis of DH851 showing a mixture of Y-shaped (Y) and I-shaped (I) antibodies. **(G)** NSEM of 2G12 showing only I-shaped antibodies. **(H)** Segmented cryo-EM map of DH851.3 bound to stabilized CH505TF SOSIP trimer (*49*) showing six Fab domains arranged in three Fab-dimer pairs and binding to the SOSIP trimer with branched glycans well resolved. SOSIP trimer = gray; glycans = black; Fab heavy chains = teal or dark pink; Fab light chains = cyan or light pink. **(I)** Zoomed-in view of the segmented map showing a single Fab-dimer with a gap in the middle indicating that the two Fabs sit side-by-side and are not VH domain-swapped. **(J)** Zoomed-in view of the map shown as transparent surface with the fitted model shown in cartoon representation and glycans shown as sticks. **(K)** Atomic model of the Fab-dimer showing a gap between the Fabs (arrow) and an interface between the Fv domains (star). **(L)** The beta-sandwich view of the interface (starred region in K, turned 90°) shows the beta-strands of the two heavy chains are nearly parallel. Pink beta-strands are labeled A-E. **(M)** Schematic diagram of the Fab-dimer interface showing a patch of four hydrophobic or aromatic residues (green) surrounded by small hydrophilic residues (orange). Beta strands in the schematic correspond to the pink strands in (L) and are labeled A-E accordingly.

Like 2G12, DH851 mAbs bound HIV-1 CH848 SOSIP trimers (*32*) in a glycan-dependent manner; bound Man_9_-V3 glycopeptide, but not the corresponding non-glycosylated aglycone V3 peptide (*33*), and bound *Candida albicans* or *Cryptococcus neoformans* yeast glycans (**Figure 1C and S2C-F**). Both bnAbs 2G12 and IgG DH851.2 cross-blocked each other for binding CH848TF gp120 (**Figure 1D-1E**), suggesting that they bound overlapping epitopes.

Negative stain electron microscopy (NSEM) of DH851 IgGs (DH851.1-DH851.3) revealed a mixture of canonical Y-shaped antibodies as well as I-shaped Fab-dimerized antibodies (**Figure 1F and S3**) similar to the shape of the previously reported 2G12 bnAb (*22*) (**Figure 1G**). In contrast, neither the DH851 inferred unmutated common ancestor (UCA) nor the non-bnAb DH851 lineage member, DH851.4, had an I-shaped conformation (**Figure S3**). That the DH851 clonal lineage antibody DH851.4 did not neutralize wild-type HIV-1 (**Figure S2B**) suggested that Fab dimerization may be a prerequisite for HIV-1 neutralization.

To visualize the DH851 bnAb Fab-dimer interface, we solved a cryo-EM structure at 4.5-Å resolution of DH851.3 bound to a CH505TF SOSIP trimer (**Figures 1H-1I and S4; Table S4**). The cryo-EM reconstruction revealed three independent Fab-dimers bound to each Env trimer with the Fabs oriented side-by-side (**Figure 1H**). However, unlike 2G12, the Fab-dimers were not domain-swapped (**Figure 1I**). The Fab-dimers contacted Env glycans at N241, N289 and N334 in the V3 glycan super-site (*34*) mainly via heavy chain interactions (**Figure 1J**), which is consistent with the observed cross-blocking with 2G12 (**Figure 1D-1E**). The DH851 Fab-dimers also contacted the adjacent Env protomer via the light chain of the inner Fab (**Figure S5A-S5B**). The structure revealed a dimer interface between the two V_H_ domains (**Figure 1K**) that was similar to the 2G12 Fab-dimer interface (*22*); however, the V_H_ domains of DH851 were rotated such that the beta-strands of both chains were nearly parallel (**Figure 1L**), as opposed to being angled as seen in 2G12 (**Figure S6**).

Analysis of the DH851 antibody lineage sequences and contact residues revealed hydrophobic and aromatic residues within the dimer interface (**Figures 1M, S3K and S5C**) that were similar but not identical to those in the 2G12 Fab-dimer interface (**Figure S6D-S6E and S6K**). Antibodies DH851.4 and DH851UCA lacked these key hydrophobic residues (**Figure S3K**) and did not show Fab-dimerized structures by NSEM (**Figure S3F and S3J**). In particular, DH851.4 has a T81R mutation that would place a bulky charged residue in the dimer interface, analogous to the I19R mutation that disrupts V_H_ domain-swapping in 2G12 (*23*). These results indicated that FDG antibody neutralization breadth requires Fab-dimerization, which can be achieved without swapped V_H_ domains.

### SHIV-induced DH898 FDG Precursor B cell lineage

A second seven-member IgG antibody clonal lineage, DH898, was isolated from RM6163 lymph node memory B cells at 52 weeks post-infection using tetramerized fluorophore-labeled CH848TF SOSIP trimers (**Figure 2A**). DH898 antibodies used macaque Vκ2 gene paired with a V_H_1 gene encoding a 13 amino acid HCDR3 (**Table S2**). While DH898 mAbs neutralized autologous tier 2 wild-type CH848TF virus in a N332-dependent manner, only [Kif]-treated heterologous HIV-1 isolates were neutralized by DH898 mAbs (**Figure 2B and Table S3**). Similar to 2G12, DH898 mAbs bound Man_9_-V3 glycopeptide but not the aglycone V3 peptide (**Figures 1C and S7A**), and demonstrated binding to yeast glycans (**Figure S7A**). DH898 mAbs also mediated glycan-dependent binding to a CH848 SOSIP trimer (*32*) (**Figure S7B-S7C**).

**Figure 2.**
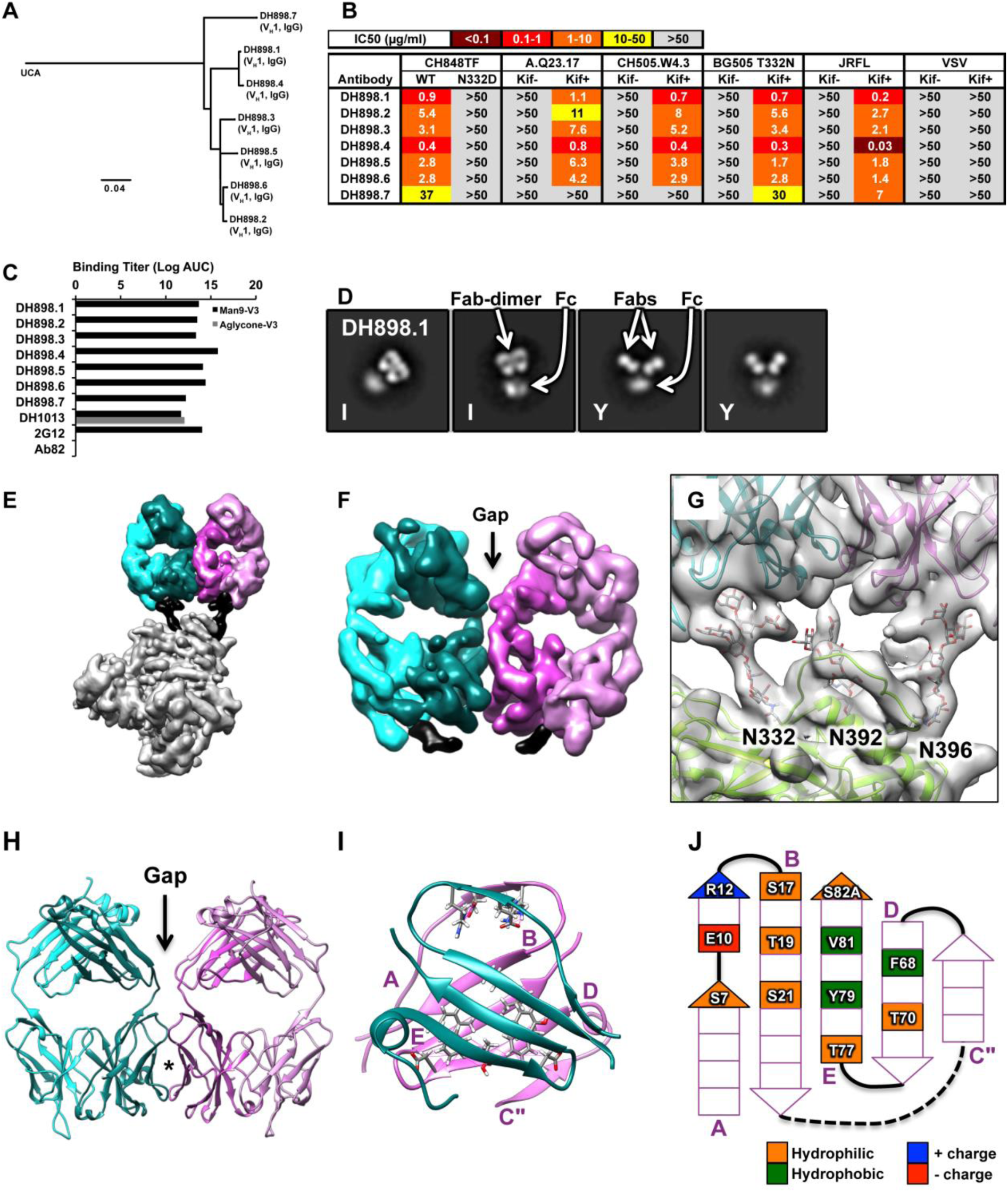
Characteristics of macaque CH848TF SHIV-induced Env glycan-reactive neutralizing antibody B cell lineage, DH898. **(A)** Phylogram and immunogenetics of DH898 neutralizing antibodies. **(B)** Neutralization of DH898 mAbs against autologous tier 2 HIV-1 CH848TF strains bearing wild-type and mutant Envs in TZM-bl cells. Data shown are from a representative assay and neutralization titers are IC50 in µg/ml. **(C)** ELISA binding titers (Log Area Under the Curve, AUC) of DH898 mAbs to synthetic Man_9_-V3 glycopeptide (black bars) and non-glycosylated aglycone V3 peptide (gray bars) (*33*). Control antibodies tested included peptide-reactive DH1013, Env glycan-reactive 2G12, and negative-control anti-influenza CH65 mAbs. Data shown are representative of four assays. **(D)** NSEM analysis of DH898 IgG showing a mixture of I-shaped (I) and Y-shaped (Y) antibodies. **(E)** Segmented cryo-EM map of DH898.1 Fab domains bound to stabilized CH848 10.17 SOSIP trimer (*32*) showing an Env trimer with a single Fab-dimer bound to the trimer. Colors are as in Figure 1. **(F)** Zoomed-in view of the Fab-dimer showing a gap between the two Fabs. **(G)** A fitted model showing the Fab-dimer binding to glycans N332, N392 and N396 at the V3 glycan bnAb epitope. **(H)** Atomic model of the Fab-dimer showing a gap between the Fabs (arrow) and an interface between the Fv domains (star). **(I)** The beta-sandwich view of the interface demonstrating that the beta-strands of the two heavy chains are angled relative to one another. **(J)** Schematic diagram of the Fab-dimer interface showing a patch of three hydrophobic or aromatic residues (green) surrounded by small hydrophilic residues (orange) and a complementary pair of charged residues (blue +, red -).

NSEM analysis of the DH898 lineage antibodies also showed a mixture of Y-shaped and I-shaped antibodies (**Figure 2D and S8),** thus demonstrating that DH898 antibodies are FDG antibodies. To visualize the antibody epitope and Fab-dimer interface, we collected two cryo-EM datasets of a CH848 SOSIP trimer bound with recombinantly expressed Fab, either DH898.1 or DH898.4 (**Figures 2E-2G and S9-S11; Table S5-S6; Movie S1 and S2**). Within both cryo-EM datasets, we identified three populations yielding three independent reconstructions (six total) of the unliganded trimer and two distinct complexes of Fab-dimer-bound trimer. There were no distinguishable differences between the corresponding reconstructions from DH898.1 and DH898.4, within the limits of resolution, ranging from 3.8 to 7.7 Å (**Figures S11A-S11J**). One population of Fab-dimer bound glycans N332, N392 and N396 near the base of the Env V3 loop (**Figures 2G and S11J; Movie S1**), consistent with the N332-dependence of the CH848TF virus neutralization (**Figure 2B**). The second Fab-dimer population bound glycans N386, N362 and N276 surrounding the Env CD4-binding site (**Figure S11I; Movie S2**). In all four DH898.1 and DH898.4 Fab-dimer complexes, the two Fabs were not V_H_ domain-swapped (**Figure 2F and S11G-S11H**). Similar to DH851 and 2G12, the DH898 Fabs dimerized via their V_H_ domains (**Figure 2H**) such that the beta-strands had aromatic and hydrophobic residues in the interface. The DH898 Fab-dimer interface also involved a glutamate-arginine pair, residues E10 and R12, that formed salt bridges between the two V_H_ chains (**Figure 2I-2J**), analogous to a D72 and R57 aspartate-arginine pair seen in 2G12 (**Figure S6D-S6E**). Mutations predicted to disrupt the Fab-dimer interfaces for DH898 as well as DH851 mAbs yielded a lower proportion of I-shaped antibodies and a concomitant loss of antibody binding activity (**Figure S12**). Conversely, mutations predicted to strengthen the Fab-dimer interface resulted in a higher proportion of I-shaped antibodies without any enhancement of biological activity (**Figure S12**), suggesting that the I-shaped conformation is necessary but not sufficient for biological function. These data demonstrated that DH898 antibodies are candidate FDG bnAb precursors.

To determine the reproducibility of SHIV-induced bnAbs, we sought FDG bnAbs from a second SHIV-infected macaque infected with SHIV BG505, and found a 42-member IgG FDG B cell clonal lineage (DH1003). Recombinant IgGs of three representative DH1003 antibodies bound to Man_9_-V3 glycopeptide and *Candida albicans*, neutralized three of nine members of the global panel of HIV-1 isolates (*35*), and showed predominant I-shaped antibodies by NSEM (**Figures S13-S14**).

### Macaque FDG antibodies prior to SHIV infection

In peripheral blood of macaque RM6163 collected before SHIV infection, we found class-switched and mutated DH898 and DH851 clonally-related V_H_ by Illumina next generation sequencing (NGS) (**Table S7**), suggesting that host or environmental antigens may have initiated FDG antibody lineages. From NGS VDJ transcripts, DH898 and DH851 FDG V_H_ clonal lineage members were found at a relative frequency of 0.02% of 3.5 million V_H_1 reads and 0.01% of 2.1 million V_H_2 reads, respectively, and were predominantly IgM sequences. To determine the binding profile of week 0 clonally-related DH851 VH sequences, recombinant antibodies bearing pre-infection (week 0) DH851 IgG (DH851.5) or IgM (DH851.6 and DH851.8) clonally-related V_H_DJ_H_ were paired with DH851 Vλ genes; pre-infection DH851 mAbs weakly bound yeast antigens and HIV-1 CH848 SOSIP trimers, but did not bind Man_9_-V3 glycopeptide (**Figure S2C, and S2E-S2F**). However, post-SHIV-infection DH851 mAbs robustly bound yeast antigens, Man_9_-V3 glycopeptide and HIV-1 Env SOSIP trimers (**Figure S2D-S2F**). Pre-infection DH851 lineage antibodies had Y-shaped conformations, whereas post-SHIV infection DH851 lineage antibodies showed I-shaped Fab dimerization (**Figure S3**). Thus, FDG precursors existed as mutated or class-switched non-Fab-dimerized antibodies prior to SHIV infection whereby lineage stimulation potentially occurred by glycan-bearing host or environmental antigens.

### FDG antibodies induced by macaque Env vaccination

Next we immunized four rhesus macaques intramuscularly (IM) with monomeric Man_9_-V3 glycopeptide (*33*) followed by a single dose of a Man_9_-V3 multimer comprising multiple copies of the Man_9_-V3 glycopeptide scaffolded on a N-(2-Hydroxypropyl) methacrylamide (HPMA)-linear copolymer (**Figure 3A**). After Man_9_-V3 multimer immunization, plasma antibodies of all four macaques bound Man_9_-V3 glycopeptide (**Figure 3B**). Tetramerized fluorophore-labeled [Kif]- treated HIV-1 JRFL gp140 was used to isolate memory B cells from two monkeys following either Man_9_-V3 monomer or multimer immunizations. We isolated twenty-two antibodies, fifteen of which were IgM isotype, and the majority used either macaque V_H_3 or V_H_4 (20/22) gene segments with HCDR3s ranging from 13-17 amino acids (**Figure 3C**). All antibodies bound Man_9_-V3 glycopeptide, 18/22 bound [Kif]-JRFL gp140, and 2/22 weakly bound JRFL gp140 with heterogeneous glycans (**Figure 3D**). From one of the two macaques (RM5996), we isolated a FDG B cell clonal lineage with four members (DH717) following either immunization with Man9- V3 glycopeptide monomer (DH717.1 IgG) or multimer (DH717.2-DH717.4 IgMs) (**Figure 3E; Table S2**), as well as three additional candidate FDG antibody lineages that were isolated following the Man_9_-V3 glycopeptide monomer (DH715 lineage) or multimer (DH780 and DH781 lineages) immunizations (**Table S2**). Vaccine-induced mAbs bound Man9-V3 glycopeptide but not the aglycone-V3 peptide, and bound HIV-1 CH848 SOSIP trimers as well as *Candida albicans* or *Cryptococcus neoformans* yeast glycans (**Figure S15A-S15C**). These vaccine-induced mAbs also demonstrated binding to high mannose glycans, including Man_9_GlcNAc_2_ (**Figure S15D**). DH717 mAbs showed broad neutralization of nine [Kif]-treated HIV-1 strains, but did not neutralize the corresponding wild-type HIV-1 (**Figure 3F**).

**Figure 3.**
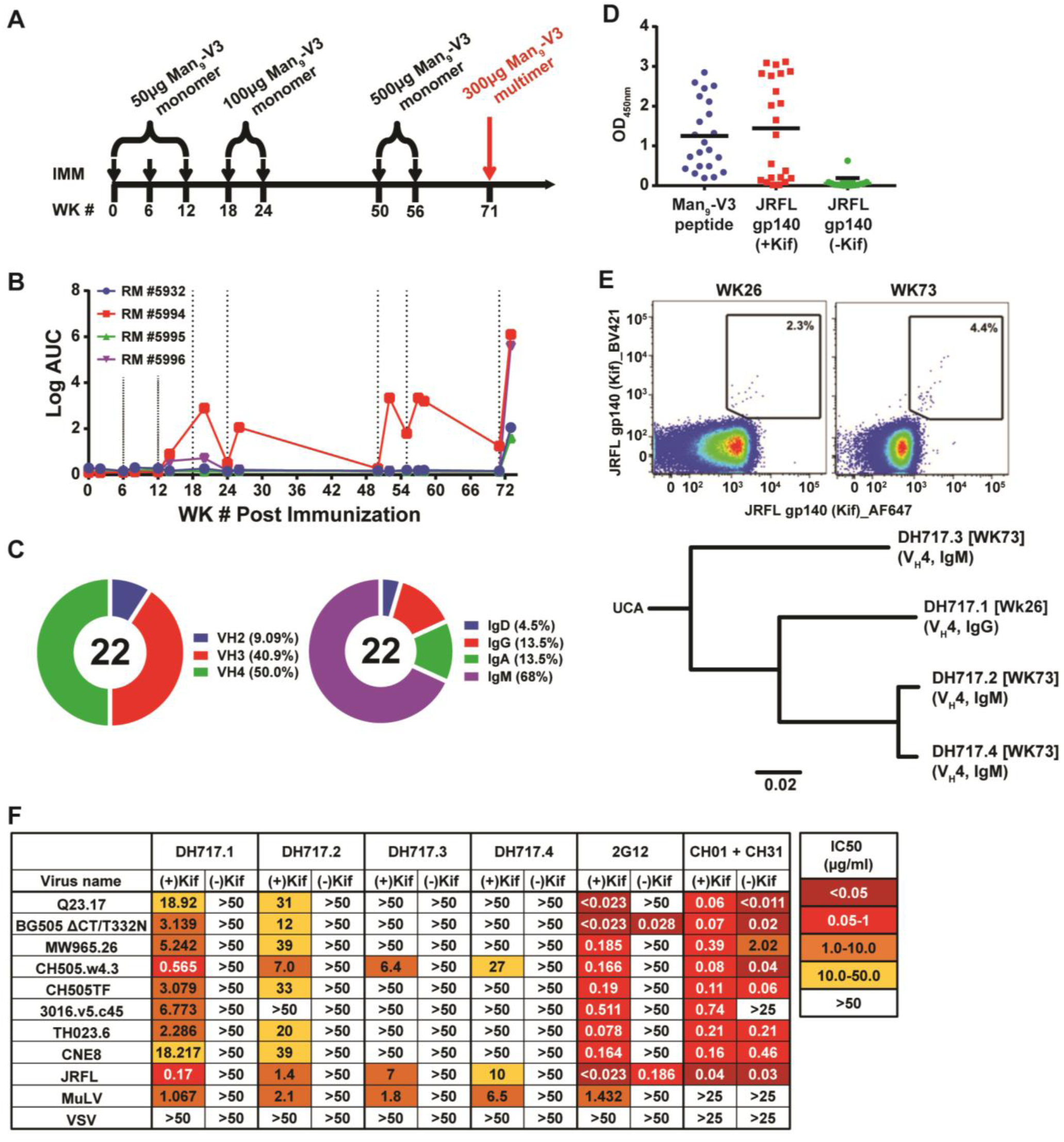
Characteristics of macaque HIV-1 vaccine-induced Env glycan-reactive neutralizing antibody B cell lineage, DH717. **(A)** Dose-escalation of synthetic Man_9_-V3 glycopeptide that mimics a V3 glycan bnAb epitope (*33*) in rhesus macaques (RMs). **(B)** ELISA binding levels (OD_450nm_) of longitudinal plasma IgG from four immunized animals to Man_9_-V3 glycopeptide. Longitudinal plasma from different weeks (WK) post immunizations (IMM) were screened. Data shown are from a single assay, but are representative of multiple ELISAs using these samples. **(C)** Summary of heavy chain gene usage and isotype for glycan-binding antibodies isolated from peripheral blood of RMs. **(D)** ELISA binding levels (OD_450nm_) of transiently-expressed antibodies with Man_9_-V3 glycopeptide, kifunensine [Kif]-treated JRFL gp140 bearing predominantly high mannose glycans, and non-[Kif]-treated JRFL gp140 bearing heterogeneous glycans. Each dot represents a single antibody. Data shown are from a single ELISA. **(E)** Representative plot of Env-reactive memory B cells at weeks 26 and 73 that were individually sorted by flow cytometry, and a phylogram of the isolated DH717 lineage antibodies. **(F)** Neutralization of DH717 lineage mAbs against a multiclade of HIV-1 strains grown in either the presence or absence of [Kif]. Neutralization was tested in TZM-bl cells in a single experiment and titers are reported as IC50 in µg/ml.

To determine if different Env forms can boost mutations in FDG antibodies, we performed NSEM on a macaque Env-reactive neutralizing IgG (DH501) that neutralized [Kif]-treated HIV-1 strains and was induced by a group M consensus Env (*36*), and found it also to have I-shaped antibody populations (**Figures S12 and S16**).

### Structure of DH717 FDG antibodies

NSEM of DH717 antibodies revealed a mixture of I- and Y-shaped antibody populations for DH717.1, DH717.2 and DH717.4, whereas DH717.3 had only Y-shaped antibody populations (**Figure 4A-4D**). The I-shaped structures observed in 2D class averages show one Fab edgewise and one Fab flatwise, suggesting that the two are Fabs turned ∼90° to each other (**Figure 4A**); this arrangement was confirmed by a 3D negative stain reconstruction of the antibody (**Figure S17A**). This arrangement is unlike the parallel, side-by-side Fab-dimers observed in 2G12, DH851 and DH898, suggesting that DH717 Fab-dimerized by a different mechanism. Indeed, all DH717 antibodies lacked the pattern of hydrophobic heavy chain residues seen in the other three Fab-dimerized antibodies (**Figure S17B**). However, the DH717 heavy chain sequences encoded an unpaired cysteine at residue 74, with the exception of DH717.3 which had a serine at this position (**Figure S17B**), suggesting that a disulfide linkage between the heavy chains was the mechanism of Fab-dimerization in the DH717 lineage. For structural analyses of DH717 Fab dimers, recombinantly expressed Fabs were separated by size exclusion chromatography into monomer and dimer fractions (**Figure 4E-4F**). We obtained a 2.6-Å crystal structure of the monomeric DH717.1 Fab complexed with a synthetic Man_9_-V3 glycopeptide (**Figure 4G; Table S8**). The DH717.1 heavy chain dominated interactions with the Man_9_ glycan by using the three HCDR loops to form a binding pocket into which Man_9_ glycan was inserted and no interactions were observed with the peptide itself (**Figures 4G and S17C**), reminiscent of the glycan-only binding mechanism of 2G12 (*22-24, 26, 37*). We also obtained a 3.5 Å crystal structure of unliganded dimeric DH717.1 Fab (**Figures 4H and S17D; Table S8)**. The structure of the DH717.1 Fab dimer consisted of two disulfide-linked Fab dimers in the asymmetric unit, with an inter-Fab disulfide bond between the cysteine 74 residues in the heavy chains of the two Fabs in a dimer (**Figure 4H**). Disrupting this disulfide bond with a cysteine to serine (C74S) mutation in the DH717.1 antibody disrupted the I-shape of DH717.1 (**Figure 4I**), and resulted in reduced or loss of biological activity of DH717.1 (**Figure 4J-4K and S17E**). Thus, a novel disulfide linkage between the heavy chains was indeed the mechanism of Fab-dimerization in the DH717 lineage.

**Figure 4.**
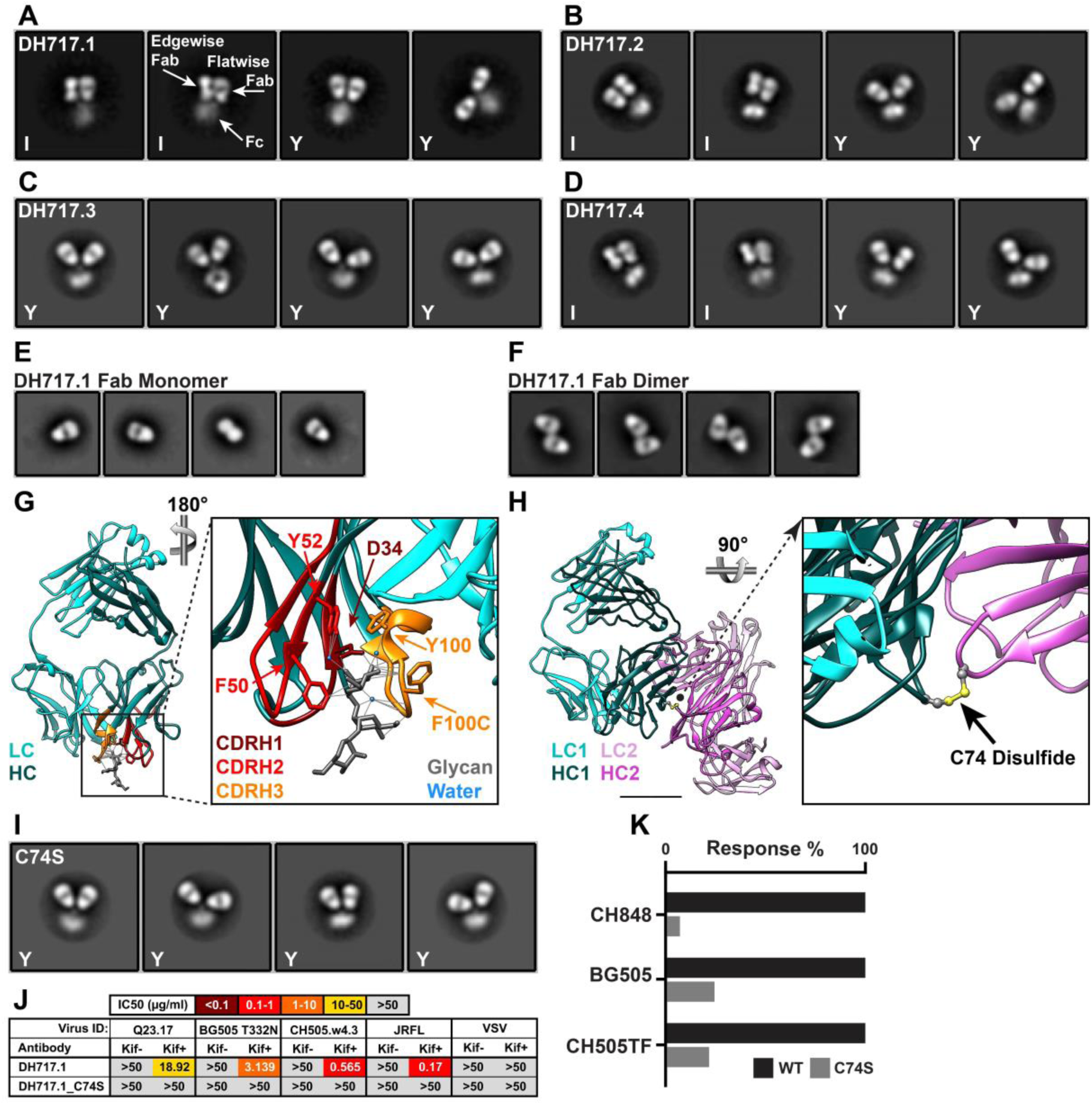
Structural features of DH717 lineage antibodies. NSEM 2D class averages for **(A)** DH717.1 IgG, **(B)** DH717.2 IgG, **(C)** DH717.3 IgG and **(D)** DH717.4 IgG. The I-shaped IgGs are labeled “I” within the 2D class averages. NSEM 2D class averages for DH717.1 monomeric **(E)** and dimeric **(F)** Fabs are also shown. **(G)** Crystal structure of DH717.1 Fab monomer bound to synthetic Man_9_-V3 glycopeptide (*33*), with the antibody heavy chain colored dark teal and light chain colored light teal. The terminal three glycan-moieties of the Man_9_-V3 glycopeptide are represented as sticks and shown bound to a pocket formed by the CDRH1-3 loops (dark red, red and orange) of the DH717.1 Fab. The inset shows a zoomed-in view of the glycans bound to DH717.1. **(H)** Crystal structure of DH717.1 dimer with the heavy chains colored dark teal or pink, and light chains colored light teal or pink. The inset shows a zoomed-in view of the inter-Fab disulfide represented as yellow spheres. **(I)** NSEM 2D-class averages of DH717.1 IgG with cysteine (C) to serine (S) mutation at heavy chain Kabat position 74 (C74S). **(J)** Neutralization of kifunensine [Kif]- treated and non-[Kif]-treated HIV-1 strains in TZM-bl cells by DH717.1 and DH717.1 C76S IgG in a single assay. Neutralization titers were reported as IC50 in µg/ml. **(K)** Binding of stabilized SOSIP trimers (CH848 10.17, BG505 and CH505TF strains) to DH717.1 IgG (dark gray) and DH717.1 C76S IgG (light gray) tested in a single SPR assay. The binding responses for the mutant antibody are reported as a percentage of the wild type antibody binding per SOSIP tested.

Vaccine-induced Env glycan-reactive antibodies in the DH715, DH780 and DH781 lineages showed only Y-shaped antibody populations by NSEM (**Figures S17F-S17J**). They likewise lacked the signature of hydrophobic heavy chain residues seen in 2G12, DH851 and DH898 as well as the crucial cysteine at residue 74 seen in the three Fab-dimerized DH717 antibodies. (**Figure S17K**). These data are consistent with the observation that precursor FDG lineage antibodies such as DH851.4 may have only Y-shaped antibody populations.

### Strategies for boosting mutations in vaccine-induced FDG antibodies

DH715 and DH717 heavy chain gene lineage members were present prior to vaccination, but were mutated IgM B cells (**Table S9**), in agreement with pre-infection DH851 and DH898 heavy chain genes (**Table S7**), thus suggestive of a role for host or environmental antigens in initiating FDG B cell lineages. Importantly, DH715 and DH717 lineages were boosted by Man_9_- V3 glycopeptide (**Figure 5A-5C, and Table S9**). Two recombinant mAbs bearing DH717 clonally-related mutated IgM V_H_DJ_H_ (DH717.5-DH717.6) found at week 0, prior to vaccination and paired with DH717.1 Vλ, demonstrated binding to HIV-1 CH848 SOSIP trimers, Man_9_-V3 glycopeptide, *Candida albicans* yeast antigens, and high mannose glycans (**Figure 5D-5G, and Figure S18**). Additionally, the inferred DH717UCA mAb weakly bound *Candida albicans* glycans, while the mature DH717 lineage antibodies demonstrated affinity maturation after immunization as evidenced by enhanced binding to Man_9_-V3 glycopeptide, to CH848 SOSIP trimers and to *Candida albicans* or *Cryptococcus neoformans* yeast glycans (**Figure 5D-5G**).

**Figure 5.**
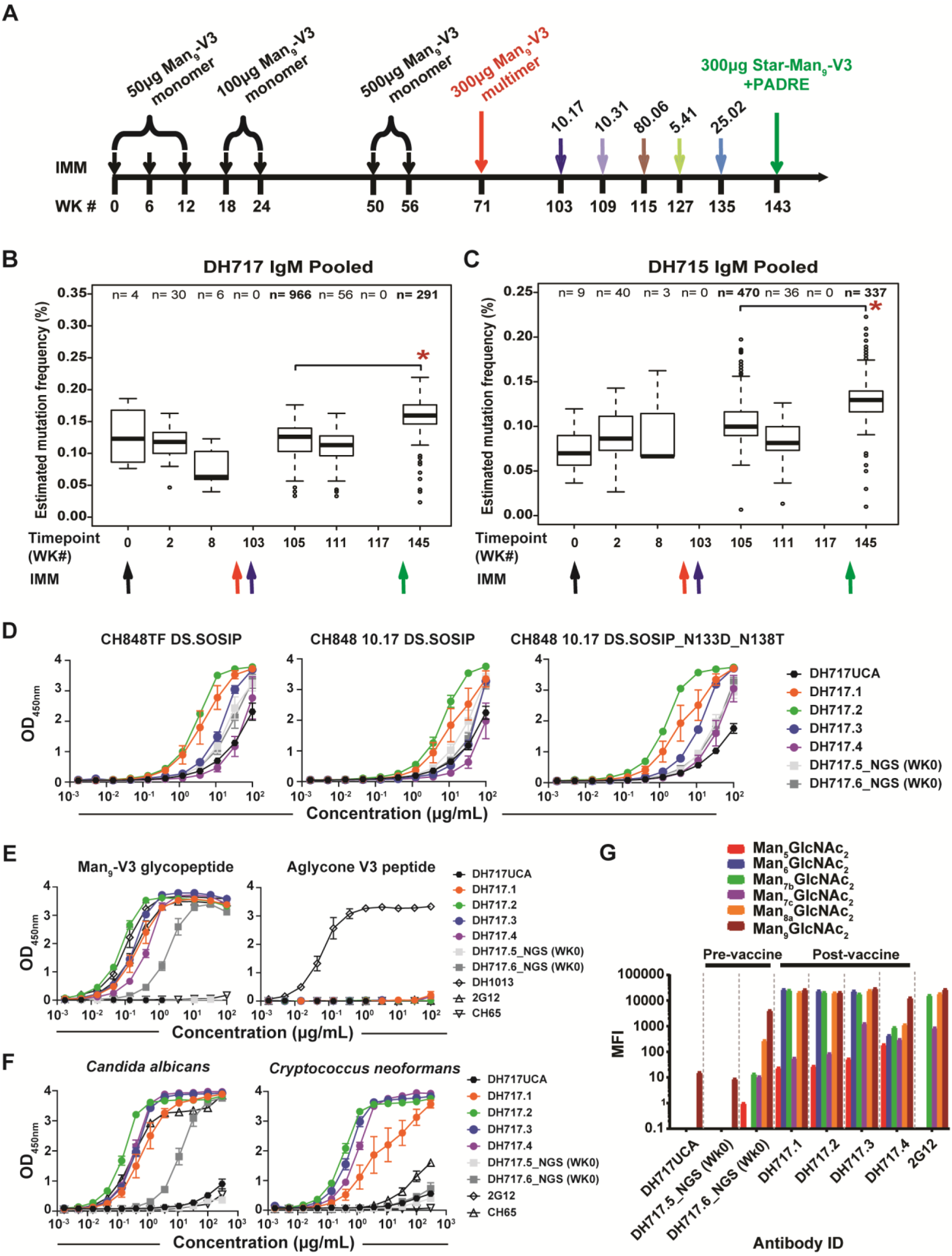
Ability of immunogens to boost the DH717 and DH715 FDG antibodies to undergo affinity maturation. **(A)** Immunization regimens and schedule in rhesus macaques; different forms of synthetic Man_9_-V3 glycopeptides present a mimic of the V3 glycan bnAb epitope (*33*), and SOSIP Env trimers of HIV-1 CH848 strains (10.17, 10.31, 80.06, 5.51 and 25.02) represent sequentially evolved Envs in the CH848 subject who made V3 glycan bnAbs (*20, 32*). **(B-C)** Illumina next generation sequencing (NGS) analysis of vaccine-induced FDG lineages, DH717 and DH715, respectively. Absolute read numbers of DH717 or DH715 lineage members were pooled from two NGS datasets generated from separate aliquots of longitudinal blood-derived RNA at indicated weeks. Statistical significance of an increase in the distribution of estimated mutation frequencies (Cloanalyst software program) at Week 105 compared to Week 145 was determined by a non-parametric Wilcoxon test; asterisks indicate *p* < 0.0001. ELISA binding levels (OD_450nm_) of recombinant DH717 lineage mAbs, including the week 0, computationally inferred UCA, and mature antibodies isolated from single B cells, for reactivity with HIV-1 CH848 Env SOSIP trimers (*32*) **(D)**, synthetic Man_9_-V3 glycopeptide (*33*) **(E)**, and heat-killed *Candida albicans* and *Cryptococcus neoformans* yeast **(F)**. CH65 was used as a negative control mAb, whereas biotinylated trimer-specific mAbs (see Figure S18), and peptide (DH1013) and Env glycan (2G12) specific mAbs were used as positive controls. ELISA data shown are average of 3-5 separate assays where we tested mAbs; error bars represent standard error of the mean. **(G)** Binding of DH717 mAbs to individual glycans in a luminex binding assay, where glycan reactivity was reported as mean fluorescence intensity (MFI) after background subtraction.

Next, we investigated stabilized Env SOSIP trimers of CH848 strains that were isolated from an HIV-1 infected individual that developed a V3-glycan bnAb (*20, 32*) (**Figure 5A**) and Man_9_-V3 arrayed multivalently on a star polymer (“Star-Man_9_-V3”) (*38*) bearing the PADRE T cell helper epitope (*39*) for their ability to boost mutations in FDG antibodies. The number of FDG B cell lineage members were boosted following CH848 10.17 DS.SOSIP immunizations at week 105 (2 weeks post immunization) compared to week 103 (prior to CH848 10.17 DS.SOSIP immunization) (**Figure 5B-5C; Table S9**). DH715 (n=466) and DH717 (n=966) IgM clonal lineage members were found among four million unique V_H_4 gene sequences at week 105 compared to zero lineage members found at week 103. Moreover, DH715 and DH717 IgM lineage members had a higher estimated mutation frequency following immunization with Star- Man_9_-V3 bearing the PADRE T cell helper epitope at week 145, when compared with the distribution of mutation frequencies for DH715 and DH717 lineage members at week 105 that were boosted by CH848 10.17 DS.SOSIP trimer (Wilcoxon test, *p*<0.0001) (**Figure 5B-5C; Table S9**). Thus, these data demonstrated that the CH848 10.17 DS.SOSIP trimer and Man_9_- V3 star polymer are candidate immunogens to boost FDG antibodies.

In an effort to assess vaccine-induced maturation of DH717 lineage for glycan reactivity, we performed SPR affinity measurements of whole IgG and Fab forms of DH717.1 (**Figure S19**). DH717.1 Fab dimer bound HIV-1 CH848 SOSIP trimers, albeit at high micromolar affinity, whereas the DH717.1 Fab monomer did not bind. Moreover, DH717.1 IgG mAb demonstrated nanomolar affinity for binding CH848 SOSIP trimers, thus demonstrating a role for avidity in glycan recognition of FDG antibodies.

### Precursor FDG antibodies in HIV-1-naïve humans

From the B cell repertoire analysis of nine HIV-1 naïve individuals using antigen-specific flow sorting (**Figure 6A**), candidate FDG antibodies were present at an average estimated frequency of 1 per 340,000 B cells; they used a variety of heavy and light chain genes, and were predominantly of the IgM isotype (**Table S2**). One such FDG antibody (DH1005) used an IgG3 mutated V_H_1 gene with a 10 amino acid long HCDR3 (**Table S2**), and was found to bind CH848 DS.SOSIP trimers, and Man_9_-V3 glycopeptide but not aglycone V3 peptide, as well as *Candida albicans* and *Cryptococcus neoformans* yeast (**Figure 6B**). DH1005 neutralized nine heterologous tier 1 and tier 2 HIV-1 strains bearing Man_9_-enriched Envs (**Figure 6C**), but did not neutralize wild-type HIV-1 strains with Envs bearing complex glycoforms (not shown), a similar neutralization profile as other FDG precursors. Moreover, NSEM of DH1005 IgG revealed a mixture of Y-shaped and I-shaped forms of antibodies (**Figure 6D**) with the I-shaped dimer mediating Env glycan interactions (**Figure 6E**). Additional Env glycan-reactive human mAbs (DH1008-DH1010) also had a mixture of I- or Y- shaped antibody populations (**Figure S20**). Thus, these data demonstrate that FDG antibodies are abundant in the B cell repertoire of HIV seronegative humans.

**Figure 6.**
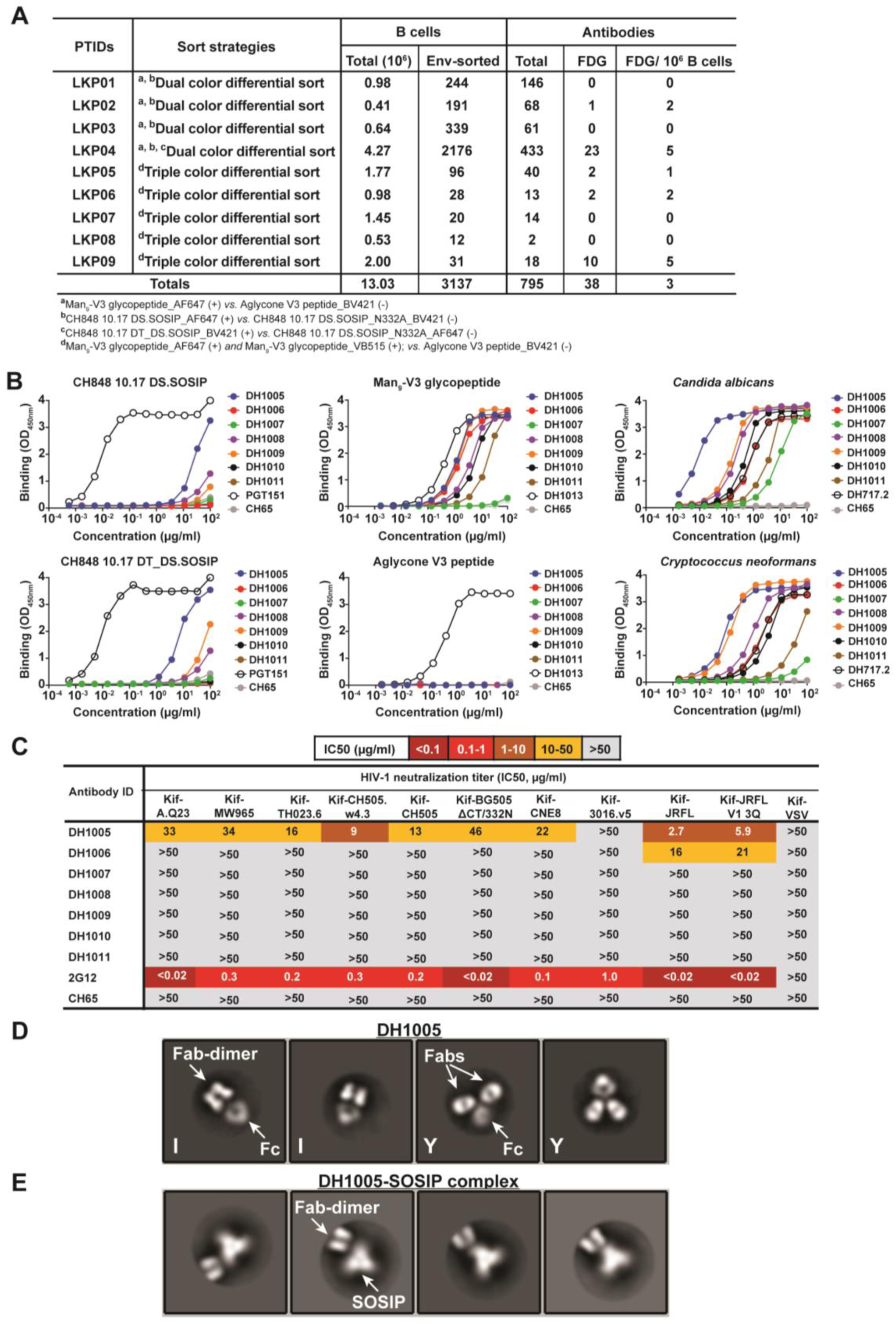
Characteristics of Env glycan-reactive neutralizing antibodies in the B cell repertoire of HIV-1 naïve humans. **(A)** Repertoire of Env glycan-reactive B cells studied in a cohort of nine individuals. B cells were enriched from leukapheresis peripheral blood cells and sorted using fluorophore-labeled Env probes that have been shown to isolate V3 glycan bnAb B cells (*32–33*). Here, we show that these strategies also isolated FDG antibodies. **(B)** ELISA binding levels (OD_450nm_) of recombinant mAbs to CH848 10.17 SOSIP trimers (*32*), synthetic Man_9_-V3 glycopeptide and non-glycosylated aglycone V3 peptide (*33*), and heat-killed *Candida albicans* and *Cryptococcus neoformans* yeast. Data shown are from a representative ELISA. Reference mAbs included peptide-reactive DH1013, Env glycan-reactive DH717.2 and 2G12, trimer-reactive PGT151 and negative control CH65 antibodies. **(C)** Neutralization of mAbs against multiclade kifunensine [Kif]-treated heterologous tier 1 and tier 2 HIV-1 strains bearing Envs with predominantly high mannose glycans. Neutralization titers are representative of two assays in TZM-bl cells, and neutralization titers were reported as IC50 in µg/ml. None of the human FDG mAbs neutralized non-[Kif]-treated HIV-1 strains (not shown). Control mAbs were 2G12 and CH65. **(D)** NSEM 2D class averages of DH1005 antibody; I-shaped antibody conformations denoted by “I” in image. **(E)** 2D class averages of DH1005-SOSIP complex shows Fab-dimer binding to SOSIP. Fc domain is not visible here because it lies outside the circular mask used during class averaging.

Previous reports indicated that the 2G12 bnAb was not polyreactive for binding recombinant human autoantigens (*11, 40*), but a recent study demonstrated that HIV-1 Env bnAbs that bind glycans and the Env peptide backbone also bound host cells from HIV-1 uninfected individuals in a glycan-dependent manner (*41*). Similarly, we found that 12 of 23 (52%) human or macaque FDG antibodies as well as 2G12 bnAb bound HEp2 cells, but were either weakly or non-reactive with recombinant autoantigens (**Figure S21-S22**).

## Discussion

In this study, we demonstrate a new category of prevalent glycan-reactive antibodies with dimerized Fab domains, similar to the glycan-reactive bnAb 2G12 but lacking the unique 2G12 heavy chain V_H_ domain-swapped conformation. The SHIV-induced DH851 antibodies are FDG bnAbs with 26% neutralization breadth of HIV-1 isolates, while the SHIV-induced DH898 and vaccine-induced DH717 antibodies have characteristics of FDG bnAb precursors.

The phenotype of FDG bnAb precursors includes non-domain swapped Fab dimerization and/or reactivity with yeast high mannose glycans, reactivity with HIV-1 Env, and neutralization of HIV-1 with high mannose glycans but not wild-type HIV-1. One DH851 IgM clonal lineage member, DH851.4, did not have bnAb activity suggestive of a DH851-like bnAb precursor. Interestingly, both the FDG bnAb lineages, DH851 and DH1003 were class-switched and IgG. Moreover, DH851 clonal lineage FDG precursors were isolated prior to SHIV infection that had the same characteristics of vaccine-induced FDG antibodies, which is also strong evidence for the phenotype of FDG precursors as yeast glycan-binding, Fab-dimerized antibodies. That IgM DH851.4 bound HIV-1 Env SOSIP trimers at relatively higher affinity than week 0 DH851 antibodies suggest that DH851.4 was a participant in the SHIV response, but we cannot rule out that DH851.4 could have been selected by an environmental antigenic challenge.

FDG antibodies isolated prior to SHIV infection or HIV vaccination were mainly mutated IgM, thus raising the hypothesis that IgM memory B cells are the source of FDG antibodies and provide a possible explanation for their common occurrence in humans and macaques. FDG and 2G12 bnAbs have short HCDR3 segments, and target only glycans and are more common than HIV-1 bnAbs with long HCDR3s (*42, 43*).

Additionally, our data demonstrated that approximately half of the FDG antibodies and 2G12 had polyreactivity profiles. Despite glycan polyreactivity by FDG and 2G12 bnAbs, differential affinity of FDG for glycans presented in a non-self arrangement on pathogens compared to host proteins may be a discriminating factor for antibody maturation and neutralization potency (*28*), thus supporting the hypothesis for affinity maturation of FDG B cell lineages via HIV Env vaccinations. Moreover, we demonstrated a role for avidity in modulating maturation of glycan recognition by FDG antibodies.

RM6163 from whom the FDG bnAb DH851 clonal lineage was isolated had weak heterologous neutralizing activity that mapped to N332 and the GDIR motif at the base of the V3 loop (Shaw, GM, unpublished, manuscript submitted). Thus, it is likely that both the V3-glycan activity and the FDG bnAb activity were minor components of the B cell repertoire of this macaque.

HIV-1 Env glycan bnAb with a domain-swapped conformation has only been reported for 2G12, but here we show two examples of I-shaped Fab-dimerized antibodies without domain-swapped conformation (DH851, DH1003) that can also bind the Env glycan shield and neutralize wild type HIV-1. Mouse B cell lines bearing mature 2G12 have been reported to express domain-swapped forms of the BCR (*44*), but it remains unknown if the membrane-bound BCR of FDG bnAbs in this study exist in an I-shaped conformation and whether this conformation on the B cells is analogous to BCR cross-linking that results in B cell activation.

It is interesting that the FDG bnAbs in this report were only isolated from SHIV-infected macaques. We have previously shown that those HIV-1-infected individuals that make bnAbs have multiple immune-regulatory defects including defective T regulatory cells and dysfunctional NK cells that regulate B cell development, thus creating a permissive germinal center environment for bnAb development (*45–47*). It will be of interest to determine if SHIV-infected macaques that make bnAbs have similar immunoregulatory abnormalities.

Finally, that FDG antibodies can bind to multiple glycan clusters and use multiple V_H_ and Vκ or V_λ_ pairs, combined with FDG precursor frequency make this new category of glycan-reactive antibodies an attractive target for HIV-1 bnAb vaccine development. V3-glycopeptide monomers can induce FDG precursor expansion, and Man_9_-V3 multimers and SOSIP trimers can boost mutations in FDG antibodies. It will be important to produce these and related immunogens for testing in human phase I clinical trials for their ability to initiate and boost FDG clonal B cell lineages to bnAb status.

## Acknowledgements

We thank Kelly Soderberg, Ashley Trama, Hilary Bouton-Verville, Jordan Cocchiaro and Amanda Eaton for program management support, Lawrence Armand for generating Env hooks for sorts, Caroline Jones and Yousef Abuahmad as sort operators, William Faison and Sravani Venkatayogi for computational support, and Charles Giamberardino and Jennifer Tenor for generating heat-killed yeast antigens. The following individuals have provided technical assistance: Jordan Sprenz, Li Jing, Andy (Shi) Huang, Shi-Mao Xia, Melissa Cooper, Kara Anasti, Maggie Barr, Callie, Vivian, Giovanna Hernandez, Esther Lee, Aja Sanzone and Paige Rawls. Cryo-EM data were collected at the Shared Materials Instrumentation Facility at Duke University. The adjuvant GLA-SE was provided by Infectious Disease Research Institute, Seattle, WA 98102, USA; Christopher Fox and Steve Reed. Statistical support was provided by the DHVI Biostatistics team; Wes Rountree and Yunfei Wang. This work was supported by UM1 AI100645, the Duke Center for HIV/AIDS Vaccine Immunology-Immunogen Discovery, and UM1 AI44371, the Duke Consortium for HIV/AIDS Vaccine Development (CHAVD), Division of AIDS, NIAID, NIH; Duke University Center for AIDS Research (CFAR); Translating Duke Health Initiative (P.A.), NIH/NIAID Contract #HHSN272201800004C (C.C.L), NIH NIAID extramural project grants P01AI131251 (G.M.S, G.K. and W.B.W), 1R01-AI128832 (G.K.), R01 AI145687 (P.A.) and R01AI120801 (K.O.S.). R.R.M. was supported by a Medical Scientist Training Program (MSTP) training grant (T32GM007171) and by the Ruth L. Kirschstein National Research Service Award F30-AI22982-0 from NIAID. Use of the Advanced Photon Source was supported by the U. S. Department of Energy, Office of Science, Office of Basic Energy Sciences, under Contract No. W-31-109-Eng-38. Support was also provided in part by the Intramural Research Program of the Vaccine Research Center, NIAID, NIH (G.S.J and P.D.K). D.F was supported by the Mathilde Krim Fellowship in Basic Biomedical Research (109502-61-RKVA) from amfAR and Swarthmore College Startup and Faculty Research Funds. This study utilized the computational resources offered by Duke Research Computing (http://rc.duke.edu). We thank M. DeLong, C. Kneifel, M. Newton, V. Orlikowski, T. Milledge, and D. Lane from the Duke Office of Information Technology and Research Computing for providing assistance with the computing environment.

## Author contributions

B.F.H. conceived and designed the study, evaluated all data, and wrote the paper. P.A. designed the structural studies, analyzed the data, co-wrote, and edited the paper. W.B.W. and R.R.M. designed antibody isolation and characterization studies, analyzed data, co-wrote, and edited the paper. R.J.E. performed structural analyses, analyzed data, co-wrote and edited the paper. D.F., N.I.N., A.H, M. Borgnia, Y.Z., A.B., K. Manne, K.M., M.L., V.S., R.H., N.B. and G.S-J performed structural characterization studies ranging from study design to data collection and analyses. J.P. produced and quality controlled fungal antigens. M.A.M. designed flow cytometry sort experiments. T.B., T.B.K., K.W. generated or analyzed antibody gene sequences. S.M.A., K.J., R.J.P., and A.F. characterized antibody binding profile and kinetics. K.O.S. generated recombinant Envs and mAbs, and designed glycan binding assays. D.C.M., C.C.L, M. Bonsignori and M.S. carried out neutralization assays and data analysis. S.S. performed NHP immunizations and care. B.A., W.E.W., J.F., G.M.L., R.L. and R.S. designed or constructed Man_9_-V3 glycopeptides for NHP immunizations. P.D.K. contributed to structural studies. G.K. contributed to B cell lineage evolution studies. G.M.S. and H.L. designed macaque SHIV infection studies.

## Materials and Methods

### Synthesis of Man_9_-V3 glycopeptide constructs

The Man_9_-V3 monomer comprising a 30 amino acid V3-glcocopeptide with high-mannose (Man_9_GlcNAc_2_) glycans at position 301 and 332 based on the clade B JRFL mini-V3 construct (*14*) was synthesized as previously described (*33*). Man_9_-V3 multimer was synthesized by reacting multiple copies of Man_9_-V3 monomer with an amine-reactive HPMA-based co-polymer that was prepared as previously described (*48*). The concept, design and synthesis of Man_9_ glycopeptides multimerized on star polymers (Star-Man_9_-V3) with the universal PADRE T helper epitope (*39*) has been described (*38*). In this study, Man_9_-V3 multimer and Star-Man_9_-V3 were used for immunization of rhesus macaques.

### Expression of recombinant HIV-1 Env proteins

Production of recombinant HIV-1 Envs, including gp120 monomers, and gp140 and stabilized SOSIP trimers was previously described (*32, 36, 38, 49*). Recombinant Envs were used for binding assays and NHP immunizations. The name of SOSIP trimers were abbreviated in the text. Below is the complete nomenclature of the transmitted-founder (TF) and 10.17 versions of CH848 SOSIP trimers: CH848.3.TFchim.6R.SOSIP664v4.1;

CH848.3.D0949.10.17chim.6R.DS.SOSIP664;

CH848.3.D0949.10.17chim.6R.DS.SOSIP664_N301A;

CH848.3.D0949.10.17chim.6R.DS.SOSIP664_ N301AN332A;

CH848.3.D0949.10.17chim.6R.SOSIP664v4.1_N133DN138T;

CH848.3.D0949.10.17chim.6R.DS.SOSIP664_N133DN138T; and

CH848.3.D0949.10.17chim.6R.DS.SOSIP664v4.1degly4.

For SOSIP trimer immunizations, the following CH8484 strains were generated as SOSIP trimers; CH848 10.17., CH848 10.31, CH848 80.06, CH848 5.41 and CH848 25.02. These SOSIP trimers constitute a sequential 5-valent series of CH848 Env stabilized trimers that had evolved during human CH848 infection and were predicted to be involved in the induction of V3-glycan bnAbs (*20*).

In instances where high mannose glycosylation was desired, Kifunensine (Sigma-Aldrich) was dissolved in phosphate-buffered saline (PBS) and added once to the cell culture media to a final concentration of 25 μM. The cells were cultured for 5 days and on the fifth day the cell culture media was cleared of cells by centrifugation and filtered with 0.8 um filter. The cell culture was concentrated with a Vivaflow 50 (Sartorius) with a 10 kd MWCO. The concentrated cell culture supernatant was rotated with lectin beads (Vistar Labs) overnight at 4 °C. The beads were pelleted by centrifugation the next day and resuspended in MES wash buffer. The lectin beads were washed twice and the protein was eluted with methyl-α-pyranoside. The protein was buffer exchanged into PBS and stored at –80C.

### NHP immunization studies

Immunization of Indian origin rhesus macaques and blood draws were performed at (Bioqual Inc., Rockville, MD). Rhesus macaques were immunized intramuscularly with either 50 μg, 100 μg, or 500 μg of Man_9_-V3 monomeric glycopeptide formulated in GLA-SE adjuvant, followed by GLA-SE adjuvanted (∼6-mer) Man_9_-V3 N-(2-Hydroxypropyl) methacrylamide (HPMA) linear copolymer. Macaques were also immunized with CH848 SOSIP trimers in Poly ICLC adjuvant followed by a single boost with Star-Man_9_-V3 in GLA-SE adjuvant. Blood samples were collected two weeks post-immunization. All rhesus macaques were maintained in accordance with the Association for Assessment and Accreditation of Laboratory Animals. Research was conducted in compliance with the Animal Welfare Act and other federal statutes and regulations relating to animals and experiments involving animals and adheres to principles stated in the Guide for the Care and Use of Laboratory Animals, NRC Publication, 2011 edition.

### Isolation of antibodies from single B cells

Antigen-specific memory B cells were isolated from rhesus macaques or humans using fluorophore-labeled antigens as hooks in various sort strategies outlined below. Antigens used for sorts were conjugated with Alexa Fluor 647 (AF647), Brilliant Violet 421 (BV421), or VioBright 515 (VB515) fluorophores. With the exception of peptides, immune cells were concurrently incubated with a cocktail of staining antibodies and fluorophore-labeled antigens. Fluorophore-labeled peptides required an incubation with immune cells prior to addition of the staining antibody cocktail to improve binding efficiency. Fluorophore-labeled peptides were used at a ratio of 2.5µg per 10 million cells, recombinant gp120 and gp140 proteins at 10µg per 10 million cells, and SOSIP trimers at 90pmol per 10 million cells, to stain antigen-specific memory B cells. All incubations took place in approximately 1mL of buffer per 10 mL cells. Flow cytometry sorts were performed using a BD FACSAria II (BD Biosciences, San Jose, CA), and the data were analyzed using FlowJo (Treestar, Ashland, OR).

Single cell Env-reactive antibodies were CD3 (BD #552852; 5µl per test)/ CD14 (BioLegend #301832; 2.5µl per test)/ CD16 (BD #557744; 5µl per test), and/or surface IgD (Southern Biotech #2030-09; 1µl per test) negative. Macaque B cells were also CD27+/- (BioLegend #302816; 5µl per test) and CD20 (BioLegend #302336; 1µl per test) positive, while human B cells were CD38+/- (Beckman Coulter #6699531; 1µl per test) and CD19 (BD #557791; 1µl per test) positive. Single sorted B wells were obtained from rhesus macaques and humans using previously described flow cytometry-based sorts (*33, 50, 51*). Cells were individually sorted into 96 well plates containing lysis buffer and immediately stored at −80, as previously described (*52*). Rhesus macaque and human V_H_DJ_H_ and V_L_J_L_ segments were isolated by single-cell PCR approaches that were previously described (*33, 52*). Antibody sequences were analyzed using a custom-built bioinformatics pipeline for base-calling, contig assembly, quality trimming, immunogenetic annotation with Cloanalyst (https://www.bu.edu/computationalimmunology/research/software/), VDJ sequence quality filtering, functionality assessment, and isotyping.

In vaccinated rhesus macaques, Env-reactive B cells were sorted using Man_9_-enriched HIV-1 JRFL gp140 protein (Kifunensine-treated Env) conjugated to both AF647 and BV421 fluorophores. In SHIV CH848TF infected macaque, several flow cytometry sort strategies were used to investigate the repertoire of envelope (Env)-reactive B cells, including candidate V3- glycan reactive antibodies – LN wk52 [Man_9_-V3 peptide (+) and Aglycone V3 peptide (-); CH848TF gp120 (+) and CH848TF gp120 N332A (-); and CH848TF SOSIPv4.1 (+)]; PBMCs wk52 [CH848TF gp120 (+) and CH848TF gp120 N332A (-)]; and PBMCs wk104 [Man_9_-V3 peptide (+) and Aglycone V3 peptide (-);CH848TF SOSIPv4.1 (+); CH848.d949.10.17 DS.SOSIP (+) and CH848.d949.10.17 DS.SOSIP N332A (-); and CH848.d949.10.17 DS.SOSIP (+) and CH848TF SOSIPv4.1 (-)].

In HIV-1 naïve humans, Env-reactive B cells were sorted using the following differential sort strategies: (#1) Man_9_-V3 glycopeptide (+) vs Aglycone V3 peptide (-) in a dual or triple color sort strategy; (#2) CH848 10.17 DS.SOSIP (+) vs CH848 10.17 DS.SOSIP_N332A (-); and (#3) CH848 10.17 DS.SOSIP_N133DN138T (+) vs CH848 10.17 DS.SOSIP_N133DN138T_N332A (-). In three individuals (LPK01-LPK03), we sorted 20M PBMCs using strategy #1 (dual color differential sort) or strategy #2 (total for 2 sorts = 40 million PBMCs). For dual color differential sort strategy #1, we sorted B cells that were reactive with Man_9_-V3 glycopeptide conjugated to one fluorophore, but not the aglycone V3 peptide conjugated to a different fluorophore. In one individual (LPK04), we studied B cells enriched from 100 million leukapheresis PBMCs per experiment using sort strategy #3 (total for 2 sorts = 200 million PBMCs). In five individuals (LKP05-LKP09), we studied B cells enriched from 100 million leukapheresis PBMCs using sort strategy #1 (triple color differential sort); here we used a more stringent approach to sort B cells that were reactive with Man_9_-V3 glycopeptide conjugated to two different fluorophores, but was not reactive with aglycone V3 peptide conjugated to a third fluorophore. For subjects LKP04-LKP09, we performed negative B cell enrichment from 100M PBMCs in each experiment using a commercially available B cell enrichment kit according to manufacturer’s protocol (StemCell Technologies; Vancouver, BC, Canada).

### Expression of antibodies

Recombinant monoclonal antibodies (mAbs) were first generated in human embryonic kidney epithelial cell lines (HEK 293T, ATCC, Manassas, VA; catalog #CRL3216) in small scale transfections using linear cassettes encoding antibody heavy and light chain genes that were PCR amplified from single B cells (*52*). For generation of larger quantities commercially-obtained (GeneScript, Piscataway, NJ) plasmids with antibody heavy and light chain genes were used to transfect suspension Expi 293i cells using ExpiFectamine 293 transfection reagents (Thermo Fisher Scientific) as described (*20*). Purified recombinant mAbs were dialyzed against PBS, analyzed, and stored at 4 °C. All recombinant mAbs were expressed from plasmids encoding a human or macaque IgG constant region.

### Expression of DH717 Fabs for structural studies

To purify DH717.1 Fab monomer for crystallization studies, the heavy- and light-chain variable and constant domains were cloned into a modified pVRC-8400 expression vector using NheI and NotI restriction sites and the tissue plasminogen activation signal sequence. The C-terminus of the heavy-chain constructs contained a non-cleavable 6x-histidine tag. Fabs were expressed using transient transfection of HEK 293T cells using linear polyethylenimine (PEI) following manufacturer’s protocols. After 5 days of expression, supernatants were clarified by centrifugation. His-tagged Fabs were loaded onto Ni-NTA superflow resin (Qiagen) preequilibrated with Buffer A (10 mM Tris, pH 7.5, 100 mM NaCl), washed with Buffer A + mM imidazole, and eluted with Buffer A + 350 mM imidazole, Fabs were then purified by gel filtration chromatography in Buffer A using a superdex 200 analytical column. The fractions corresponding to monomer were pooled and concentrated.

### ELISA binding of antibodies

Envelope binding to recombinant mAbs, and plasma or sera were tested in ELISA as previously described (*32, 53*). Binding was assessed using synthetic peptides, recombinant HIV-1 Env gp120, gp140, and stabilized chimeric SOSIP trimers, and heat-killed yeast antigens. ELISA protocols had modifications that supported binding to different antigens. In general, antigens were directly coated to Nunc-absorb (ThermoFisher) plates overnight at 4°C or captured using streptavidin, AbC-mAb (AVIDITY, Colorado, USA) or PGT151 mAb that were coated to Nunc-absorp plates overnight at 4°C. Unbound proteins were washed away and the plates were blocked with either goat serum-based (SuperBlock) (*52*) or non-serum-based (*54*) blocking media for 1 hour. Serial dilution of serum and mAbs were added to the plate for 60 and 90 minutes, respectively. Binding antibodies were detected with specie-specific HRP-labeled anti-IgG Fc antibodies using 20µl per reaction with 1 hour incubation at room temperature. HRP detection was subsequently quantified with 3,3′,5,5′-tetramethylbenzidine (TMB). ELISA binding levels were measured at an optical density of 450 nm (OD_450nm_) and binding titers were analyzed as area-under-curve of the log-transformed concentrations (Log AUC).

Binding to biotin-Man_9_-V3 and biotin-aglycone-V3 was determined as previously described using streptavidin for capturing each peptide on Nunc-absorb ELISA plates (*20, 33*). In brief, we coated 30ng of streptavidin in 15µl at 2µg/ml per well of a 384-well Nunc-absorb ELISA plate, sealed and incubated overnight at 4°C. Synthetic peptides (10ng) were added to streptavidin in 10µl at 1µg/ml for one hour at room temperature to facilitate peptide capture. Binding to yeast antigens was tested at 1:400 (*Cryptococcus neoformans*) and 1:2000 (*Candida albicans*) dilutions made in sodium bicarbonate buffer; 15µl of each yeast solution was coated per well of a 384-well Nunc-absorb ELISA plate, sealed and incubated overnight at 4°C. Yeast antigen reactivity was performed using ELISA conditions for assessing glycan reactivity on glycopeptides (*33*). HRP-conjugated specie-specific secondary antibodies were used to detect antibody binding to synthetic peptides and yeast antigens - Goat anti-human IgG-HRP, 1:15000 dilution (Jackson ImmunoResearch Laboratories, CAT# 109-035-098); and Mouse anti-monkey IgG-HRP, 1:20000 dilution (Southern Biotech, CAT# 4700-05). The yeast dilutions tested in ELISA were found to be the lowest dilutions at which we observed maximal binding by glycan-reactive antibodies, such as 2G12 in optimization assays. DH1013 isolated from RM5996 was used as a positive control mAb for peptide binding to Man_9_-V3 and Aglycone V3 peptides, and 2G12 or DH501 (*36*) were used as positive control mAbs for binding Man_9_-V3 glycopeptide, but not Aglycone V3 peptide. 2G12 bnAb was also used as a reference mAb for Env glycan and *Candida albicans* yeast antigen binding (*23*).

For ELISAs with SOSIP trimers bearing an Avi-tag, the trimer was captured by the AbC-mAb antibody against C-terminus Avi-tag, while non Avi-tagged SOSIP trimers were captured by PGT151 bnAb expressed in 293 cells as described (*32, 53*). In brief, Avi-tag and PGT151 mAbs were coated at 30ng per well of a 384-well plate in 15µl at 2µg/ml. SOSIP trimers (20ng) were added to Avi-tag or PGT mAbs in 10µl at 2µg/ml for one hour at room temperature to facilitate SOSIP capture. PGT151 mAb was used as a positive control for binding Avi-tagged SOSIP trimers, while influenza-specific antibody CH65 (*55*) was used as a negative control. HRP-conjugated specie-specific secondary antibodies were used to detect binding antibodies (see above). For non-Avi-tagged SOSIPs that were captured by PGT151 mAb, biotinylated mAbs were used as positive (B-2G12 and B-PGT128) and negative (B-CH65) controls - biotinylated-mAbs were tested at 10µg/ml and 3-fold dilutions. This latter approach was used to test only rhesus mAbs and b-mAbs, since the secondary antibodies would not detect human PGT151 mAb used for SOSP capture; Mouse anti-monkey IgG-HRP (see above) and Streptavidin-HRP, 1:30000 dilution (Thermo Scientific, REF# 21130).

Competitive ELISA to assess cross-blocking of recombinant mAbs were previously described (*20, 36, 56*). We biotinylated the antibodies using the following product: BIOTIN-X-NHS, Cayman Chemicals, CAT# 13316. Here, we studied biotinylated 2G12 and DH851 mAbs in competition ELISAs using CH848TF or CH848 10.17 gp120 Envs. Competitive inhibition of biotiniylated-mAbs was measured as a percent of binding in the presence of a competing non-biotinylated mAb relative to binding in the absence of this competing mAb. A successful assay had positive control blocking ≥40%.

### Neutralization assays

Antibody neutralizing activity was assessed in TZM-bl cells as described (*57–59*). MAbs were tested for neutralization against the global panel of HIV-1 references strains (*35*) or 119 heterologous tier 2 isolates (*60*). Kif-treated HIV-1 strains were grown in cells treated with 25µM Kifunensine to facilitate high mannose glycan expression on HIV-1 Envs and Kif-treated HIV-1 strains were tested for neutralization in TZM-bl cells as previously described (*36*). For neutralization assays, a mixture of CH01+CH31 bnAbs is used as a positive control for neutralization of all HIV-1 strains, and murine leukemia virus (MLv) or vesicular stomatitis virus (VSV) were used as negative retrovirus controls. For neutralization assays, a positive for neutralizing antibody activity in a sample is based on the criterion of >3X the observed background against the MLV negative control pseudovirus. Note that DH898 mAbs were tested for neutralization of all tier 2 HIV-1 strains in the global panel, except CNE8. Additionally, we found neutralization titer variabilities in DH851 antibodies across different batches of antibodies, in agreement with the unstable dimerization of Fabs for these antibodies.

### Glycan binding of antibodies Oligomannose bead immunoassay

Recombinant mAbs were tested for binding glycans in an oligomannose bead immunoassay as described (*36*). Custom glycan microspheres were generated with different individual glycans to evaluate mAb binding. Binding of mAb to glycan was determined with A Bio-Plex 200 HTS (Bio-Rad) machine with the Bioplex manager software (Bio-Rad) was used to quantify binding of mAb to glycan, and binding was measured as background-subtracted fluorescence.

### Anti-nuclear antibody (ANA) reactivity

Monoclonal antibody reactivity to nine autoantigens was measured using the AtheNA Multi-Lyte ANA kit (Zeus scientific, Inc, #A21101). Antibodies were 2-fold serially diluted starting at 50 mcg/mL. The kit SOP was followed for the remainder of the assay. Samples were analyzed using AtheNA software. Positive (+) specimens received a score >120, and negative (-) specimens received a score <100. Samples that score 100-120 were considered indeterminate.

### HEp-2 cell staining

Indirect immunofluorescence binding of monoclonal antibodies to human epithelial (HEp-2) cells (Zeus Scientific, Somerville, NJ) was performed per manufacturer’s instructions. Briefly, 20µL of diluted antibodies (50µg/ml, and 25µg/ml) were added to the appropriate wells on the antinuclear antibody (ANA) slide. Slides were incubated for 20 minutes at room temperature in a humid chamber, and then washed with 1X PBS. Goat-anti-rhesus Ig-FITC (Southern Biotech, Birmingam, AL) secondary was added at a concentration of 30µg/ml to each well. The slides were incubated for 20 minutes, then washed twice, dried, fixed with 33% glycerol and cover-slipped. Slides were imaged using an Olympus AX70 microscope with a SpotFlex FX1520 camera. Images were acquired on a 40X objective using the FITC fluorescence channel with different acquisition time as indicated on each image. Positivity was determined by comparing antibodies of interest to positive and negative non-human primate antibody controls, DH1037 and DH570.30 respectively. Staining patterns were identified using the Zeus Scientific pattern guide found on the website and previously reported (*11, 61, 62*).

### Next-generation sequencing and analysis of antibody genes

Illumina MiSeq sequencing of antibody heavy chain VDJ sequences was performed on peripheral blood cells as previously described (*50, 51*). Briefly, for each timepoint, RNA from peripheral cells was divided equally into two separate portions that were used to generate independent cDNA aliquots for sequencing to confirm the absence or presence of antibody sequences of interest. Paired-end sequences were merged using FLASH (*63*), quality filtered (Q score >30 for >95% of sequence) and deduplicated. V, D, and J gene segment, clonal relatedness testing and reconstruction of clonal lineage trees were performed using the Cloanalyst software package (*64*). Immunogenetics information of rhesus and human antibody sequences were assigned by Cloanalyst using Cloanalyst’s default libraries of rhesus and human immunoglobulin genes, respectively (https://www.bu.edu/computationalimmunology/research/software/). B cell clonality was determined based on similar heavy chain rearrangements and CDR3 length as described (*65*).

The unmutated common ancestor (UCA) of antibody lineages were inferred using the Cloanalyst software program with the macaque immunoglobulin gene library. DH851UCA was initially inferred based on the isolation of DH851.1-DH851.3 lineage members. DH851UCA.2 was later inferred using DH851.1-DH851.4. We found that DH851UCA and DH851UCA.2 had one amino acid difference in a framework region of both the heavy and light chain genes; the UCA sequences were deposited in GenBank. Data shown in this study describes the characteristics of DH851UCA.

To exclude the possibility of generating unmutated common ancestor antibodies that do not account for the allelic variability observed in the rhesus macaque model (*66, 67*), we generated a NGS dataset as described above that comprised only of VH1 (DH898 lineage), VH2 (DH851 lineage) and VH4 (DH717 lineage) IgM+ repertoires at baseline or week 0 of the macaques. Here, we sought to find unmutated or least mutated FDG antibody clonal lineage members indicated from macaque baseline PBMCs. Week 0 clonally-related mAbs were generated by pairing the week 0 NGS-derived VH genes with the light chain of the single B cell-derived antibody isolated following infection or vaccination as described (*68*).

### Macaque SHIV CH848TF infection

The generation of chimeric simian-human immunodeficiency virus (SHIV) bearing transmitted-founder (TF) CH848 Envs and infection of rhesus macaques were previously described (*31*). Blood and plasma samples were collected for binding and neutralization assays.

### Preparation of *Candida albicans* and *Cryptococcus neoformans*

A single colony of yeast was inoculated into 250 ml YPD broth and grown for 2 days at 30°C on a shaking incubator set at 225 rpm. The cells were harvested and washed thrice with PBS. The pellets were resuspended in 11 ml PBS and the cell concentration was determined (C. albicans, 3.40×10e9 CFUs/ml and C. neoformans, 1.62×10e9 CFUs/ml, respectively). To heat-kill the yeast, the cells were incubated for 24h at 60°C in a water bath. Three hundred microliters of the heat-killed yeast were plated onto YPD agar, incubated for 7 days to ensure no viable yeast were present.

### Crystallography

The His-tagged DH717.1 Fab monomer was mixed with 3 molar excess of Man_9_-V3 at a final complex concentration of 15 mg/mL for crystallization trials. The complex crystallized over a 100uL reservoir of 100mM Na Citrate, pH 5.0, 30% PEG 4000 in 1-2 days at room temperature. DH717.1 Fab dimer was crystallized over a 60 µl reservoir of 0.1M citric acid pH 3.5, 14% PEG 1,000 in a drop composed of 0.5 µl protein plus 0.5 µl reservoir. The crystal of monomer in complex with glycopeptide was cryoprotected by brief immersion in reservoir solution supplemented with 25% glycerol and the crystal of dimer was cryoprotected with 30% ethylene glycol before being flash frozen in liquid nitrogen. Diffraction data of monomer-glycopeptide complex were collected at NE-CAT beamline 24-ID-E and data of dimer were collected at SER-CAT with an incident beam of 1 Å in wavelength. Data were processed using HKL-2000 (*69*). Matthews analysis of the monomer-glycopeptide complex and dimer data suggested 1 Fab and four Fabs present in the unit cell, respectively (*70*). Molecular replacement calculations for the monomer-glycopeptide complex were carried out with PHASER, using published DH270.5 (Protein Data Bank (PDB) ID 5TTP) as the starting model. The Fab model was separated into its variable and constant domains for molecular replacement. A molecular replacement solution for the dimeric DH717.1 with four Fabs was found using Phaser with search models DH717.1 Fv (above) and the constant region from DH522UCA (*50, 71*). The solution was improved through alternating rounds of manual rebuilding in Coot and reciprocal space refinement in PHENIX (*72, 73*). The crystal structure of the monomeric DH717.1 Fab complex showed well-defined electron density for the three protein-proximal mannose residues, while the rest of the Man_9_ glycan and the V3 peptide were disordered.

DH717.1 IgG was expressed and purified as above. Subsequently IgG samples were digested with papain and further purified to produce Fab fragment. DH717.1 Fab was further purified via size exclusion chromatography, and elution fractions corresponding to approximately 50 kD (the predominant peak in the elution profile) were pooled and concentrated to 10 mg/ml in 0.1M Hepes buffer with 0.15M NaCl. DH717.1 Fab was crystallized over a 60 ul reservoir of 0.1 M citric acid pH 3.5, 14% PEG 1,000 in a drop composed of 0.5 ul protein plus 0.5 ul reservoir. The crystal was cryoprotected by brief immersion in reservoir solution supplemented with 30% ethylene glycol then flash frozen in liquid nitrogen. Diffraction data were collected at SER-CAT with an incident beam of 1 Å in wavelength. Data were processed using HKL-2000 (*69*). Matthews analysis of the data suggested four Fabs present in the unit cell (*70*). A molecular replacement solution with four Fabs was found using Phaser with search models a preliminary structure of DH717.1 Fv and the constant region from DH522UCA (*71*). The solution was improved through alternating rounds of manual rebuilding in Coot (*72*) and reciprocal space refinement in PHENIX (*73*), and geometry optimization using Rosetta-Phenix refinement (phenix.rosetta_refine) (*74*).

### Negative-stain electron microscopy

A 100 µg/ml final concentration of the antibodies were made in 1:1 ratio of 0.15% glutaraldehyde in HBS pH 7.4 to 10% GlyHBS pH 7.4. After 5 min incubation in 0.075% glutaraldehyde in 5% GlyHBS, pH 7.4, Tris pH 7.4 was added from a 1M stock to a final concentration of 0.075M to quench the glutaraldehyde. After a 5 minute incubation, samples were stained with 2% uranyl formate. Images were obtained with a Philips 420 electron microscope operated at 120 kV, at 82,000× magnification and a 4.02 Å pixel size. The RELION program was used to perform class averaging of the single-particle images.

### Cryo-electron microscopy

#### Cryo-EM data collection

Cryo-EM imaging was performed on a FEI Titan Krios microscope (Thermo Fisher Scientific) operated at 300 kV, aligned for parallel illumination. Data collection images were acquired with a Gatan K3 detector operated in counting mode with a calibrated physical pixel size of 1.066 Å with a defocus range between −1.0 and −3.5 µm using the Latitude S software (Gatan Inc.). The dose rate used was ∼1.0 e-/Å^2^·s to ensure operation in the linear range of the detector. The total exposure time was 4 s, and intermediate frames were recorded every 0.067 s giving an accumulated dose of ∼60 e-/Å^2^ and a total of 60 frames per image. A total of 6058, 3230 and 2521 images were collected for the DH851.3-, DH898.1- and DH898.4-bound complexes, respectively.

#### Cryo-EM data processing

Cryo-EM image quality was monitored on-the-fly during data collection using automated processing routines. Data processing was performed within cryoSPARC (*75*) including particle picking, multiple rounds of 2D classification, *ab initio* reconstruction, heterogeneous and homogeneous map refinements, and non-uniform map refinements.

For the DH898.1-bound complex dataset, a homogeneous set of ∼122k particles were identified and re-extracted from the original dose-weighted micrographs using a binning factor 2. 3D classification in RELION (*76*) was performed imposing C1 symmetry and using different models generated by cryoSPARC as references for alignment. All models were low-pass filtered to 60 Å before refinement. For the binding site located around a CD4 binding site glycan cluster, we identified one class with 42,745 particles which was consistent with the cryoSPARC reference model and subjected them to further 3D auto refinement. After convergence, the refinement was continued with a soft shape mask and resulted in an overall structure at 6.4 Å resolution. To obtain higher resolution, multibody refinement (*77*) was performed on the refined particles using two individual soft masks (one for the Fab-dimer, another for the Env trimer) created with UCSF Chimera (*78*) and RELION. The overall resolution for the gp120 trimer portion reached 4.7 Å and for the Fab-dimer the estimated resolution was 6.7 Å. At this point we switched to un-binned particles and repeated the same processing strategy, but it didn’t result in significant improvements. We then carried out focused refinement on the Env trimer component using the re-extracted particles and imposing C3 symmetry, which lead to an improved map at 4.3 Å resolution. Based on these refinement parameters, CTF-refine and Bayesian-polishing were performed on the particles. Another round of 3D auto refine was performed using the polished particles and the reference model of the entire complex (with a soft shape mask applied), resulting in an overall resolution of 4.8 Å. Multibody refinement was performed once again based on these updated refinements. The final resolution for the gp120 trimer after post-processing was 4.65 Å and 5.8 Å for the Fab-dimer. For the V3-binding site, 17,510 particles were classified out using the V3-binding reference model generated by cryoSPARC. The first round of 3D auto refinement yielded an 8.14 Å resolution map, and multibody refinement resulted in a 7.6 Å map for the gp120 portion and 8.4 Å for the Fab-dimer. Given the lower resolution of this refinement and the fewer number of particles compared with binding-site1, CTF-refine and Bayesian polishing were not used for this class. In order to visualize the principal motions present in the data identified by the multibody refinement procedure, we reconstructed intermediate volumes using different relative orientations of each body and produced movies using the ‘Volume Series’ tool from ChimeraX (*79*). Additional editing of the videos was performed with Movavi Video Editor (Movavi) to label the different components and provide visual aids.

### Surface Plasmon Resonance (SPR)

The binding of DH717.1 and DH717.1 C76S antibodies to 5 different CH848 Env’s was assessed by surface plasmon resonance on Biacore T-200 (GE-Healthcare) at 25°C with HBS-EP+ (10 mM HEPES, pH 7.4, 150 mM NaCl, 3 mM EDTA, and 0.05% surfactant P-20) as the running buffer. The antibodies were captured on a CM5 chip by flowing 200 nM of the antibody over a flow cell immobilized with ∼9000 RU of anti-human Fc antibody. Binding was measured by flowing over 200 nM solution of Env’s in running buffer. 2G12 was used as a control and it was immobilized on one of the flow cells. The surface was regenerated between injections by flowing over 3M MgCl_2_ solution for 10 s with flow rate of 100 µl/min. Blank sensorgrams were obtained by injection of same volume of HBS-EP+ buffer in place of trimer solutions. Sensorgrams of the concentration series were corrected with corresponding blank curves.

The SPR binding curves of 2G12 and DH717.1 mAbs and Fabs against CH848 10.17 DS.SOSIP and CH848 10.17 DS.SOSIP_N133DN138T trimers were obtained using the Biacore S200 instrument (GE Healthcare) in HBS-N 1X running buffer. Biotinylated CH848 DS.SOSIP trimers were immobilized onto a CM3 sensor via streptavidin to a level of 300-350RU. The 2G12 and DH717.2 mAbs and Fabs were diluted down to 50µg/mL and injected over the SOSIP trimer at 30µL/min for 180s. A blank streptavidin surface was used for reference subtraction to account for non-specific binding. SPR single cycle kinetic and affinity measurements of the 2G12 and DH717.1 mAbs and Fabs were also obtained using the Biacore S200 instrument with the CH848 DS.SOSIP trimers directly immobilized to a level of 300-350RU via streptavidin.

Five sequential injections of DH717.1 mAb and 2G12 mAb diluted from 5nM to 50nM were injected over the immobilized CH848 DS.SOSIP trimers at a flow rate of 50uL/min for 120s per injection. 2G12 Fab2 was injected from 50nM to 500nM and DH717.1 Fab dimer was injected from 250nM to 2000nM over the CH848 DS.SOSIP trimers. The dissociation length of the single cycle injections was 600s followed by regeneration with a 20s pulse of 25mM NaOH. Results were analyzed using the Biacore S200 Evaluation Software (GE Healthcare). A blank streptavidin surface as well as buffer binding were used for double reference subtraction to account for non-specific antibody binding and signal drift. Subsequent curve fitting analysis was performed using the 1:1 Langmuir model with a local Rmax for both 2G12 and DH717.1 mAb and the heterogeneous ligand model for 2G12 Fab2. The reported kinetic binding curves are representative of 2 data sets.

For affinity measurements, the DH717.1 Fab was purified using size exclusion chromatography (SEC-FPLC) in PBS 1x. Approximately 10mg of DH717.1 Fab was loaded onto a Superdex 200 increase 10/300 column using a 500μL loop and run at 0.5 mL/min using an Äkta Pure system (GE Healthcare). DH717.1 Fab peaks were collected via fractionation using a 96-well plate and were analyzed using the Unicorn 7.0.2 software (GE Healthcare). Using a linear regression derived from running protein standards (GE Healthcare) of known molecular weight, the molecular weights of the Fab monomer and dimer peaks were estimated. The monomer and dimer peak fractions were then pooled and concentrated using 10k, 0.5mL centrifugal filters (Amicon) and concentrations were measured using a NanoDrop UV-VIS spectrophotometer. Following SEC purification, DH717.1 Fab monomer and dimer stability were assessed using SDS-PAGE gel electrophoresis and re-analysis by SEC-FPLC. SDS-PAGE gel electrophoresis was performed using the BioRad system and 4-15% TGX stain free gels. 7µg of each DH717.1 Fab fragment was loaded onto the gel under both reducing and non-reducing conditions. 10µg of a Precision Plus Protein standard (BioRad) was also loaded to help verify the size of the fragments. The gel was run at 200V for 35-40 minutes in Tris/Glycine/SDS running buffer. The Fab bands were analyzed via the Gel Doc EZ imager (BioRad). For SEC-FPLC re-analysis, 40µg of each DH717.1 Fab monomer and dimer were loaded onto a Superdex 200 increase 10/300 column using a 100μL loop and run at 0.5 mL/min using the aforementioned Äkta Pure system.

### Data availability

The variable heavy and light chain gene sequences for functionally characterized antibodies were deposited in GenBank, and can be accessed using the following accession numbers; MT470283-MT470354. The data presented in this manuscript, and research materials used in this study are available from Duke University upon request and subsequent execution of an appropriate materials transfer agreement. The cryo-EM data are in the process of being deposited to the electron microscopy database (EMDB).

## SOM FIGURES AND TABLES

**Figure S1.**
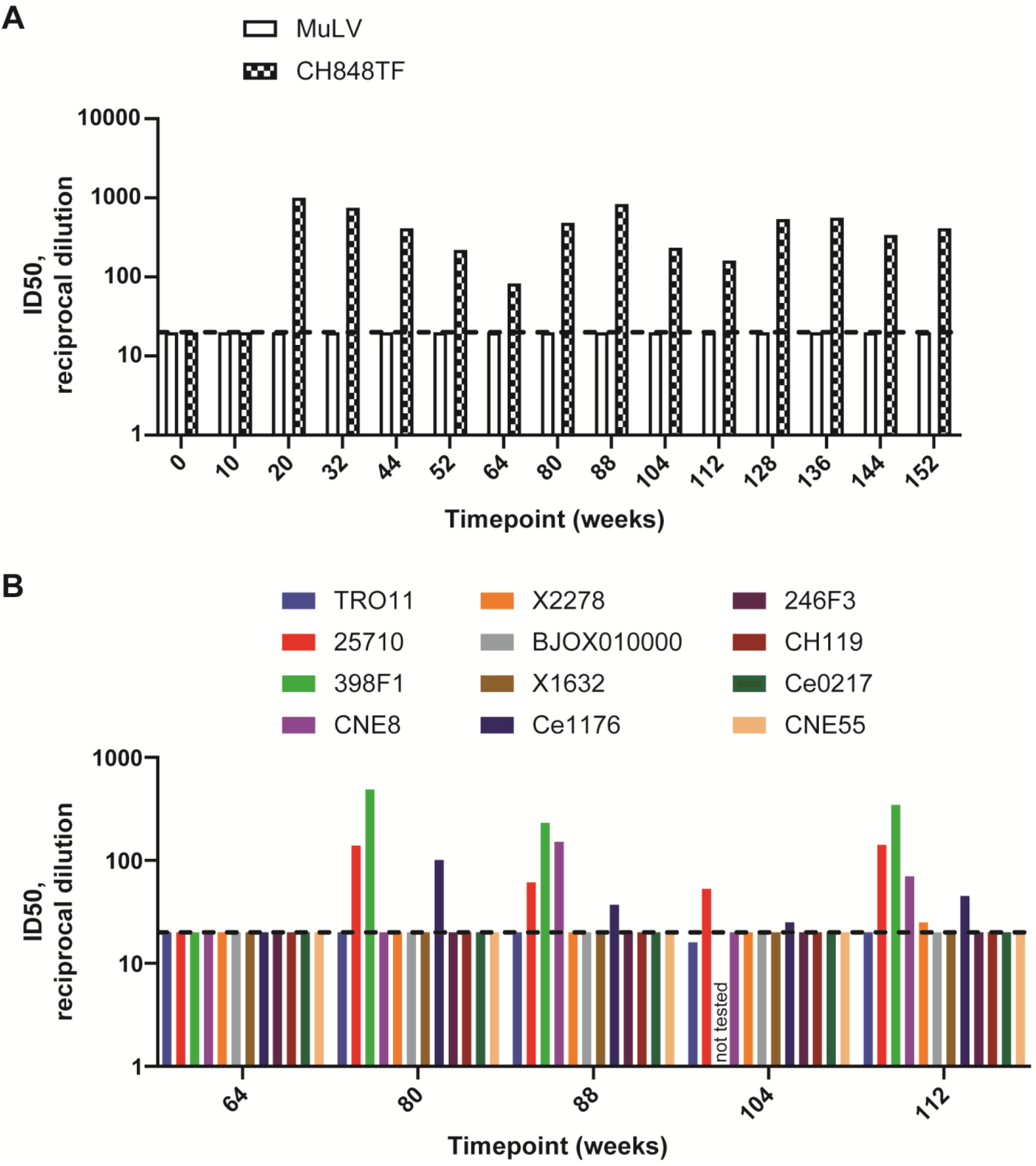
Neutralization profile of SHIV CH848TF-induced plasma antibodies in macaque RM6163. **(A)** Longitudinal plasma neutralization of autologous tier 2 HIV-1 CH848TF. Murine leukemia virus (MuLV) was used as a negative control virus. Plasma was sampled at multiple times (timepoint denoted in weeks) throughout infection. Plasma neutralization titers were combined for longitudinal samples that were studied separately. **(B)** Longitudinal plasma neutralization of the global panel of 12 heterologous tier 2 HIV-1 isolates (*1*) with one exception; 398F1 viral strain was not tested for neutralization by week 104 plasma. MuLV was negative for all timepoints tested (not shown). Plasma neutralizations of HIV-1 strains were tested in TZM-bl cells, and neutralization titers were reported as ID50. Positivity cutoff for neutralization (ID50 <20) is indicated by the dash line.

**Figure S2.**
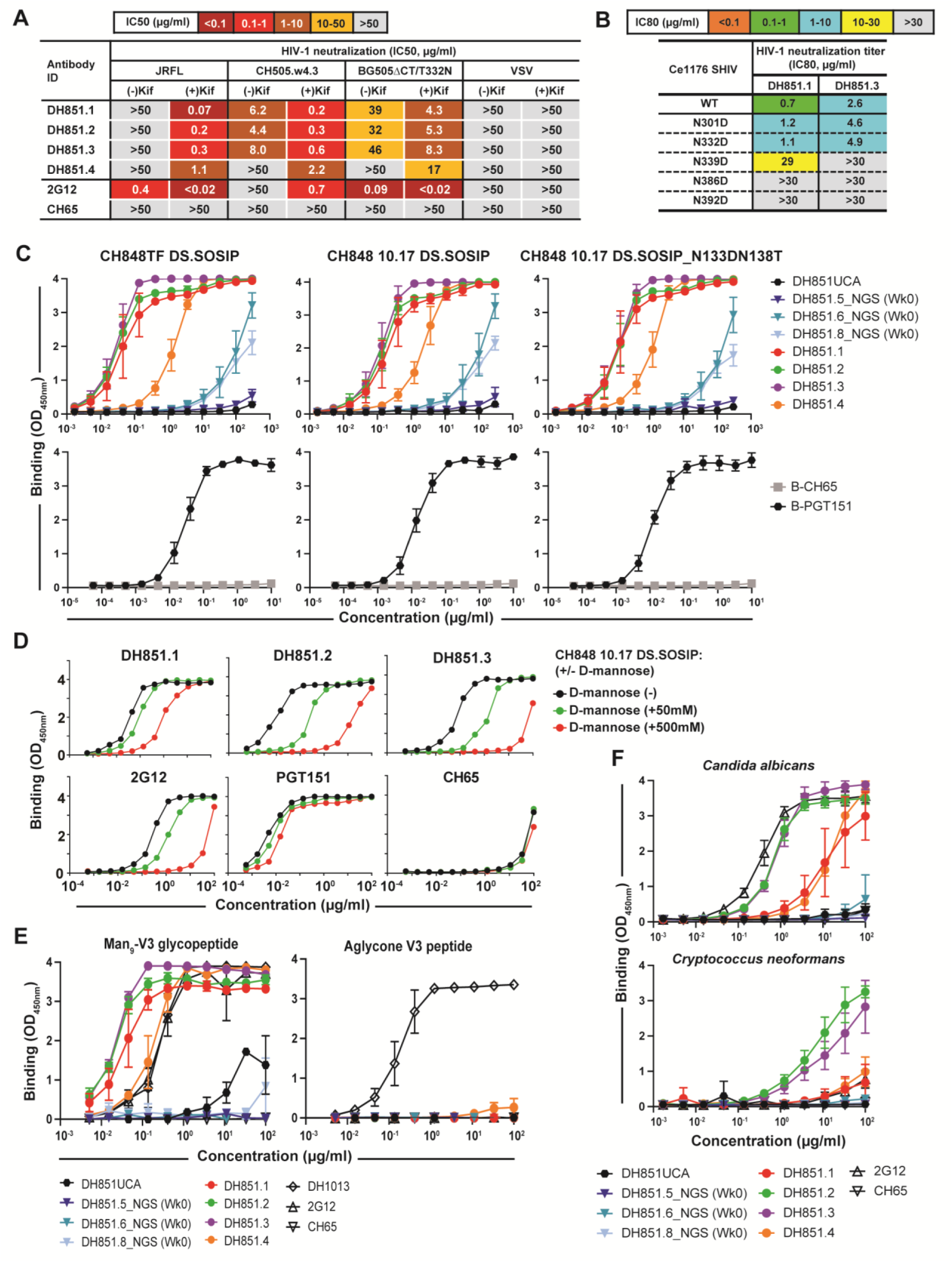
Binding specificities and neutralization profile of SHIV CH848TF-induced DH851 lineage bnAbs. **(A)** Neutralization titers of DH851 mAbs tested against kifunensine [Kif]- treated and non-[Kif]-treated HIV-1 strains. Neutralization was tested in TZM-bl cells and titers reported as IC50 in µg/ml. **(B)** DH851 neutralization epitopes were mapped on a SHIV bearing wild-type or mutant HIV-1 Ce1176-strain Envs. Mutant Envs had deletion of potential N-linked glycosylation sites that constituted HIV-1 V3 glycan bnAb epitope. While DH851 mAbs also neutralized HIV-1 Ce1176 strain, a mutation in the GDIR motif (D325N) did not impact the neutralization sensitivity of HIV-1 Ce1176 (not shown). **(C)** Binding of DH851 lineage mAbs, including week 0, unmutated common ancestor (UCA), and mature antibodies to CH848 strains of SOSIP trimers. Week 0 mAbs had clonally-related heavy chain genes detected via next generation sequencing (NGS) paired with DH851.1 light chain. The UCA was computationally inferred (Cloanalyst software program). Biotinylated (B) PGT151 (Env trimer reactive) and CH65 (influenza reactive) mAbs were tested as control antibodies. Binding levels were measured at OD_450nm_. **(D)** Binding of mature DH851 mAbs to CH848 10.17 DS.SOSIP trimer in the presence (+) or absence (-) of 0.05-0.5M D-mannose, compared with control antibodies 2G12, PGT151 and CH65. **(E)** Binding of DH851 mAbs to Man_9_-V3 glycopeptide and aglycone-V3 peptide. DH1013 (peptide reactive), 2G12 (Env glycan reactive) and CH65 mAbs were tested as control antibodies. **(F)** Binding of DH851 mAbs to heat-killed yeast antigens, *Candida albicans* (1:2000 dilution) or *Cryptococcus neoformans* (1:400 dilution). 2G12 and CH65 were tested as control antibodies. The ELISA data shown in panels C, E and F represent the average binding values and standard deviations (arrows) from triplicate experiments. CH848 10.17 DS.SOSIP binding +/- D-mannose was a single experiment that was in agreement with two additional experiments of DH851 mAbs binding to CH848 10.17 gp120 +/- D-mannose (not shown).

**Figure S3.**
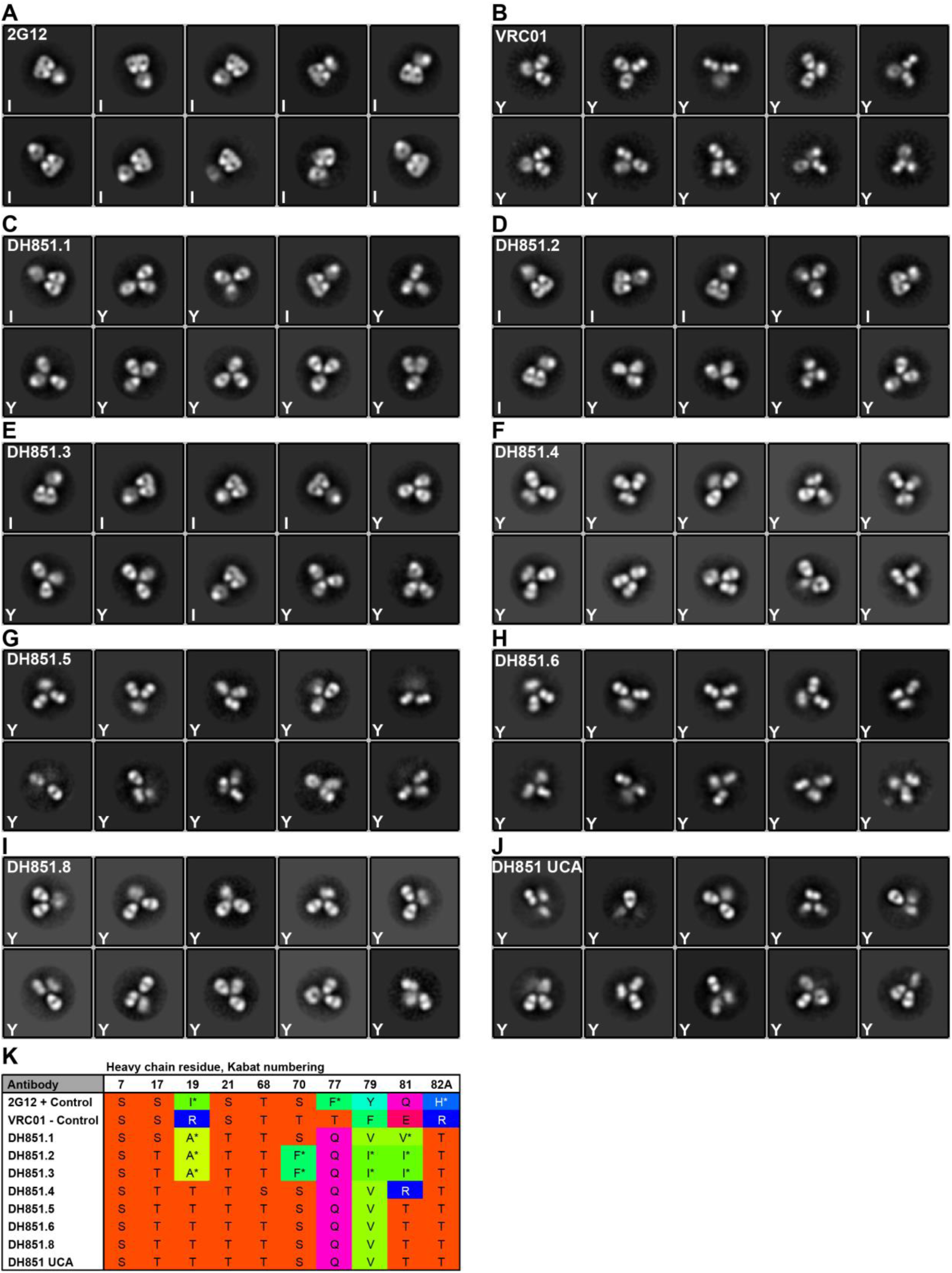
Negative stain class averages of DH851 lineage bnAbs, and control antibodies 2G12 and VRC01. We studied the following recombinant mAbs of the DH851 lineage: inferred unmutated common ancestor (UCA), mature antibodies isolated from single cell sorts (DH851.1-DH851.4), and representative week 0 antibodies bearing clonally-related DH851 VH genes generated via next generation sequencing (DH851.5-DH851.6 and DH851.8). **(A)** Positive control for Fab-dimerization bnAb 2G12 shows all I-shaped antibodies, indicated by “I”. **(B)** Negative control for Fab-dimerization bnAb VRC01 shows all Y-shaped antibodies, indicated by “Y”. **(C)** DH851.1 showing mixture of I- and Y-shaped antibodies. **(D)** DH851.2 showing mixture of I- and Y-shaped antibodies. **(E)** DH851.3 showing mixture of I- and Y-shaped antibodies. **(F)** DH851.4 showing all Y-shaped antibodies. **(G)** DH851.5 showing all Y-shaped antibodies. **(H)** DH851.6 showing all Y-shaped antibodies. **(I)** DH851.8 showing all Y-shaped antibodies. **(J)** DH851UCA showing all Y-shaped antibodies. **(K)** Sequence analysis of key residues within the Fab-dimer interface provides a possible explanation for which lineage members form I-shaped Fab-dimers and which do not. See main text for full description of Fab dimer interface. Top row, heavy chain interface residues are numbered according to standard Kabat numbering (*2*). Amino acids at each position are indicated by their one-letter code and colored according to Taylor (*3*), with polar groups orange, hydrophobic and aromatic groups in shades of green to yellow, positively charged groups blue and negatively charged groups red. Residues that are rare for a particular position, *i.e.* ≤1% in the abYsis database (*4*), are indicated with an asterisk. Positive control 2G12 has four hydrophobic residues participating in contacts within the Fab-dimer interface (*5*), heavy chain residues 19, 77, 79 and 81; residue 81 was considered hydrophobic, because the amphipathic glutamine (Q) side chain was participating in a π-N bond with the aromatic phenylalanine (F) residue 77 of the adjacent heavy chain (*5*). The isoleucine (I) at position 19 is especially important, because a single mutation to arginine (R) at this position is sufficient to completely disrupt Fab-dimerization and domain-swapping in 2G12 (*6*). The 2G12 interface contains three rare interface residues (asterisks), which were not present in the germline sequence (*7*) and thus represent mutations acquired during antibody affinity maturation. In contrast to the I-shaped 2G12, the Y-shaped antibody VRC01 has only a single hydrophobic residue, no rare residues, and three charged residues in these positions, including an arginine at position 19. The three DH851 antibodies that showed I-shaped, Fab-dimerized antibodies, DH851.1 to DH851.3, have three or four hydrophobic interface residues, most of them rare, and no charged residues. In contrast, DH851.4 to DH851.6, DH851.8 and the DH851UCA have only a single hydrophobic interface residue and these antibodies are all Y-shaped, similar to the negative control VRC01.

**Figure S4.**
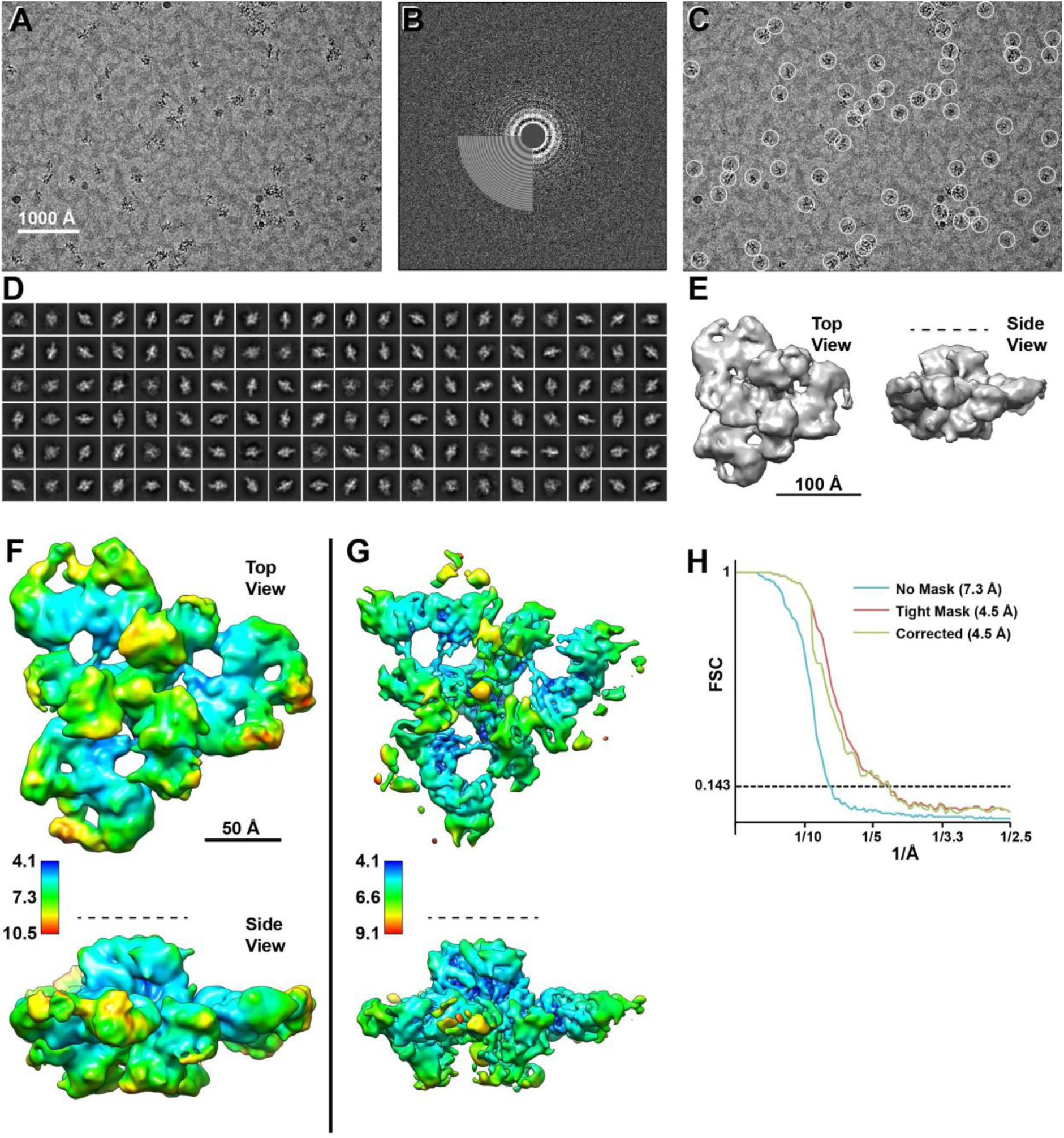
Cryo-EM data processing summary for DH851.3 complex. **(A)** Representative micrograph; **(B)** Power spectrum of micrograph, and fitted contrast transfer function; **(C)** Picked particles in white circles; and **(D)** Representative 2D class averages. **(E)** *Ab initio* volume seen in top and side view. Dashed line in side view indicated viral membrane location. **(F)** Refined 3D map, without filtering or sharpening and colored by local resolution from 4.1 to 10.5 Å, blue to red. **(G)** Map after local filtering and B-factor sharpening. Fab constant domains are noisy and at this contour level mostly disappear and are only seen as small, disconnected blobs, but greater details of the complex can be seen. **(H)** Gold standard Fourier Shell Correlation (FSC) curves indicated global resolution ranging from 4.5 to 7.3 Å.

**Figure S5.**
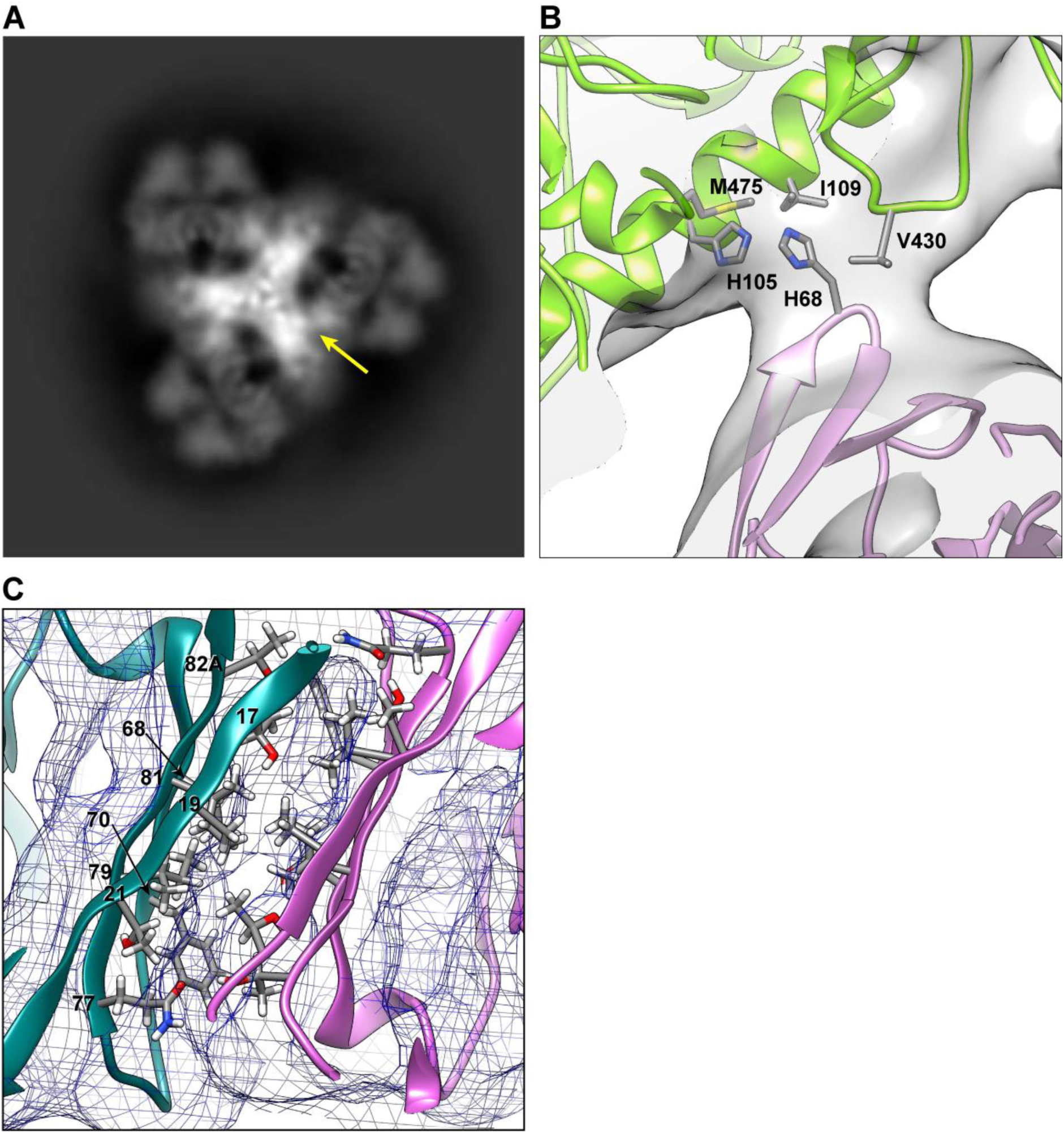
DH851.3 light chain contact to adjacent Env protomers and Fab-dimer interface. **(A)** Projection image of 3D map. Arrow marked the contact of one Fab-dimer to the adjacent protomer. Similar contacts (unmarked) were seen for the other two Fab-dimers. **(B)** Close up of the contact between the light chain (pink) and the Env gp120 (green), with the map shown as a transparent surface. Contact was between histidines 68 (H68) of the light chain and gp120 residues histidine 105, methionine 475, isoleucine 109 and valine 430 (H105, M475, I109, and V430, respectively). **(C)** Close up view of DH851.3 Fab-dimer interface with key interface residues of one chain indicated.

**Figure S6.**
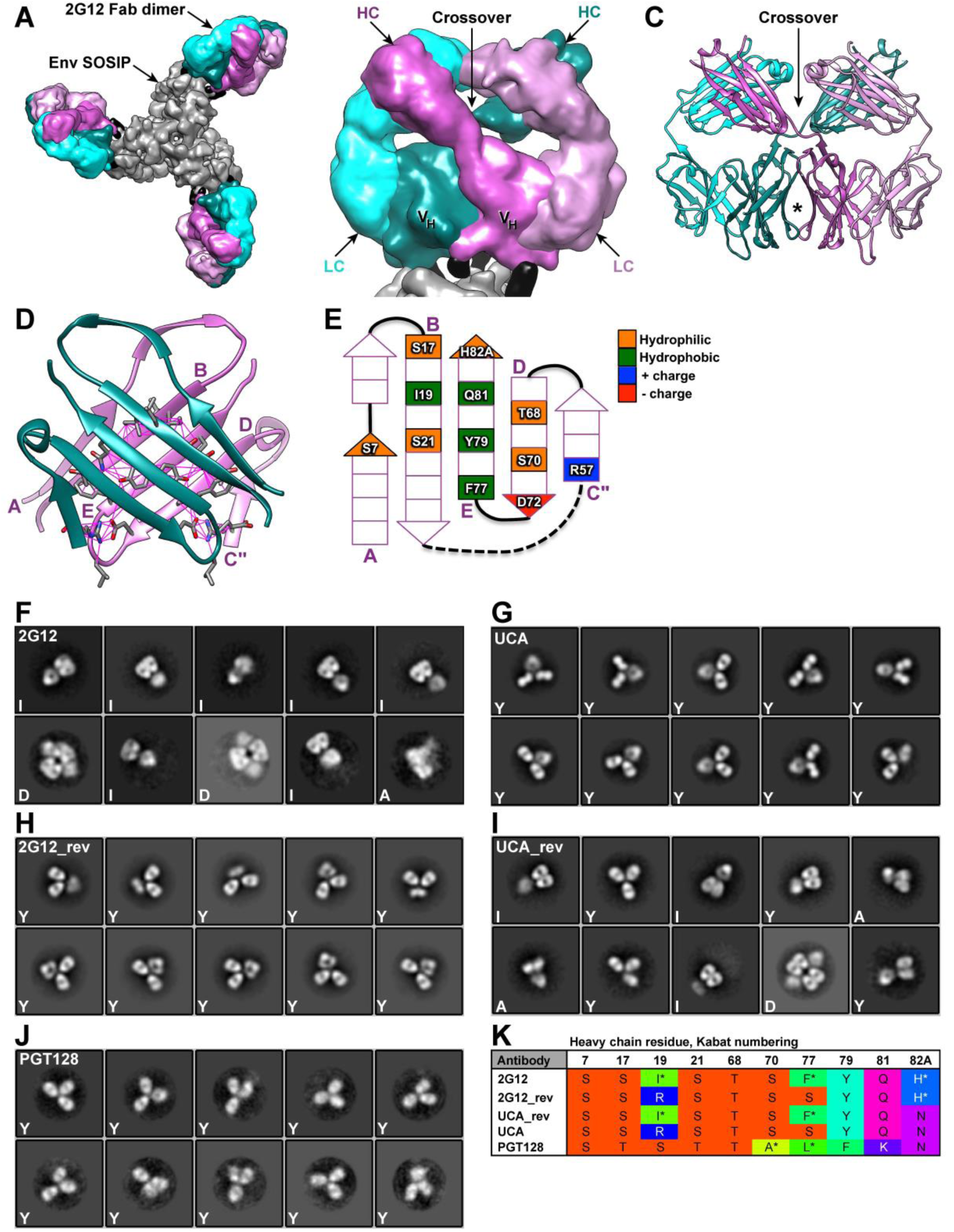
2G12 Domain-swapped Fab-dimer structure and NSEM analysis. **(A)** Published cryo-EM map EMD-8981 with three 2G12 Fab-dimers bound to BG505 SOSIP trimer (*8*). SOSIP trimer = gray; glycans = black; Fab heavy chains = teal or dark pink; Fab light chains = cyan or light pink. **(B)** Close-up of Fab-dimer showed each heavy chain (HC) crossed over one another to bind with both light chains (LC), and with each other via the V_H_ domains (V_H_). **(C)** Atomic model from published X-ray crystal structure PDB 6N32 (*5*) showed the heavy chains crossed over one another and the primary interface (star) between the two V_H_ domains. **(D)** The beta-sandwich view of the interface (starred region in K, turned 90°) showed the beta-strands of the two heavy chains were angled relative to one another. Pink beta-strands were labeled A-E. **(E)** Schematic diagram of interface surface, corresponding to labeled strands in D, showied a central hydrophobic patch (green) surrounded by hydrophilic residues (orange), as well as a charged pair, residues 57 and 72 (blue, red), which formed a pair of complementary salt bridges between the two heavy chains (*5*). **(F)** We expressed 2G12 as an intact IgG, and NSEM indicates the presences of I-shaped IgG molecules (I) as well as IgG dimers (D), as has been previously reported (*9, 10*). **(G)** In contrast, NSEM of the 2G12 UCA indicates all Y-shaped (Y) antibodies. **(H)** Huber *et al* (JVI 2010) reported eight key residues in the heavy chain that were associated with domain-swapping in 2G12. We reverted those 8 residues in the 2G12 wild-type sequence to the corresponding UCA residues (2G12_rev) and, as expected NSEM shows only Y-shaped antibodies. **(I)** We also made the complementary reversion, changing the 8 residues of the UCA sequence to the wild-type residues (UCA_rev) and NSEM shows a mixture of I-shaped and Y-shaped antibodies, as well as IgG dimers. **(J)** As a negative control, we expressed bnAb PGT128, which binds to a V3 peptide-glycan epitope, and NSEM showed all Y-shaped antibodies. **(K)** Fab-dimer interface residues, as described in Figure S3K. Of interest, PGT128 has three hydrophobic interface residues, two of them rare, and in this sense is similar to wild-type 2G12 and might be expected to Fab-dimerize. However, PGT128 also contains a positively charged lysine (K) at residue 81, which presumably prevents Fab-dimerization in the same way that the I19R mutation was shown to prevent Fab-dimerization in 2G12 (*6*).

**Figure S7.**
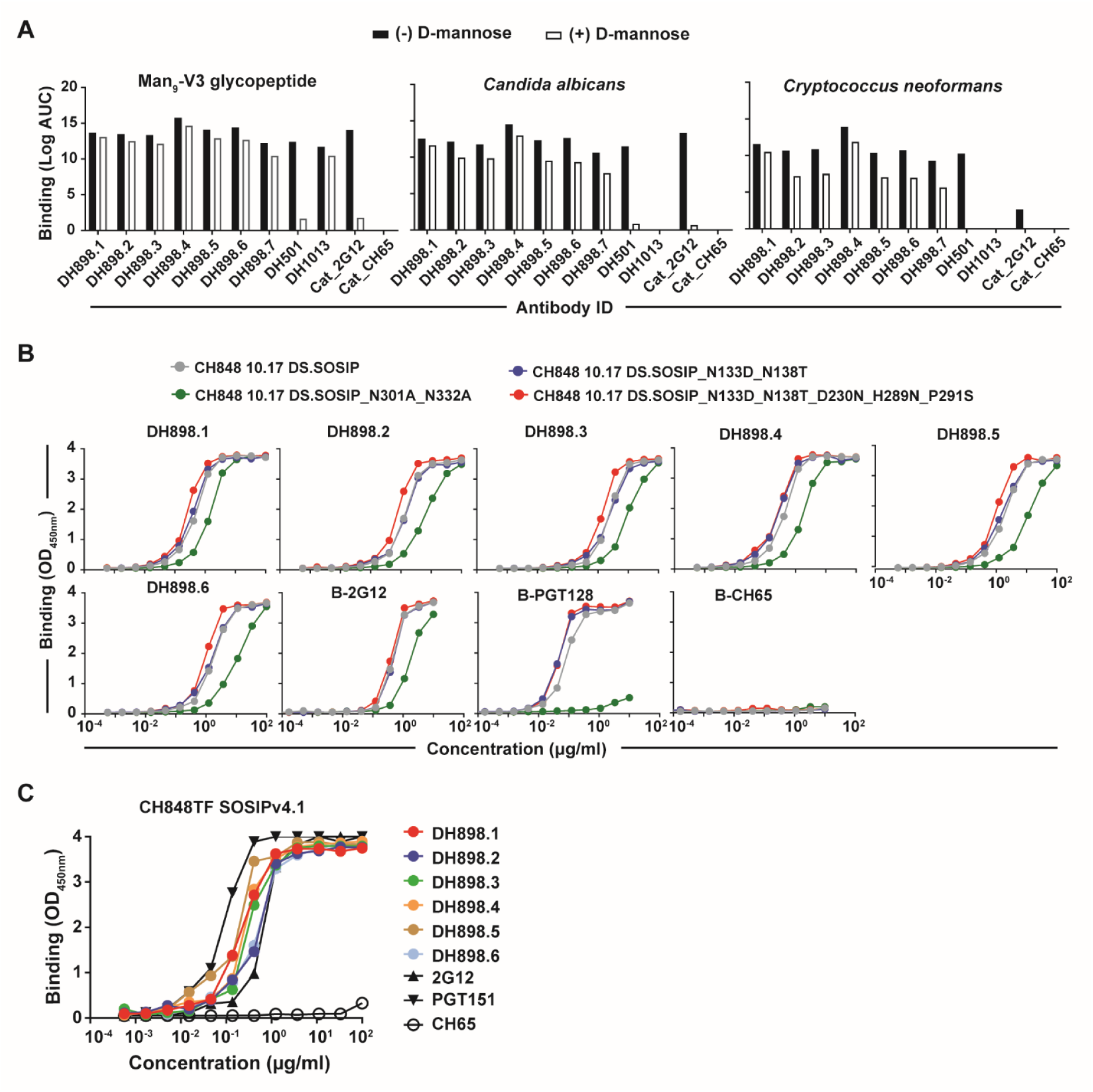
Binding specificities of SHIV CH848TF-induced DH898 lineage antibodies. **(A)** Binding titers of DH898 mAbs to Man_9_-V3 glycopeptide, and heat-killed yeast antigens *Candida albicans* (1:2000 dilution) or *Cryptococcus neoformans* (1:400 dilution), in the absence (black bars) or presence (white bars) of 1M D-mannose. Env glycan- (DH501 and 2G12), peptide- (DH1013) and influenza- (CH65) reactive mAbs were tested as control antibodies. Binding titers were reported as Log Area Under the Curve (AUC). Data shown were from a single ELISA, but were in agreement with an independent experiment of DH898 mAbs binding to Man_9_-V3 glycopeptide and yeast antigens in the absence or presence of 0.5M D-mannose (not shown). **(B)** Binding levels of DH898 mAbs for reactivity with CH848 10.17 DS.SOSIP wild-type and mutant trimers in a single ELISA experiment. Mutations in CH848 10.17 DS.SOSIP trimers included deletions of potential N-linked glycan sites in the V3 (N301A_N332A) or V1 (N133D_N138T) regions. Mutations also included amino acid insertions that filled glycan holes on the trimer (D230N_H289N). Binding levels were measured at OD_450nm_. CH848 10.17 SOSIP trimers were captured using PGT151. Biotinylated (B) 2G12, PGT128 (V3 glycan bnAb) and CH65 mAbs were tested as control antibodies. **(C)** Binding levels of DH898 mAbs to CH848TF SOSIPv4.1 trimer in one ELISA, representative of at least two independent experiments. CH848TF SOSIP trimer was captured using anti-AVI mAb. 2G12, PGT151 and CH65 were tested as control antibodies.

**Figure S8.**
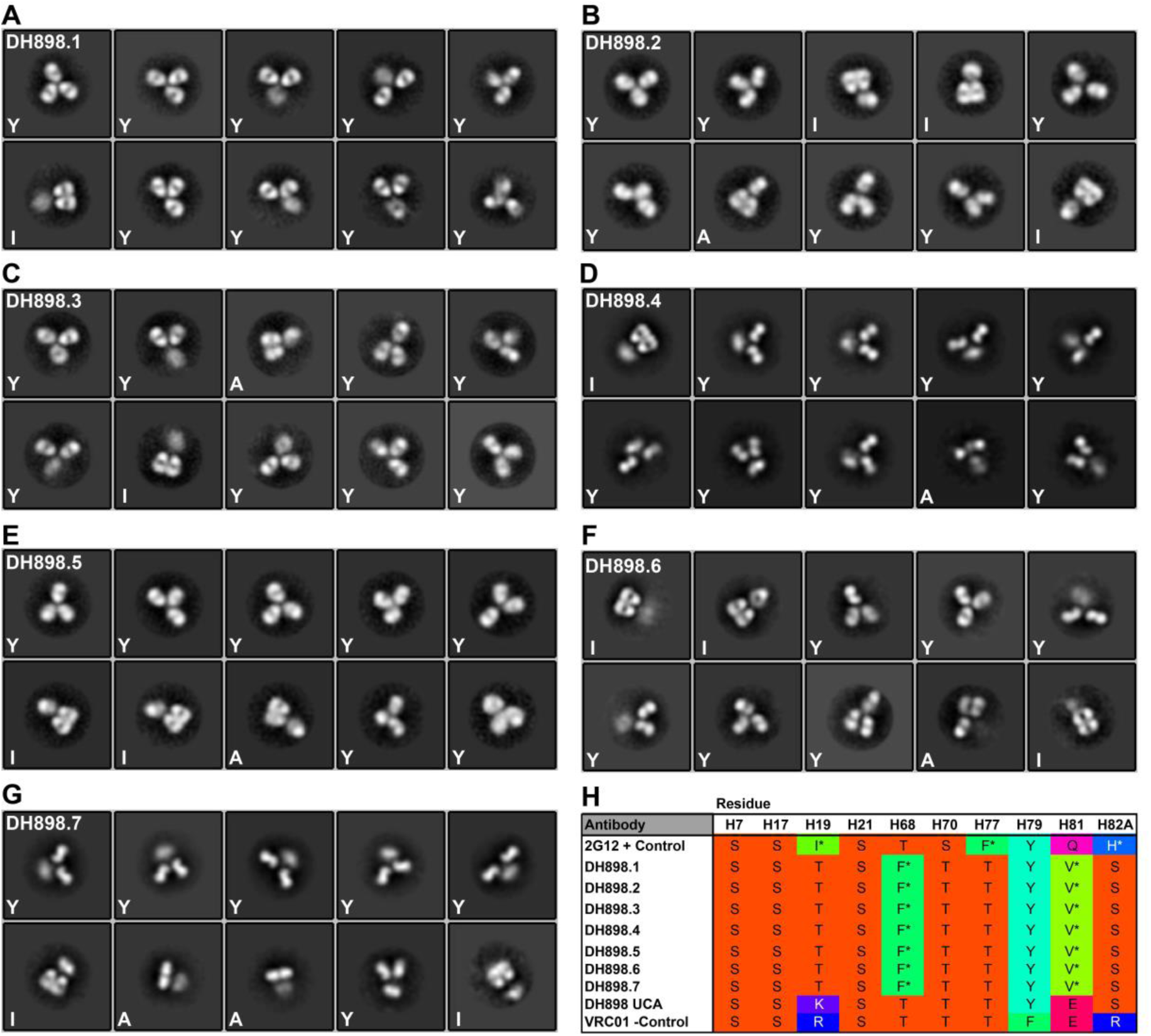
NSEM and sequence analysis of DH898 lineage antibodies. (A-G) NSEM class averages for DH898 mAbs with linage member indicated at upper left of each panel. At the lower left corner, each class average was identified as Y-shaped (Y), I-shaped (I), or ambiguous (A). **(H)** Sequence analysis of Fab-dimer interface that showed three hydrophobic or aromatic residues in the interface. Rare residues for each position marked with an asterisk. Top row indicated corresponding residues for Fab-dimerized 2G12 as a positive control, and the bottom row indicated VRC01 as a non-Fab-dimerized negative control.

**Figure S9.**
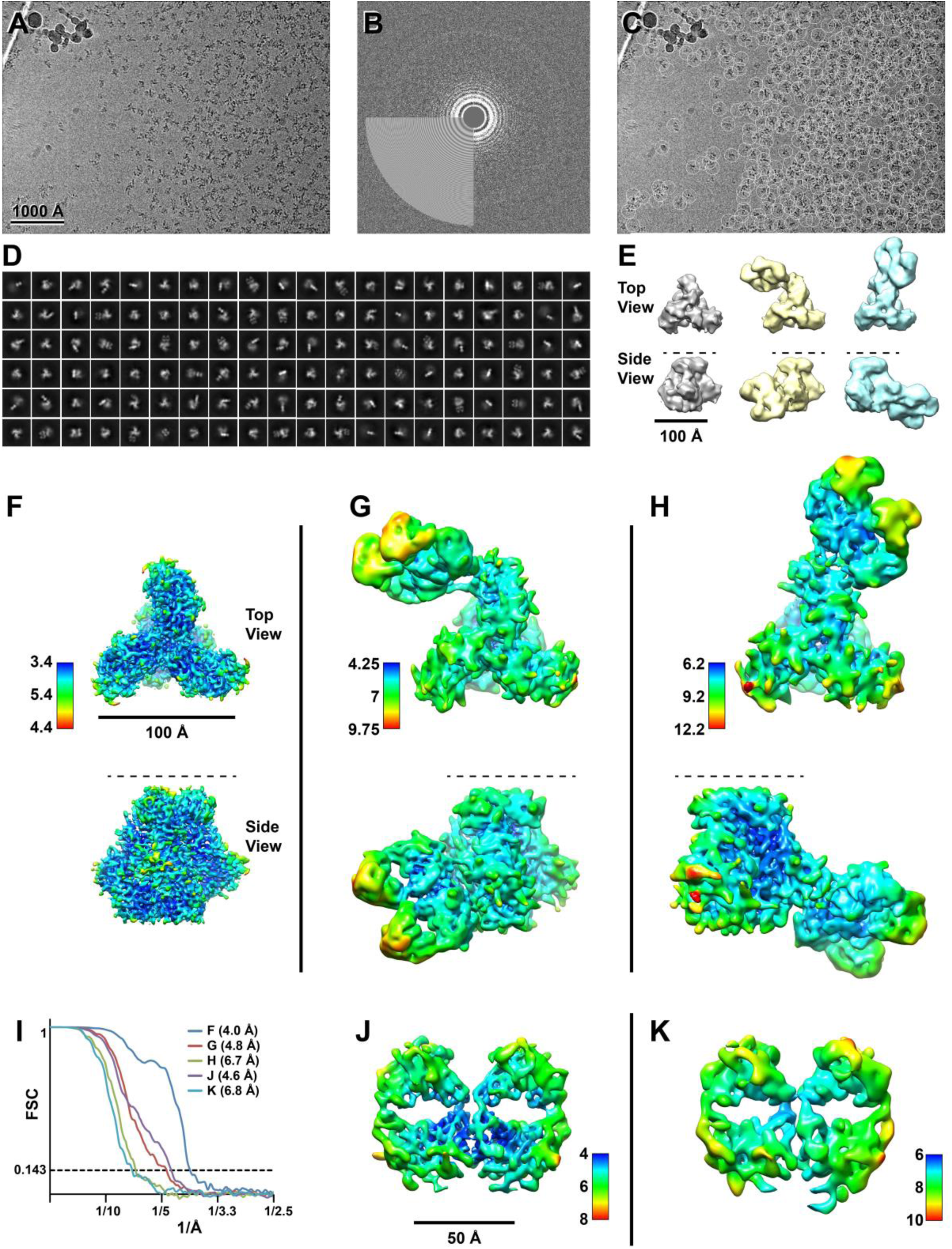
Cryo-EM data processing summary for DH898.1 complex. **(A)** Representative micrograph; **(B)** Power spectrum of micrograph, and fitted contrast transfer function; **(C)** Picked particles in white circles; and **(D)** Representative 2D class averages. **(E)** *Ab initio* volumes that showed three particle populations: free trimer, Fab-dimer bound to glycans near the CD4 binding site, and Fab-dimer bound to glycans near the base of the V3 loop (left to right) seen in top and side views (top to bottom). Dashed line in side view indicated viral membrane location. **(F)** Top and side view of free SOSIP trimer colored by local resolution. **(G)** Top and side view of SOSIP trimer with Fab-dimer bound to glycans near the CD4 binding site. **(H)** Top and side views of SOSIP trimer with Fab-dimer bound to glycans near the base of the V3 loop. **(I)** Gold standard Fourier Shell Correlation (FSC) curves for F-H and J-K indicating global resolution ranging from 4.0 – 6.8 Å. **(J)** Local refinement of Fab-dimer only from CD4-binding site particle set, colored by local resolution. **(K)** Local refinement of Fab-dimer only from V3-glycan bound particle set, colored by local resolution.

**Figure S10.**
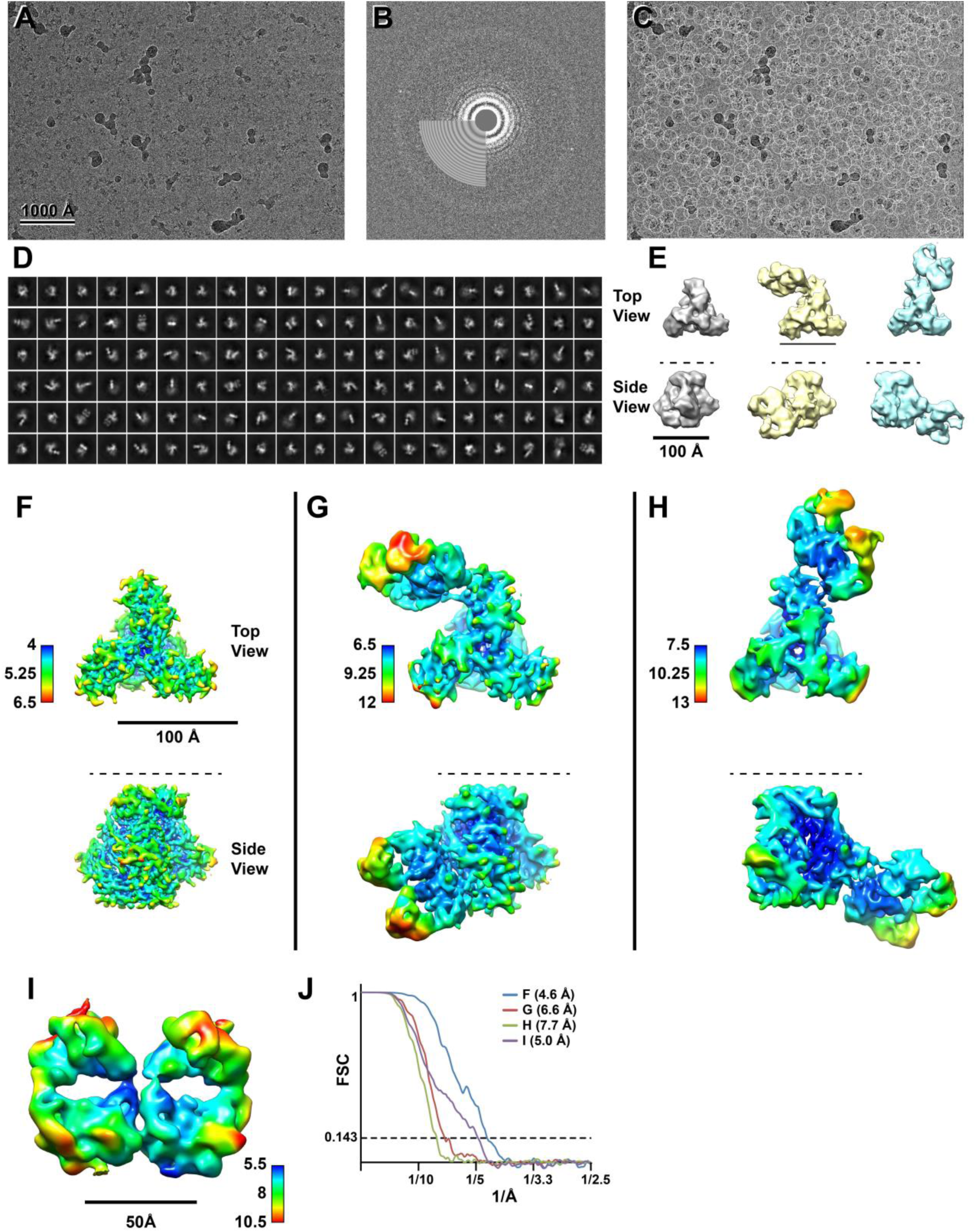
Cryo-EM data processing summary for DH898.4 complex. **(A)** Representative micrograph **(B)** Power spectrum of micrograph, and fitted contrast transfer function **(C)** Picked particles in white circles **(D)** Representative 2D class averages **(E)** *Ab initio* volumes showing three particle populations: free trimer, Fab-dimer bound to glycans near the CD4 binding site, and Fab-dimer bound to glycans near the base of the V3 loop (left to right) seen in top and side views (top to bottom). Dashed line in side view indicates viral membrane location. **(F)** Top and side view of free SOSIP trimer colored by local resolution. **(G)** Top and side view of SOSIP trimer with Fab-dimer bound to glycans near the CD4 binding site. **(H)** Top and side views of SOSIP trimer with Fab-dimer bound to glycans near the base of the V3 loop. **(I)** Local refinement of Fab-dimer only from CD4-binding site particle set, colored by local resolution **((I)** Gold standard Fourier Shell Correlation (FSC) curves for F-I indicated global resolution ranging from 4.6 – 7.7 Å.

**FIGURE S11.**
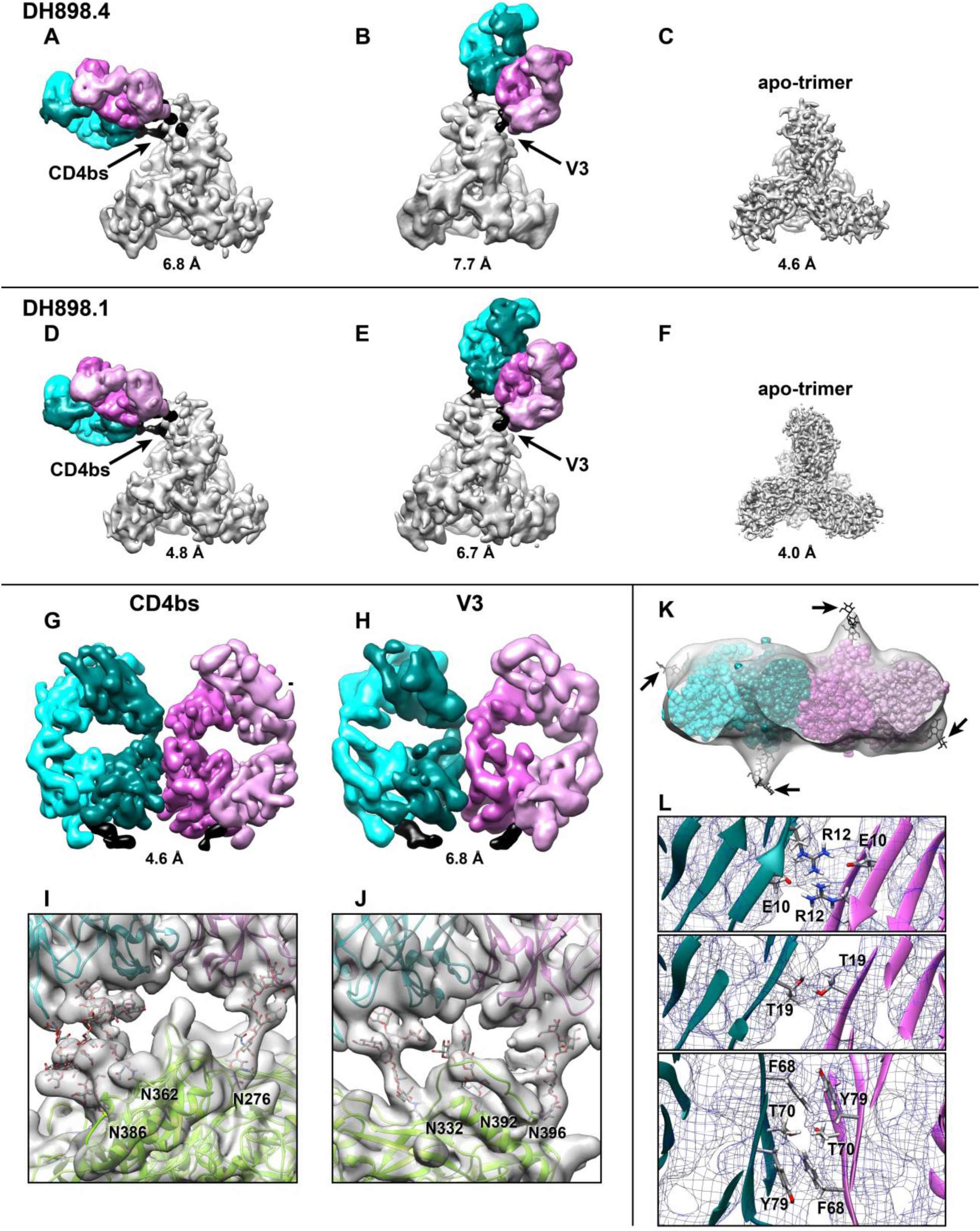
Comparison of six cryo-EM maps and structural details for DH898.1 and DH898.4 complexes. **(A)** Segmented cryo-EM map from DH898.4 dataset showed Fab-dimer bound near the CD4 binding site (CD4bs) of the SOSIP trimer. Gold-standard FSC resolution was indicated below each map. **(B)** Segmented cryo-EM map from DH898.4 dataset showed Fab-dimer bound near the V3-glycan site (V3) of the SOSIP trimer. **(C)** Cryo-EM map from the DH898.4 dataset showed the unliganded SOSIP trimer. **(D)** Segmented cryo-EM map from DH898.1 dataset showed Fab-dimer bound near the CD4bs. Gold-standard FSC resolution was indicated below each map. **(E)** Segmented cryo-EM map from DH898.1 dataset showed Fab-dimer bound near the V3-glycan site. **(F)** Cryo-EM map from the DH898.1 dataset showed the unliganded SOSIP trimer. Within the limits of resolution, there was no apparent difference between corresponding maps for the DH898.4 or DH898.1 datasets. **(G)** Segmented cryo-EM map from local refinement of the DH898.1 Fab-dimer bound near to the CD4bs of the SOSIP trimer. **(H)** Segmented cryo-EM map from local refinement of the DH898.1 Fab-dimer bound to the V3-glycan epitope of the SOSIP trimer. Within the limits of resolution, there was no apparent difference between the two Fab-dimers shown in (G) and (H). **(I)** Close-up view of the DH898.1 epitope near the CD4bs with SOSIP and Fab-dimer models shown as ribbons, glycans shown as sticks, and the cryo-EM map shown as a transparent surface. **(J)** Close-up view of the DH898.1 epitope at the V3-glycan site. Note, figures H and J duplicate Figures 2F and 2G in the main text, respectively, and are shown here for comparison to the structure with DH898.1 bound near the CD4bs of the SOSIP trimer. **(K)** Transverse view of the Fab-dimer, starting at the elbow region and showing the variable domains. View was rotated 90° relative to that in G and H. Cryo-EM map was shown as a transparent surface, Fab variable domain model was shown as spheres. Arrows marked extra map density corresponding to Fab glycans. Glycans were modeled into the density using Coot, and are shown as black sticks. Horizontal arrows indicated glycans attached to heavy chain N26; diagonal arrows indicate glycans attached to light chain N70. N-glycosylation sequons at these sites were conserved among all DH898 members, as well as a third N-glycosylation site at residue N58 within the heavy chain CDR2 region. No density was assignable to this third glycosylation site in the cryo-EM map; furthermore, glycan analysis indicated less than 2% of the antibodies carried glycans at this position (Desaire, H *et al.* unpublished). **(L)** Contacts within the Fab-dimer interface, with heavy chains shown as ribbons, contacting residues as sticks, and the cryo-EM map as mesh; viewing direction – the same as in K.

**Figure S12.**
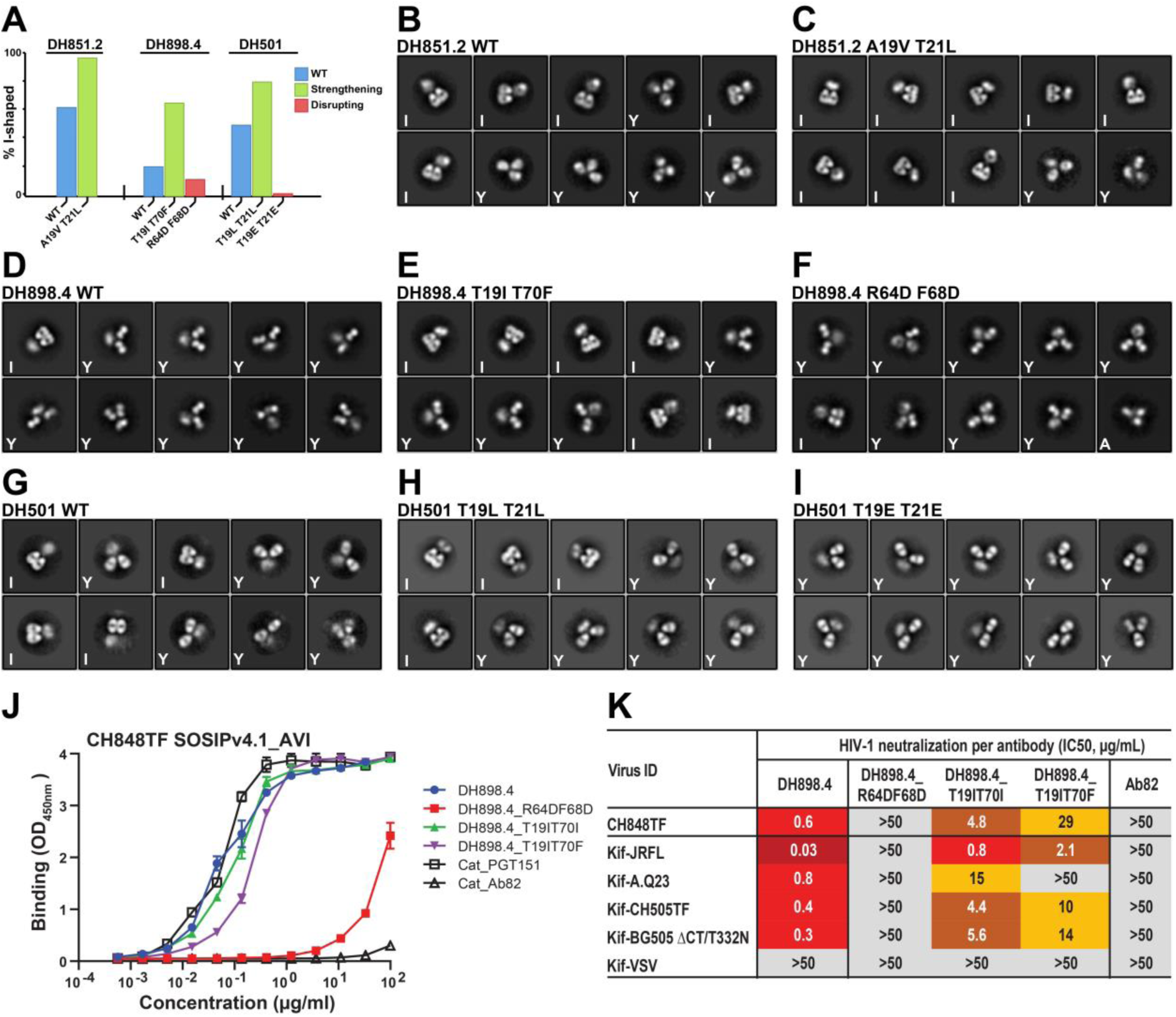
Analysis of FDG antibody mutants designed to strengthen or disrupt the Fab-dimer interface. **(A)** Bar graph indicating the fraction of I-shaped antibodies for wild type (blue bars), strengthening mutants (green bars), or disrupting mutants (red bars) in DH851, DH898 and DH501 FDG antibodies. Mutations were engineered in the antibody heavy chain genes. The % I-shaped was estimated by the fraction of particle images that sorted into I-shaped classes as indicated in each of the NSEM 2D class averages shown in B-I. **(B)** NSEM 2D class averages from DH851.2 wild type (WT). For this and each of the following panels (C-I), the class averages shown represented the average of ∼8,000 to 20,000 individual particle images classified and averaged into ten classes, arranged from the most populated class at the top left to the least populated at the bottom right, and marked as I-shaped (I), Y-shaped (Y), or ambiguous (A). **(C)** NSEM class averages of DH851.2 mAb with strengthening double mutations alanine 19 to valine (A19R) and threonine 21 to leucine (T21L). **(D)** NSEM class averages of wild type DH898.4 mAb. **(E)** NSEM class averages of DH898.4 mAb with strengthening double mutations T19I and T70F. **(F)** NSEM class averages of DH898.4 mAb with disrupting double mutations R64D and F68D. **(G)** NSEM class averages of wild type DH501 mAb. See Figure S17 for full description of DH501 lineage. **(H)** NSEM class averages of DH501 mAb with strengthening double mutations T19L and T21L. **(I)** DH501 mAb with disrupting double mutations T19E and T21E. **(J)** ELISA binding of DH898.4 wild type and mutant mAbs to HIV-1 CH848TF SOSIP trimer. MAbs were tested in technical replicates within a single ELISA and binding levels measured at OD_450nm_; error bars represent standard error of the mean. **(K)** Neutralization of DH898.4 wild type and mutant mAbs against wild-type or non-kifunensine [Kif]- treated autologous HIV-1 CH848, and [Kif]-treated heterologous HIV strains in TZM-bl cells. These data were generated in a single neutralization assay and titers were reported as IC50 in µg/ml.

**Figure S13.**
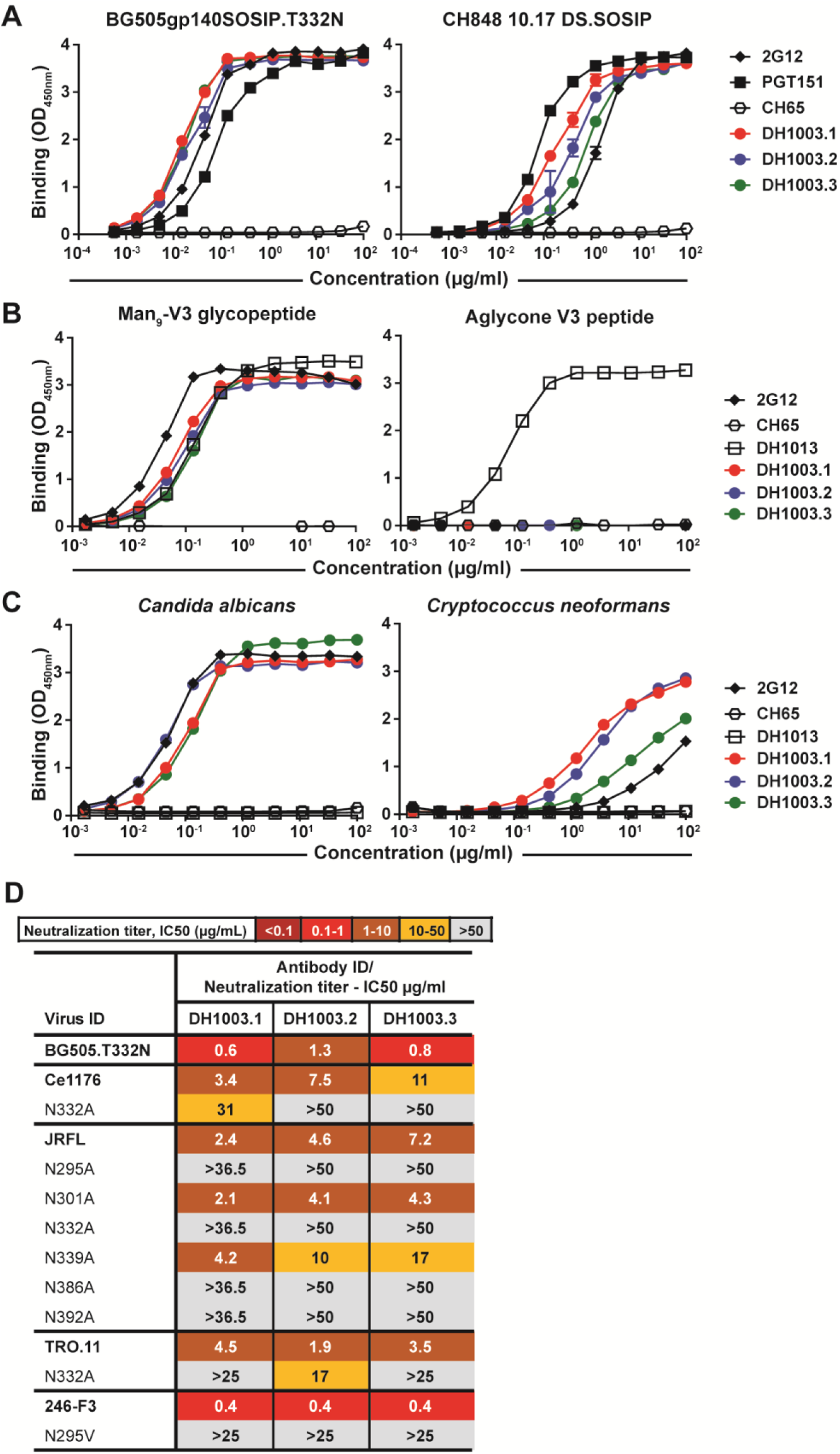
Characteristics of SHIV BG505-induced FDG bnAb lineage, DH1003. **(A)** Binding of DH1003 lineage mAbs to autologous (BG505) and heterologous (CH848 10.17) strains of SOSIP trimers from a representative ELISA. Error bars represented standard error of the mean for duplicate technical replicates per mAb. SOSIP trimers were captured using anti-AVI mAb. 2G12, PGT151 and CH65 mAbs were tested as control antibodies. **(B)** Binding of DH1003 mAbs to Man_9_-V3 glycopeptide and aglycone-V3 peptide in a single ELISA. DH1013, 2G12 and CH65 mAbs were tested as control antibodies. **(C)** Binding of DH1003 mAbs to heat-killed yeast antigens, *Candida albicans* (1:2000 dilution) and *Cryptococcus neoformans* (1:400 dilution) in a single ELISA. 2G12 and CH65 were tested as control antibodies. ELISA binding levels were measured at OD_450nm_. **(D)** Neutralization of DH1003 mAbs against HIV-1 strains bearing wild-type or mutant envelopes with a deletion of potential N-linked glycosylation site that may disrupt the V3 glycan bnAb epitope. Neutralization was tested in TZM-bl cells and titers were reported as IC50 in µg/ml. Neutralization breadth was tested against tier 2 autologous (BG505.T332N) and heterologous HIV-1 strains, including JRFL and nine global panel strains (Ce1176, TRO.11, 246-F3, CH119, 25710, BJOX002000.03.2, X1632-S2-B10, Ce703010217_B6, and CNE55); titers shown were only for viruses that were neutralized by DH1003 mAbs.

**Figure S14.**
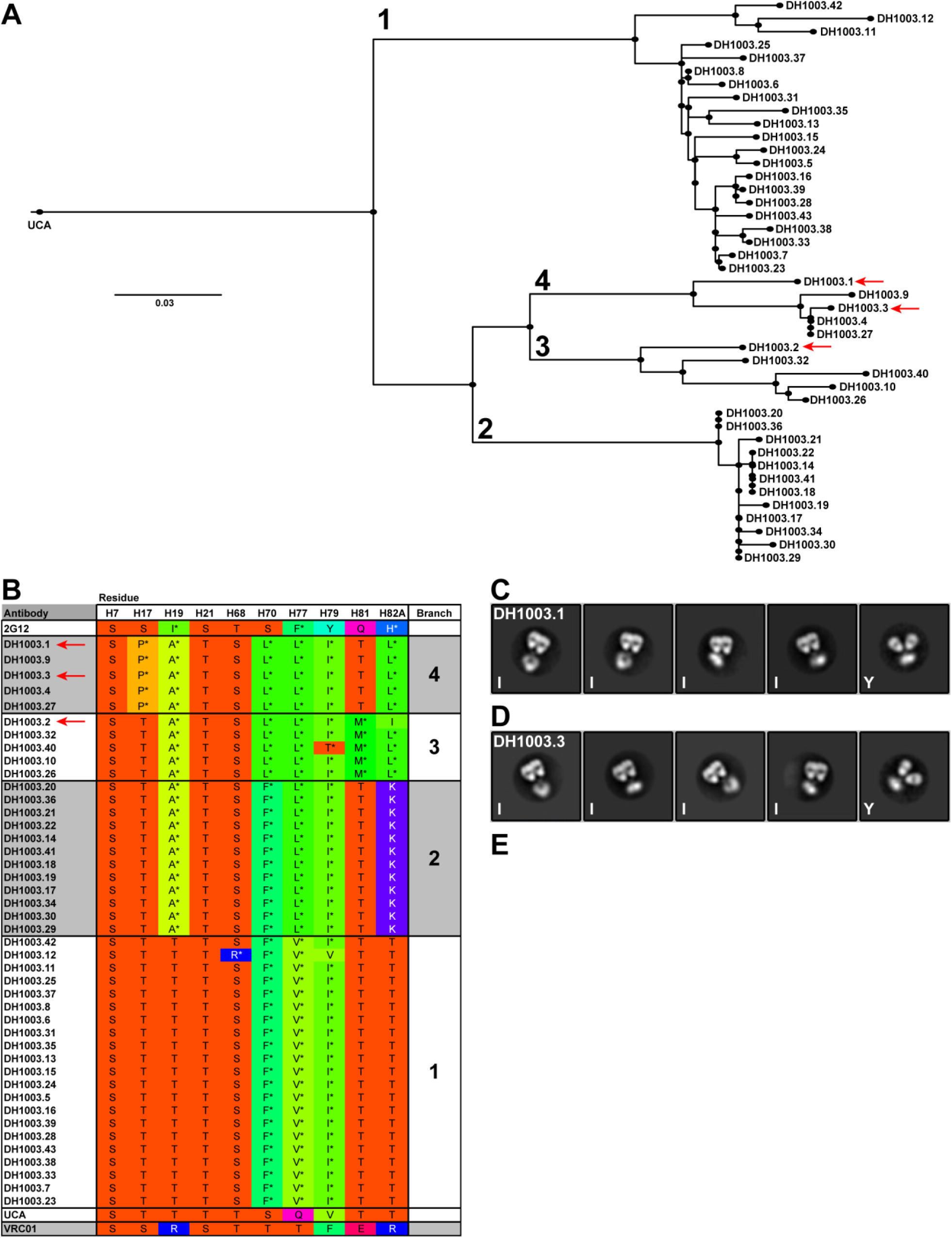
Sequence and NSEM analyses of DH1003 lineage antibodies. **(A)** Phylogram of DH1003 lineage with 42 members of paired heavy and light chain genes from single sorted B cells; the first four branches were numbered 1-4. DH1003.1-DH1003.3 (red arrows) indicated three representative mAbs initially expressed for functional and structural analyses. **(B)** Sequence analysis of DH1003 lineage antibodies showed numerous hydrophobic residues among the key interface residues suggesting a high likelihood of forming Fab-dimers. Rare mutations (<1%) for each residue were indicated with asterisks. The specific pattern of hydrophobic residues within the interface correlated with the phylogram branch points, as indicated in the last column, suggesting that evolution of the lineage correlated with Fab-dimerization; this hypothesis will be explored further in subsequent studies. Top and bottom rows indicated interface sequences of 2G12 and VRC01 as positive and negative controls for Fab-dimerization. **(C)** NSEM class averages from DH1003.1 showed a preponderance of I-shaped antibodies (I) with some Y-shaped antibodies (Y) also present. **(D)** NSEM class averages of DH1003.3. **(E)** NSEM class averages of DH1003.2.

**Figure S15.**
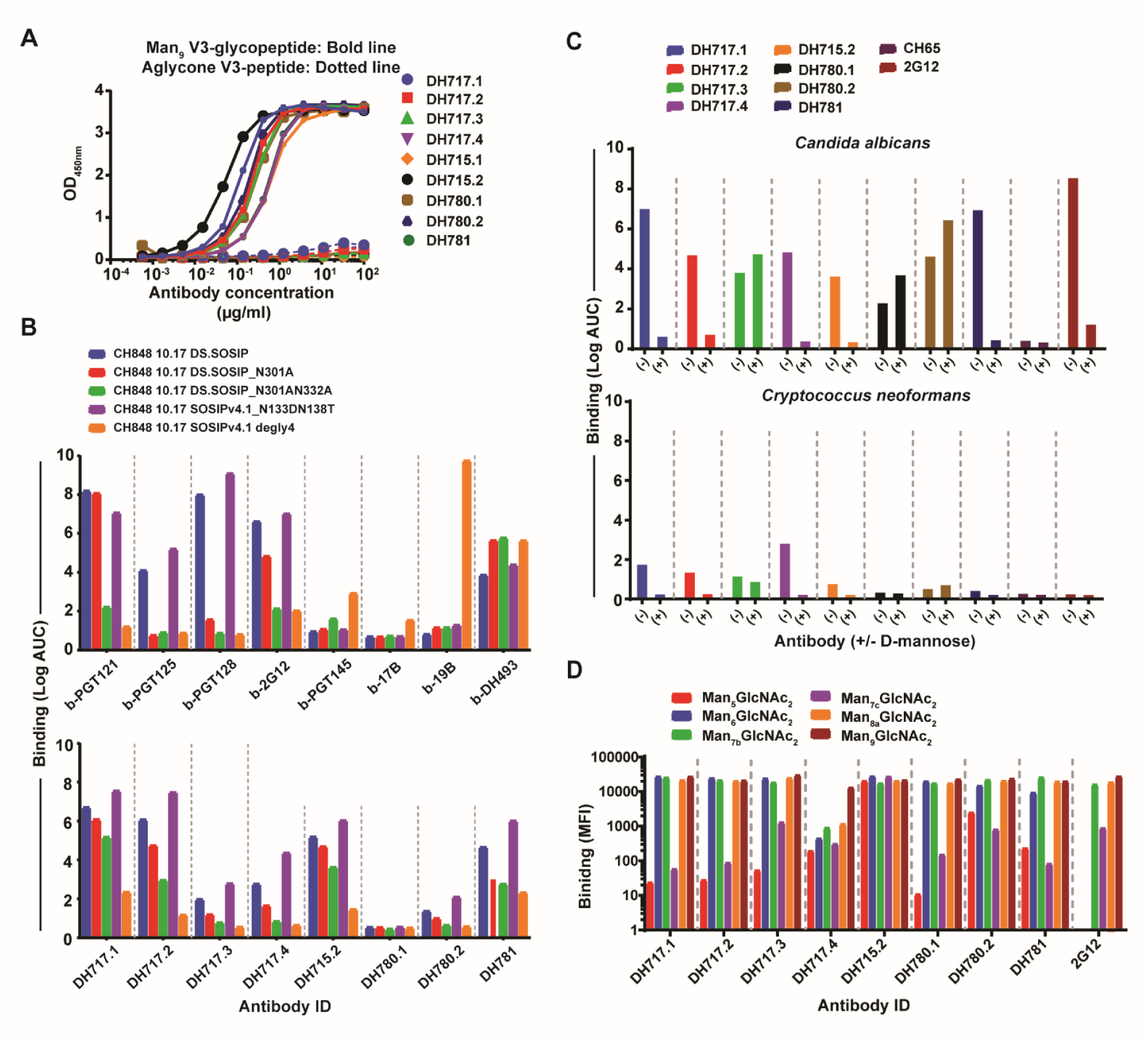
Binding specificities and neutralization profile of HIV-1 vaccine-induced FDG lineage antibodies. **(A)** Binding of DH715, DH717, DH780, and DH871 mAbs with Man_9_-V3 glycopeptide (solid line) or aglycone-V3 peptide (dotted line) in a single ELISA. Binding was measured at OD_450nm_. **(B)** Binding of DH715, DH717, DH780 and DH781 mAbs as well as biotinylated (b) control mAbs (V3 glycan-reactive – PGT121, PGT125 and PGT128; linear V3 reactive – 19B; Env glycan reactive – 2G12; co-receptor binding site – 17B; and CD4 binding site CH235 lineage member – DH493) to CH848 strains of SOSIP trimers in a single ELISA. Binding titers were reported as Log Area Under the Curve (AUC). **(C)** Binding titers of vaccine-induced and control mAbs with heat-killed yeast antigens *Candida albicans* (1:2000 dilution) or *Cryptococcus neonformans* (1:400 dilution). Antibodies were tested in standard diluent or diluent spiked with 0.5M D-mannose. Binding data were representative of duplicate ELISAs. **(D)** Glycan-reactivity of DH715, DH717, DH780, and DH781 mAbs as determined by immunostaining of glycan-coated arrays in a single experiment. Glycan binding was reported as mean fluorescence intensity (MFI) after background subtraction.

**Figure S16.**
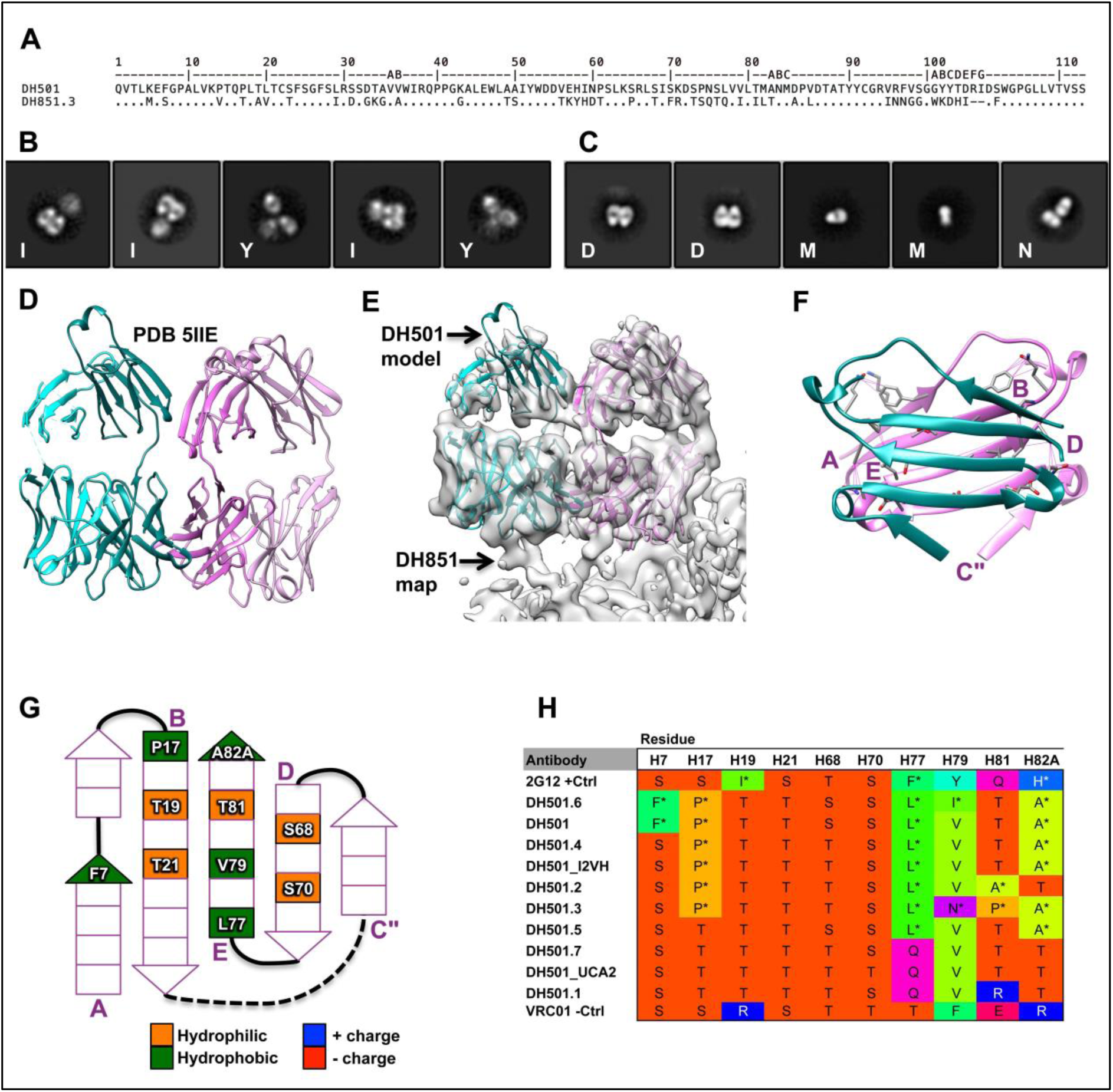
Analysis of the vaccine-elicited DH501 antibody and associated B cell clonal lineage. We previously reported on DH501, a glycan-reactive antibody B-cell lineage isolated from a rhesus macaque after long-term vaccination with a CON-S gp140 (*11*). The current discovery of Fab-dimerization in the DH851 lineage, coupled with a high sequence identity (62%), prompted us to examine the DH501 lineage for its potential to form Fab-dimers. **(A)** Sequence alignment of DH501 and DH851 V_H_ domains. **(B)** NSEM class averages of DH501 IgG mAb showed I-shaped (I) and Y-shaped antibodies, indicating that DH501 was also an FDG antibody. **(C)** NSEM class averages of recombinantly expressed DH501 Fab showed well-formed Fab-dimers (D), monomers (M) and non-specific Fab-dimers (N). The presence of well-formed Fab-dimers in recombinantly expressed Fabs indicated that Fab-dimerization can be driven by the Fabs alone, and does not require the F_c_ domain or the context of an intact IgG. **(D)** The previously published DH501 crystal structure, PDB code 5IIE, showed two Fabs in the unit cell. We now re-interpreted the presence of these two Fabs not as mere crystal contacts, but as a biological Fab-dimer, similar to DH851. **(E)** The DH501 crystal structure, as a rigid body, docks easily into DH851 cryo-EM map. **(F)** DH501 Fab-dimer interface showed the beta strands arranged similar, but slightly more angled than DH851 (compare to Figure 1L). **(G)** Schematic of Fab-dimer interface showed hydrophobic (green) and hydrophilic (orange) residues. **(H)** Sequence analysis of Fab-dimer interface residues for 2G12 (positive control), DH501 lineage, and VRC01 (negative control). Rare mutations (<1%) for each residue were indicated with asterisks. DH501 itself (third row) had five hydrophobic residues within the interface, four of them were considered rare for those positions.

**Figure S17.**
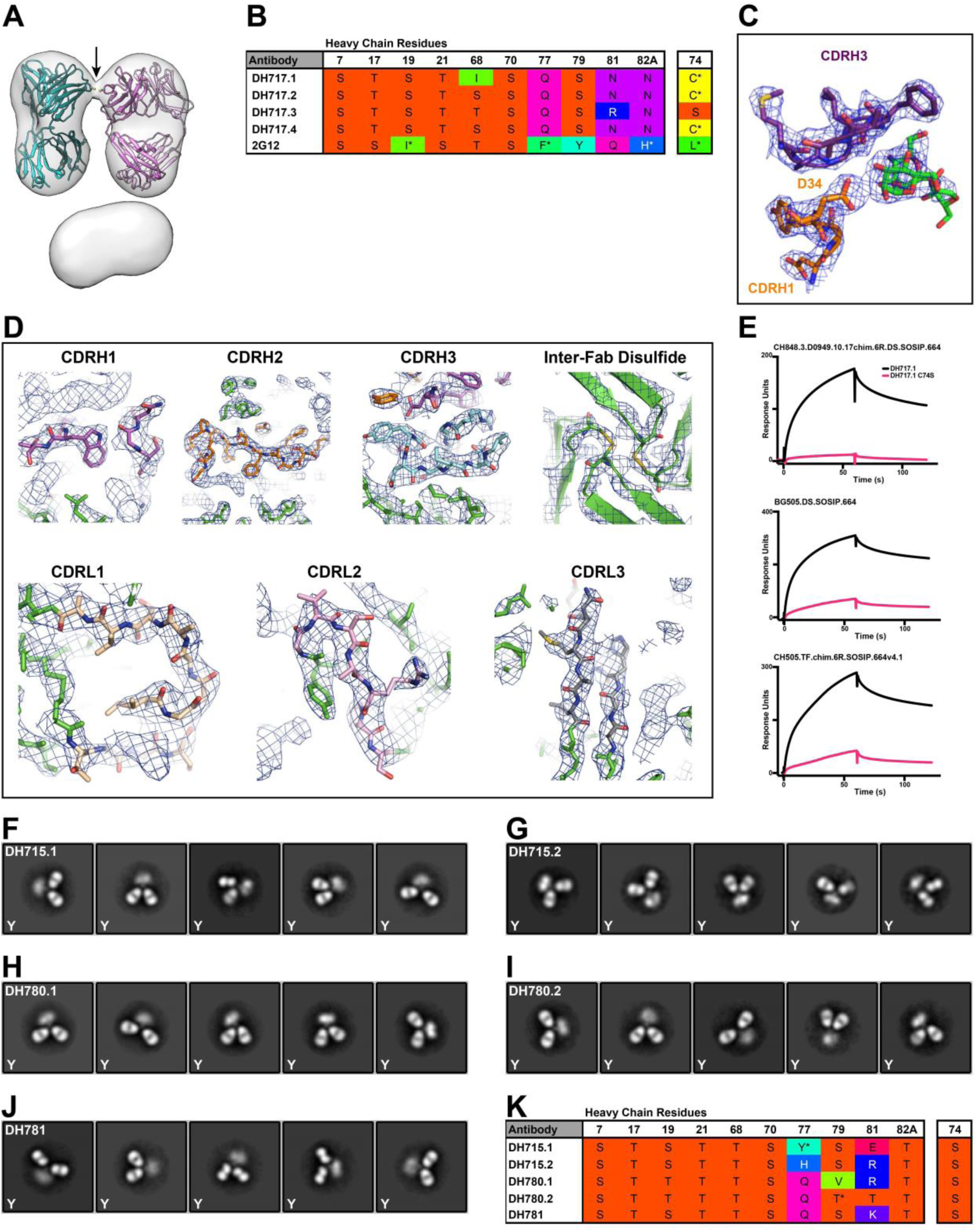
Negative-stain electron microscopy characterization of DH715, DH717, DH780 and DH781 lineage antibodies. **(A)** Negative stain 3D reconstruction of DH717 I-shaped antibody. Map shown as transparent surface. Atomic model of two DH717.1 Fabs shown as ribbon diagrams, fit as rigid bodies into the 20-Å resolution NSEM map using UCSF Chimera’s automatic fitmap function. As fit, the heavy chain cysteine at residue 74 pointed towards one another (arrow) and the terminal sulfurs were 3.5 Å apart. **(B)** DH717 sequence analysis did not show multiple hydrophobic residues, as seen in 2G12 sequence, but showed an unpaired cysteine (C) at heavy chain residue 74 in DH717.1, DH717.2 and DH717.4. **(C)** View of DH717.1 monomer crystal structure showed contact with aspartate, heavy chain residue 34 (D34), and the man-9 ligand (green); electron density shown as mesh. **(D)** Close up views of DH717.1 dimer crystal structure, labeled by region. **(E)** Surface plasmon resonance curves for binding of DH717.1 and DH717.1 C74S mutants to the following three SOSIP trimers (top to bottom): CH818.3.D0949.10.17chim.6R.DS.SOSIP.664; BG505.DS.SOSIP.664; and CH505.TF.chim.6R.SOSIP.664v4.1. Representative 2D class averages of DH715.1 **(F)**, DH715.2 **(G)**, DH7801.1 **(H)**, DH780.2 **(I)** and DH781 **(J)** antibodies showed all Y-shaped (Y) antibody populations. **(K)** Sequence analysis of DH715, DH780 and DH781 lineage antibodies.

**Figure S18.**
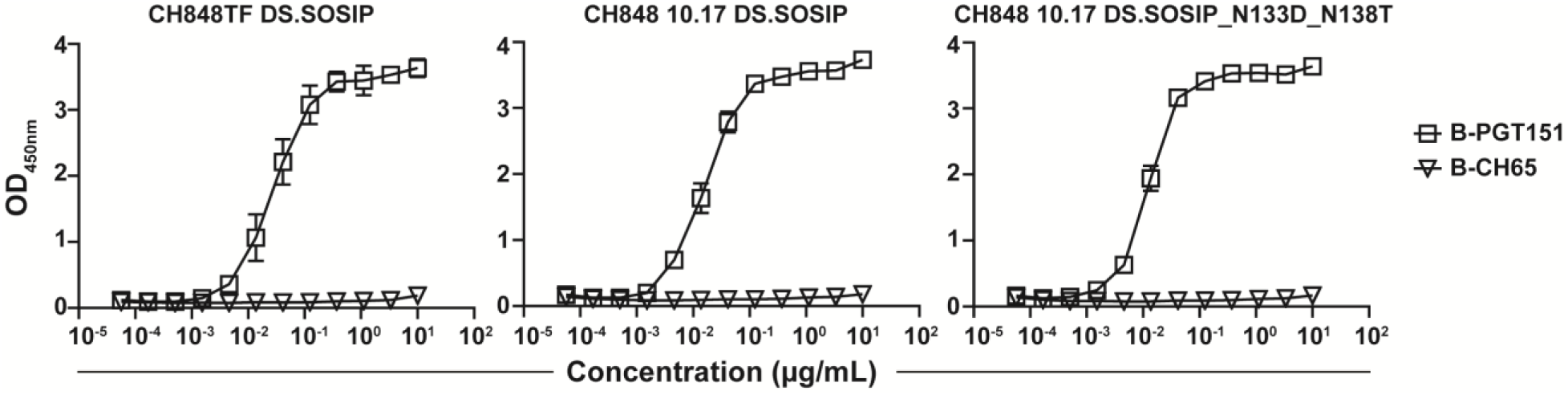
Reactivity of control antibodies with CH848 DS.SOSIP trimers. CH848TF and 10.17 wild-type or mutant SOSIP trimers were captured using PGT151 mAb, and biotinylated (B) PGT151 (V3 glycan bnAb) and CH65 mAbs were tested as control antibodies. Mutations in CH848 10.17 DS.SOSIP trimers included deletions of potential N-linked glycan sites in the V3 (N301A_N332A) or V1 (N133D_N138T) regions (*12*). Binding levels were measured at OD450nm. ELISA data shown are average of 4-5 separate assays where we tested mAbs as controls for DH717 lineage mAbs binding to SOSIP trimers (Figure 5); error bars represent standard error of the mean.

**Figure S19.**
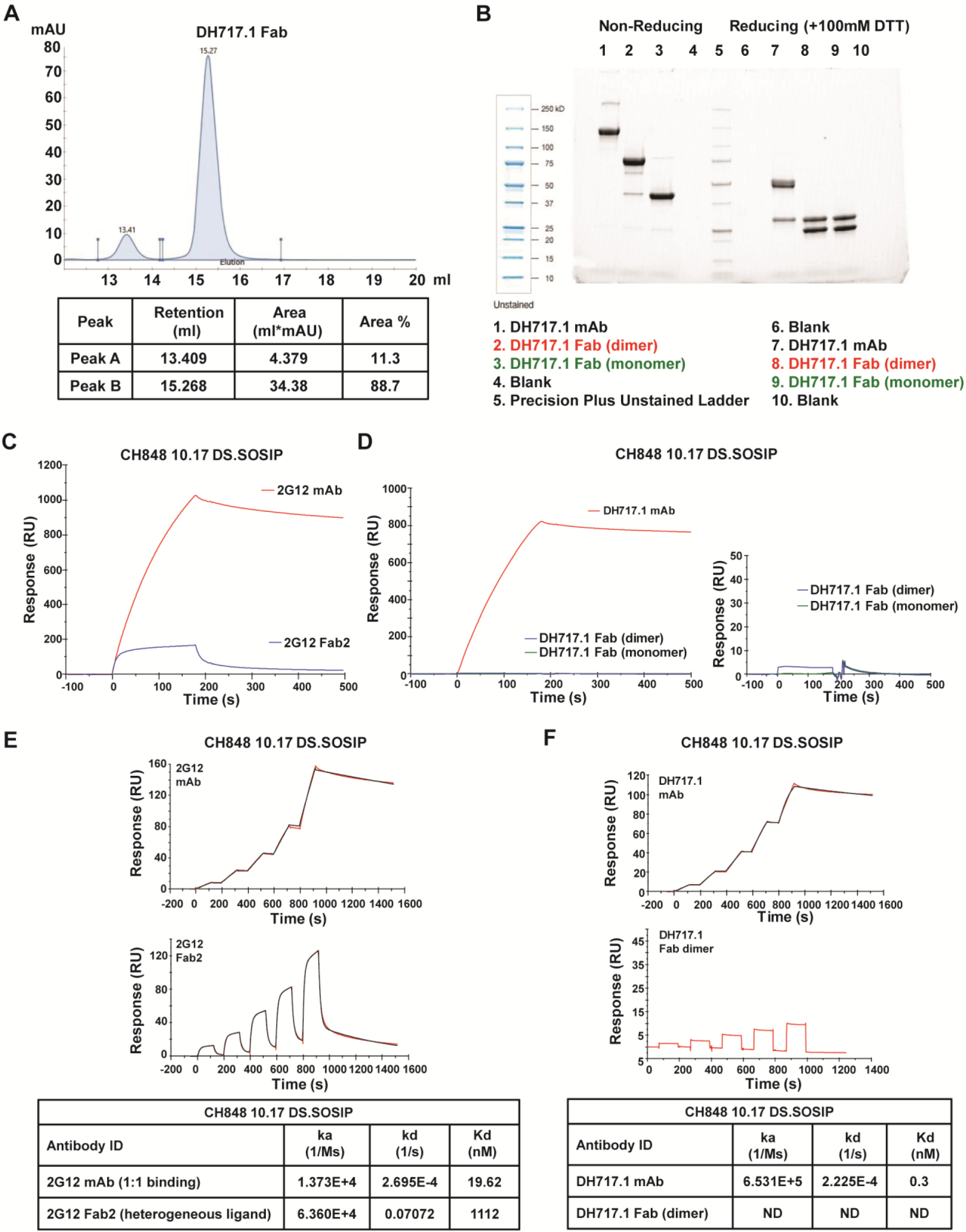
Affinity measurements of FDG Fab monomers and dimers. **(A)** SEC profile of recombinant DH717.1 Fab. Two dominant fractions equivalent to monomers (Peak B) and dimers (Peak A) were observed; these peaks were fractionated and pooled via SEC-FPLC for subsequent binding studies. **(B)** Gel electrophoresis profile of DH717.1 IgG, and monomeric and dimeric forms of Fabs observed in a single recombinant Fab sample. **(C)** Overlay of SPR response curves of 2G12 mAb and Fab2 binding to biotinylated CH848 10.17 DS.SOSIP trimer immobilized via streptavidin. The 2G12 FAb2 was previously obtained by enzymatic digestion of 2G12 IgG (Priyamvada, A *et al.* correspondence). **(D)** Response curves obtained by SPR comparing the binding of DH717.1 mAb and Fab fractions to biotinylated CH848 10.17 DS.SOSIP trimer immobilized via streptavidin. **(E)** SPR single-cycle dose response curves of 2G12 mAb and Fab2 against biotinylated CH848 10.17 DS.SOSIP trimer bound to streptavidin. Curve fitting analyses were performed using the 1:1 Langmuir model for the mAb and the heterogeneous ligand model for the Fab2. A summary of the association (k_a_, 1/Ms) and dissociation (k_d_, 1/s) rates along with relative apparent affinities (K_d_, nM) were shown in the table. The faster kinetic parameters and affinity derived from the heterogeneous ligand model were reported for the Fab2. **(F)** SPR single-cycle dose response curves of DH717.1 mAb and Fab dimer against biotinylated CH848 10.17 DS.SOSIP trimer bound to streptavidin. Curve fitting analyses were performed on the mAb using the 1:1 Langmuir model. The affinity of the Fab dimer against the SOSIP trimer was too weak to be reliably measured. The association (k_a_, 1/Ms) and dissociation (k_d_, 1/s) rates along with relative apparent affinities (K_d_, nM) were summarized in the table for the mAb, whereas no values were reported for the Fab dimer. Data shown are consistent with SPR studies done using CH848 10.17 DS.SOSIP_N133DN138T trimer (not shown).

**Figure S20.**
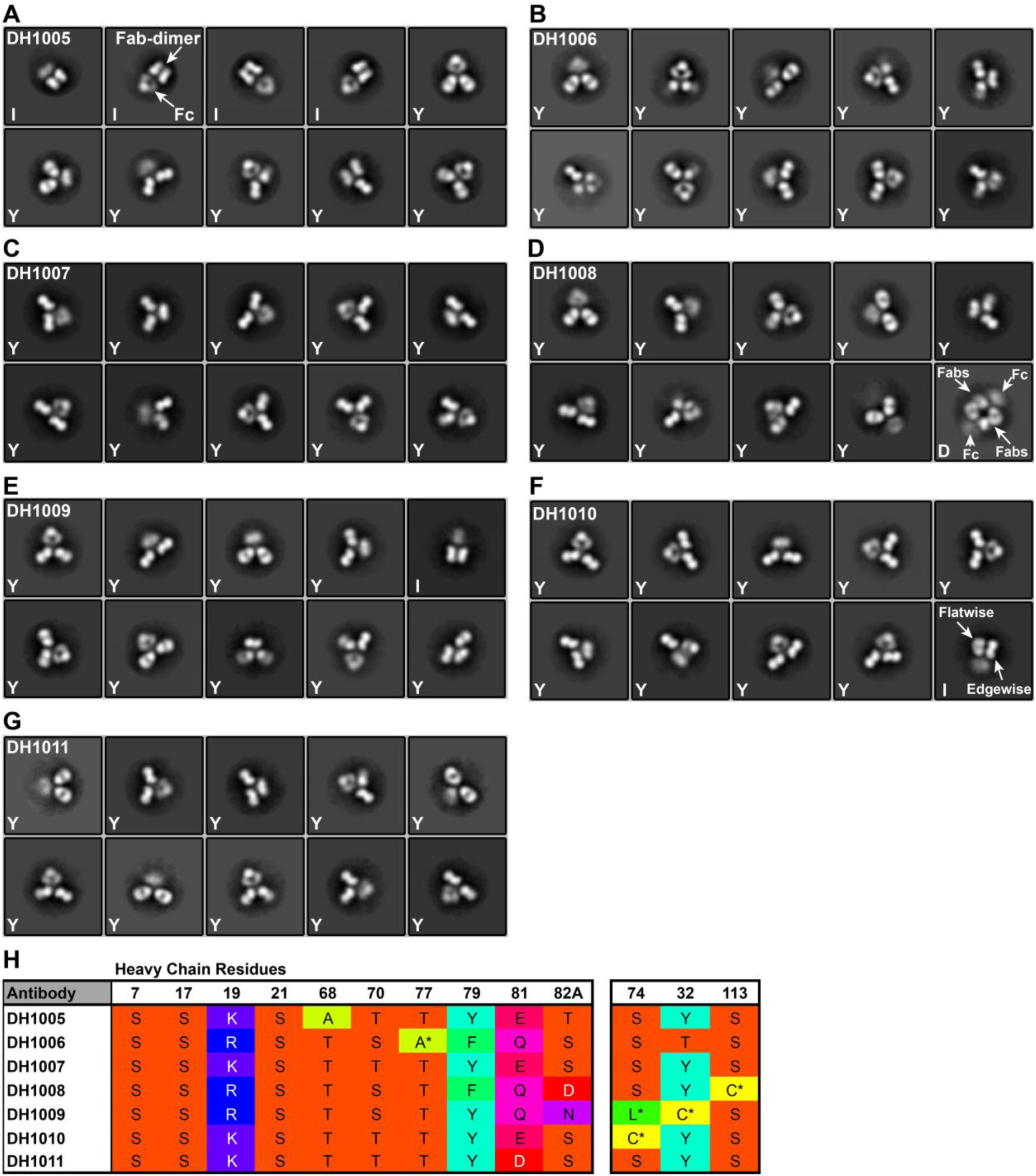
NSEM 2D class averages and sequence analysis of human antibodies DH1005-DH1011. We studied representative candidate FDG antibodies (DH1005-DH1011) that were found in a HIV-1 naïve individual (LKP04). **(A)** DH1005 class averages showed a mix of I-shaped (I) and Y-shaped (Y) antibody populations. Note that both Fabs in the DH1005 Fab-dimer appear edgewise. Thus, DH1005 represented a third type of Fab-dimerization, distinct from the side-by-side Fab dimers seen in 2G12, DH851, DH898, and DH501, and also distinct from the 90° oriented Fabs seen in DH717. **(B)** DH1006, **(C)** DH1007 and **(G)** DH1011 showed all Y-shaped antibodies. **(D)** DH1008 showed mostly Y-shaped antibody monomers and one IgG dimer (D) class at the bottom right. Here, the Fabs appeared to be angled to one another and touched at the elbow regions between the Fab variable and constant domains, in contrast to being side-by-side and touching at the V_H_ domains as seen in 2G12 IgG dimers (compare dimer here to dimers in Figure S6F and S6I). **(E)** DH1009 showed mostly Y-shaped antibody populations, but one I-shaped class (top right). **(F)** DH1010 showed mostly Y-shaped antibody populations, but one I-shaped class (bottom right). Note that this I-shaped Ab appeared with one Fab edgewise and one Fab flatwise, thus this antibody from a sero-negative human appears similar to the Fab-dimerized DH717 antibodies isolated from a vaccinated macaque. **(H)** Sequence analysis of human antibodies, DH1005-DH1011. None of the DH1005-DH1011 antibodies showed the pattern of hydrophobic residues (7 – 82A) characteristic of 2G12-like antibodies that were Fab-dimerized via the V_H_ domains, such as DH851, DH898 and DH501. Moreover, DH1005 – DH1011 all contained a positively charged arginine (R) or lysine (K) at residue 19, expected to disfavor 2G12-like side-by-side Fab-dimerization. Three antibodies, DH1008 – DH1010, showed unpaired cysteine residues and these three antibodies all showed Fab-dimerization. Of particular note, DH1010 shared the same cysteine, heavy chain residue 74, seen in the DH717 lineage, consistent with the similar appearance of the I-shaped antibody classes in the two lineages. DH1008 has a (rare) unpaired cysteine in the Fab elbow region at heavy chain residue 113, suggesting that an intermolecular disulfide linkage here drove the IgG dimer class seen in DH1008. We have not yet identified the mechanism of Fab-dimerization for DH1005.

**Figure S21.**
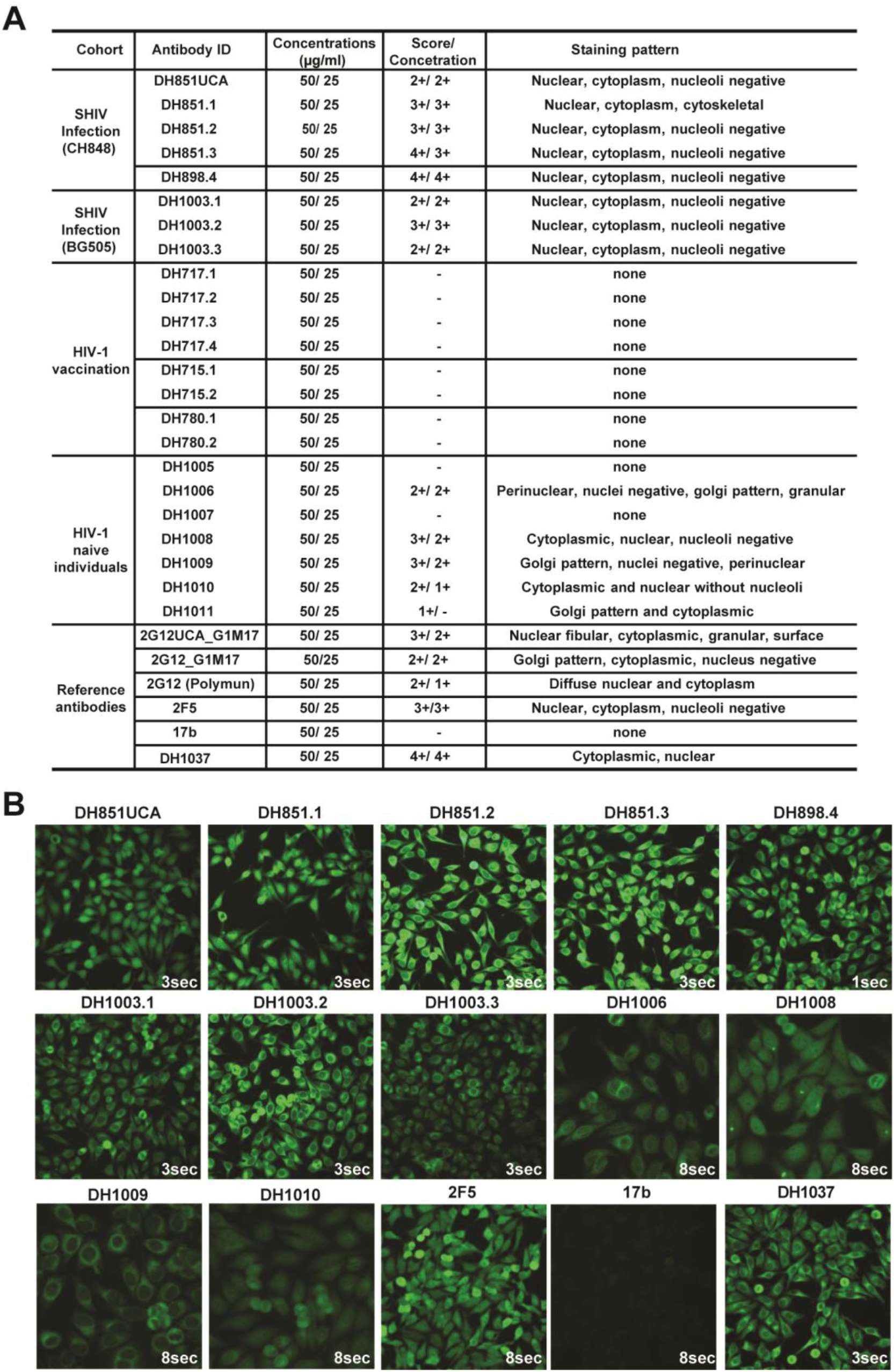
Polyreactivity profile of FDG antibodies. **(A)** Indirect immunofluorescence assay testing reactivity of FDG antibodies in HEp-2 cells. Each mAb was tested at 50 and 25 µg/ml in duplicate reactions; a representative of the replicate data were reported. The secondary antibody was a goat anti-rhesus Ig(H+L) FITC. Positivity scores were determined relative to non-human primate positive control (DH1037) and negative control (DH570.30) mAbs. Staining patterns were identified using the Zeus Scientific pattern guide. **(B)** Images of mAb staining of HEp-2 cells. Images were taken using an Olympus AX70 microscope with a SpotFlex FX1520 camera, and acquired on a 40X objective using the FITC fluorescence channel at the acquisition times indicated on the images. Antibody reactivity pattern in HEP-2 cells were described in panel A.

**Figure S22.**
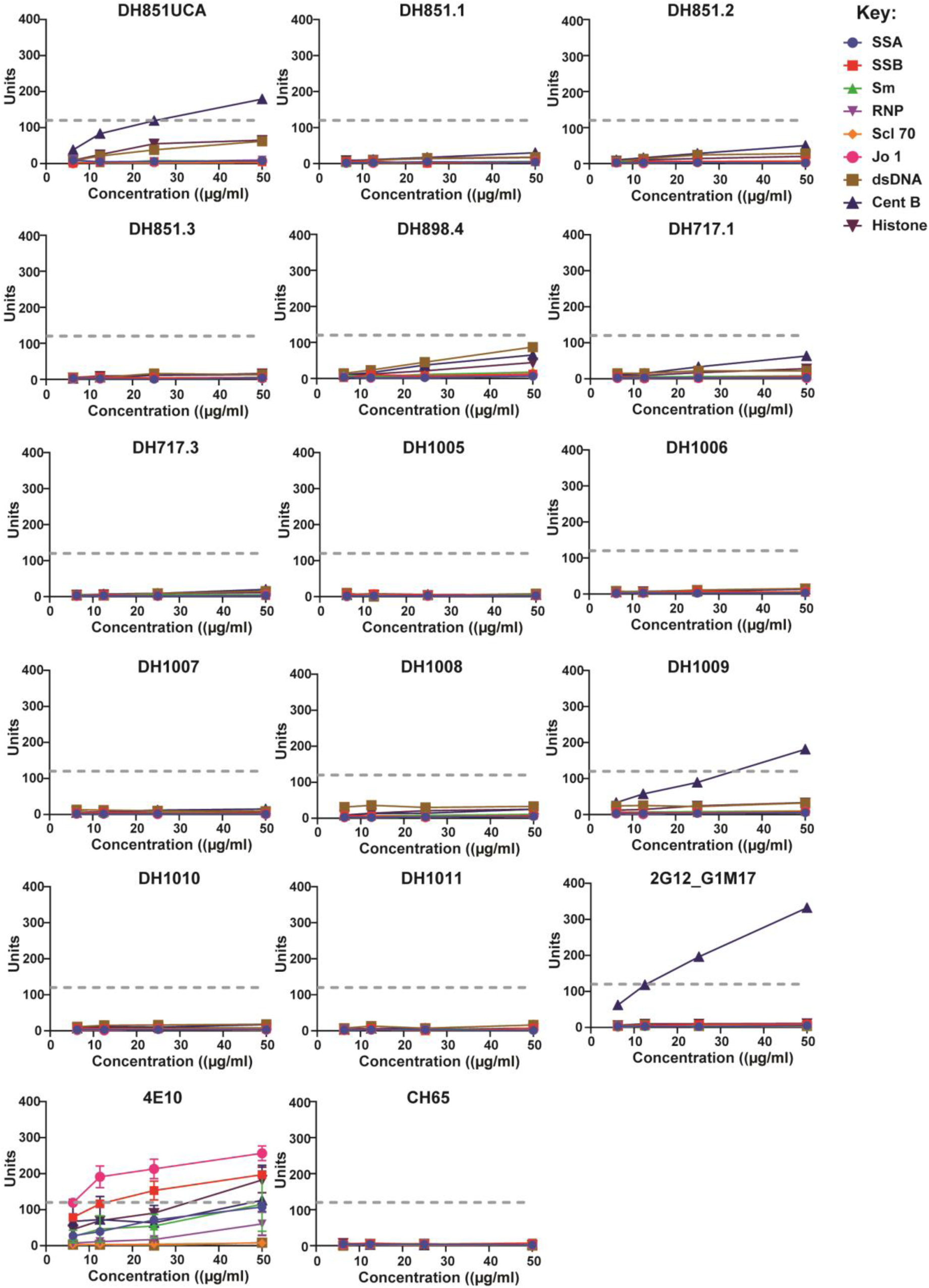
Anti-nuclear antibody (ANA) reactivity profile of FDG antibodies. Nine autoantigens were tested for reactivity by FDG antibodies using a commercially available AtheNA Multi-Lyte ANA kit (Zeus scientific, Inc.). Serially diluted mAbs were tested for binding and the data analyzed using an AtheNA software. The secondary antibody provided with the kit was Phycoerythrin-conjugated goat anti-human IgG. The dash lines represented the positivity score (121 units), which was consistent across independent experiments. 4E10 and CH65 represented positive and negative control mAbs, respectively. Data shown combined mAbs tested across independent experiments.

**Table S1.** Isolation of SHIV CH848TF-induced HIV-1 Envelope-reactive antibodies

| PTID | Sample | Timepoint (weeks) | Total antibodies | CH848 Env+ antibodies | Man<sub>9</sub>-V3 glycopeptide differential binders |
| --- | --- | --- | --- | --- | --- |
| 6163 | LN | w52 | 79 | 34 | 17 |
|  | PBMC | w52 | 22 | 20 | 1 |
|  | PBMC | w104 | 235 | 119 | 1 |
| Antibody tally: |  |  | 336 | 173 (51%) | 19 (6%) |

Single B cells from a SHIV CH848TF-infected macaque (RM6163) were sorted from lymph node (LN) and peripheral blood (PBMCs) cells at different timepoints (weeks) following infection using various sort strategies (see methods). The paired heavy and light chain genes of unique antibodies per single B cells were recovered via PCR. Recombinant IgGs bearing paired heavy and light chain genes per antibody were tested for binding gp120 and SOSIP trimer proteins, and Man<sub>9</sub>-V3 glycopeptides via ELISA. Man<sub>9</sub>-V3 glycopeptide differential binders reacted with CH848 Envs and Man<sub>9</sub>-V3 glycopeptide, but not the corresponding non-glycosylated aglycone V3 peptide. This assay represents the initial binding screen for antibodies and informs selection of monoclonal antibodies that are expressed in large scale transfection for complete functional characterization.

**Table S2.**
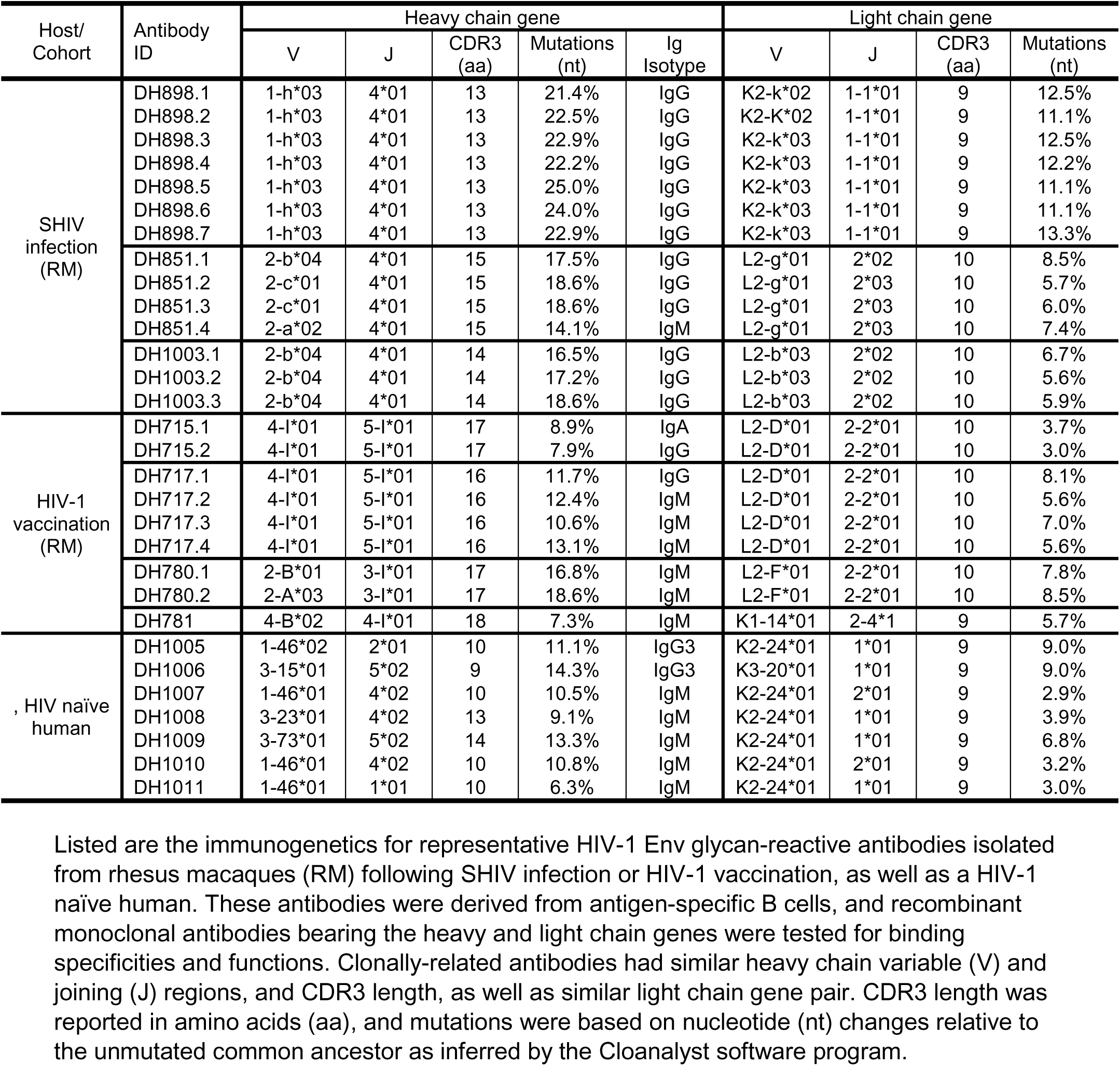
Immunogenetics of macaque and human HIV-1 Envelope glycan-reactive antibodies

**Table S3. Neutralization breadth profile of SHIV CH848TF-induced FDG antibodies, DH851 and DH898.** Neutralization breadth was tested against a panel of 119 heterologous tier 2 HIV-1 strains in TZM-bl cells. We tested representative FDG mAbs, DH851.1-DH851.3 and DH898.4, and control antibody 2G12. Data shown are neutralization titers (IC50 and IC80) in µg/ml, and neutralization breadth calculated as a percent of viruses neutralized with IC50 titers. See excel spreadsheet (Table S3).

**Table S4.** DH851.3 data collection and reconstruction statistics for complex with HIV Env CH505TF.6R.chim.SOSIP.664.v4.1.

|  |  |
| --- | --- |
| **Data Collection** |  |
| Microscope | FEI Titan Krios |
| Voltage (kV) | 300 |
| Electron dose (e<sup>-</sup>/Å<sup>2</sup>) | 60 |
| Detector | Gatan K3 |
| Pixel Size (Å) | 1.07 |
| Defocus Range (µm) | ~0.8-3.5 |
| Magnification | 22500 |
| **Reconstruction** |  |
| Software | cryoSPARC |
| Particles | 136,877 |
| Symmetry | C1 |
| Box size (pixels) | 384 |
| Resolution (Å)<sup>\$</sup> | 4.5 |

<sup>\$</sup>Resolution reported by cryoSPARC as corrected gold standard Fourier shell correlation at 0.143 level.

**Table S5.** DH898.1 data collection and reconstruction statistics for complex with HIV Env CH848.3.D0949.10.17chim.6R.DS.SOSIP.664.

|  | CD4-binding-site glycans |  | V3-glycans |  | apo-Trimer |
| --- | --- | --- | --- | --- | --- |
|  | Complex | Fab-dimer* | Complex | Fab-dimer* |  |
| Data Collection |  |  |  |  |  |
| Microscope | FEI Titan Krios | FEI Titan Krios | FEI Titan Krios | FEI Titan Krios | FEI Titan Krios |
| Voltage (kV) | 300 | 300 | 300 | 300 | 300 |
| Electron dose (e<sup>-</sup>/Å<sup>2</sup>) | 60 | 60 | 60 | 60 | 60 |
| Detector | Gatan K3 | Gatan K3 | Gatan K3 | Gatan K3 | Gatan K3 |
| Pixel Size (Å) | 1.07 | 1.07 | 1.07 | 1.07 | 1.07 |
| Defocus Range (µm) | ~0.8-3.5 | ~0.8-3.5 | ~0.8-3.5 | ~0.8-3.5 | ~0.8-3.5 |
| Magnification | 22500 | 22500 | 22500 | 22500 | 22500 |
| Reconstruction |  |  |  |  |  |
| Software | cryoSPARC | cryoSPARC | cryoSPARC | cryoSPARC | cryoSPARC |
| Particles | 56,474 | 56,474 | 14,930 | 14,930 | 56,281 |
| Symmetry | C1 | C1 | C1 | C1 | C3 |
| Box size (pixels) | 384 | 384 | 384 | 384 | 384 |
| Resolution (Å)<sup>\$</sup> | 4.8 | 4.6 | 6.9 | 6.9 | 4.8 |

\*Local refinement of the Fab-dimer alone was performed after masking and signal subtraction of density corresponding to the Env trimer, per standard cryoSPARC protocols.
<sup>§</sup>Resolution reported by cryoSPARC as corrected gold standard Fourier shell correlation at 0.143 level.

**Table S6.**
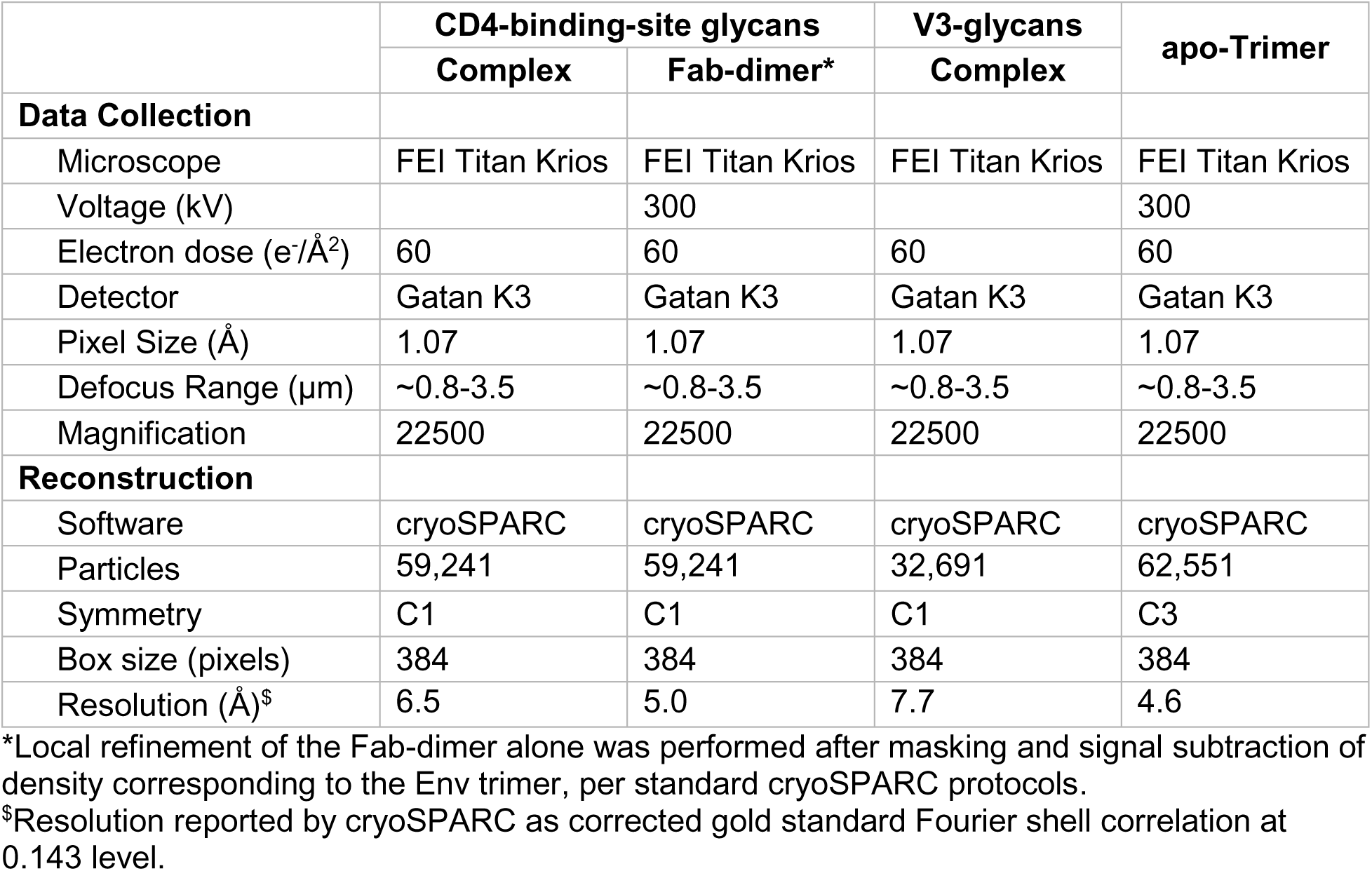
DH898.4 data collection and reconstruction statistics for complex with HIV Env CH848.3.D0949.10.17chim.6R.DS.SOSIP.664.

|  | CD4-binding-site glycans |  | V3-glycans | apo-Trimer |
| --- | --- | --- | --- | --- |
|  | Complex | Fab-dimer* | Complex |  |
| Data Collection |  |  |  |  |
| Microscope | FEI Titan Krios | FEI Titan Krios | FEI Titan Krios | FEI Titan Krios |
| Voltage (kV) |  | 300 |  | 300 |
| Electron dose (e<sup>-</sup>/Å<sup>2</sup>) | 60 | 60 | 60 | 60 |
| Detector | Gatan K3 | Gatan K3 | Gatan K3 | Gatan K3 |
| Pixel Size (Å) | 1.07 | 1.07 | 1.07 | 1.07 |
| Defocus Range (µm) | ~0.8-3.5 | ~0.8-3.5 | ~0.8-3.5 | ~0.8-3.5 |
| Magnification | 22500 | 22500 | 22500 | 22500 |
| Reconstruction |  |  |  |  |
| Software | cryoSPARC | cryoSPARC | cryoSPARC | cryoSPARC |
| Particles | 59,241 | 59,241 | 32,691 | 62,551 |
| Symmetry | C1 | C1 | C1 | C3 |
| Box size (pixels) | 384 | 384 | 384 | 384 |
| Resolution (Å)<sup>\$</sup> | 6.5 | 5.0 | 7.7 | 4.6 |

\*Local refinement of the Fab-dimer alone was performed after masking and signal subtraction of density corresponding to the Env trimer, per standard cryoSPARC protocols.
§Resolution reported by cryoSPARC as corrected gold standard Fourier shell correlation at 0.143 level.

**Table S7.**
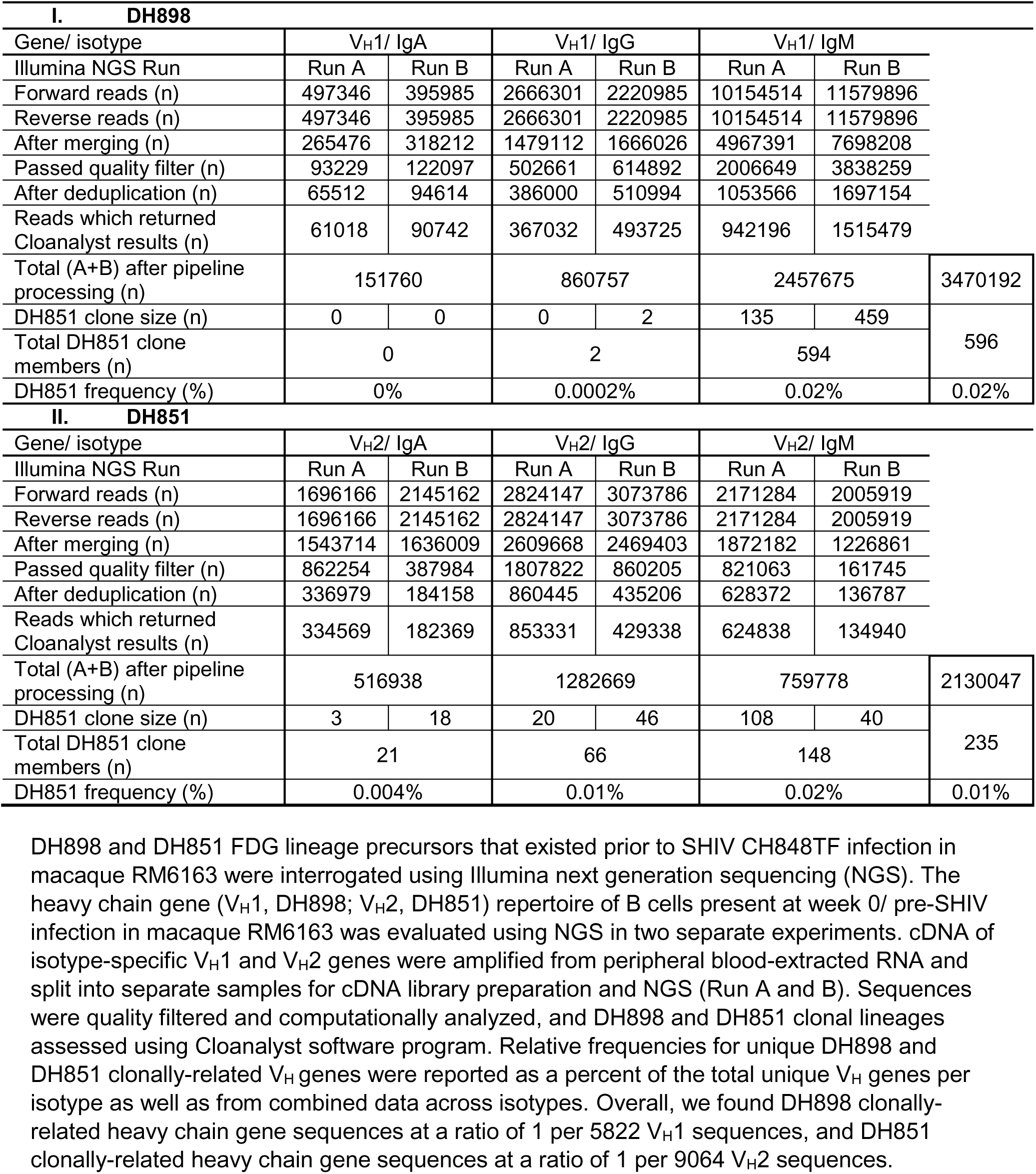
Interrogation of macaque FDG lineage precursors prior to SHIV CH848TF infection

| I. DH898 |  |  |  |  |  |  |  |
| --- | --- | --- | --- | --- | --- | --- | --- |
| Gene/ isotype | V<sub>H</sub>1/ IgA |  | V<sub>H</sub>1/ IgG |  | V<sub>H</sub>1/ IgM |  |  |
| Illumina NGS Run | Run A | Run B | Run A | Run B | Run A | Run B |  |
| Forward reads (n) | 497346 | 395985 | 2666301 | 2220985 | 10154514 | 11579896 |  |
| Reverse reads (n) | 497346 | 395985 | 2666301 | 2220985 | 10154514 | 11579896 |  |
| After merging (n) | 265476 | 318212 | 1479112 | 1666026 | 4967391 | 7698208 |  |
| Passed quality filter (n) | 93229 | 122097 | 502661 | 614892 | 2006649 | 3838259 |  |
| After deduplication (n) | 65512 | 94614 | 386000 | 510994 | 1053566 | 1697154 |  |
| Reads which returned Cloanalyst results (n) | 61018 | 90742 | 367032 | 493725 | 942196 | 1515479 |  |
| Total (A+B) after pipeline processing (n) | 151760 |  | 860757 |  | 2457675 |  | 3470192 |
| DH851 clone size (n) | 0 | 0 | 0 | 2 | 135 | 459 | 596 |
| Total DH851 clone members (n) | 0 |  | 2 |  | 594 |  |  |
| DH851 frequency (%) | 0% |  | 0.0002% |  | 0.02% |  | 0.02% |
| II. DH851 |  |  |  |  |  |  |  |
| Gene/ isotype | V<sub>H</sub>2/ IgA |  | V<sub>H</sub>2/ IgG |  | V<sub>H</sub>2/ IgM |  |  |
| Illumina NGS Run | Run A | Run B | Run A | Run B | Run A | Run B |  |
| Forward reads (n) | 1696166 | 2145162 | 2824147 | 3073786 | 2171284 | 2005919 |  |
| Reverse reads (n) | 1696166 | 2145162 | 2824147 | 3073786 | 2171284 | 2005919 |  |
| After merging (n) | 1543714 | 1636009 | 2609668 | 2469403 | 1872182 | 1226861 |  |
| Passed quality filter (n) | 862254 | 387984 | 1807822 | 860205 | 821063 | 161745 |  |
| After deduplication (n) | 336979 | 184158 | 860445 | 435206 | 628372 | 136787 |  |
| Reads which returned Cloanalyst results (n) | 334569 | 182369 | 853331 | 429338 | 624838 | 134940 |  |
| Total (A+B) after pipeline processing (n) | 516938 |  | 1282669 |  | 759778 |  | 2130047 |
| DH851 clone size (n) | 3 | 18 | 20 | 46 | 108 | 40 | 235 |
| Total DH851 clone members (n) | 21 |  | 66 |  | 148 |  |  |
| DH851 frequency (%) | 0.004% |  | 0.01% |  | 0.02% |  | 0.01% |

DH898 and DH851 FDG lineage precursors that existed prior to SHIV CH848TF infection in macaque RM6163 were interrogated using Illumina next generation sequencing (NGS). The heavy chain gene (V<sub>H</sub>1, DH898; V<sub>H</sub>2, DH851) repertoire of B cells present at week 0/ pre-SHIV infection in macaque RM6163 was evaluated using NGS in two separate experiments. cDNA of isotype-specific V<sub>H</sub>1 and V<sub>H</sub>2 genes were amplified from peripheral blood-extracted RNA and split into separate samples for cDNA library preparation and NGS (Run A and B). Sequences were quality filtered and computationally analyzed, and DH898 and DH851 clonal lineages assessed using Cloanalyst software program. Relative frequencies for unique DH898 and DH851 clonally-related V<sub>H</sub> genes were reported as a percent of the total unique V<sub>H</sub> genes per isotype as well as from combined data across isotypes. Overall, we found DH898 clonally-related heavy chain gene sequences at a ratio of 1 per 5822 V<sub>H</sub>1 sequences, and DH851 clonally-related heavy chain gene sequences at a ratio of 1 per 9064 V<sub>H</sub>2 sequences.

**Table S8.** DH717 data collection and refinement statistics (X-ray crystallography)

| **Data Collection** |  |  |
| --- | --- | --- |
|  | **DH717 – Man<sub>9</sub>-V3** | **DH717.1 dimer** |
| Resolution (Å) | 50.00 – 2.61 (2.66 - 2.61) | 41.71 - 3.14 (3.26 - 3.14) |
| Space group | P 4<sub>1</sub> 2<sub>1</sub> 2 | P 4<sub>3</sub> 2 2 |
| Unit cell<br>a,b,c (Å) | 115.2 115.2 98.5 | 201.6, 201.6, 154.3 |
| Unit cell<br>$\alpha, \beta, \gamma$ (°) | 90, 90, 90 | 90, 90, 90 |
| Total reflections | 90397 | 497617 |
| Unique reflections | 21131 (1949) | 55471 (2730) |
| Redundancy | 4.4 (3.3) | 9.0 (9.6) |
| Completeness (%) | 97.9 (92.42) | 99.9 (100.0) |
| $\langle I/\sigma \rangle$ | 7.5 (1.4) | 13.4 (1.3) |
| Wilson B-factor | 45.4 | 82.5 |
| R<sub>merge</sub> (%) | 14.9 (88.4) | 16.6 (228.6) |
| CC1/2 | 0.984 (0.505) | 0.992 (0.525) |
| **Refinement** |  |  |
| Reflections used in refinement | 21126 (1949) | 55412 (5393) |
| Reflections used for R-free | 1045 (107) | 1999 (195) |
| R<sub>work</sub>/R<sub>free</sub> (%) | 19.1/22.4<br>(29.8/29.3) | 23.0/27.9<br>(29.7/32.5) |
| No. atoms | 3364 | 12965 |
| Protein | 3170 | 12965 |
| Ligand | 39 | 0 |
| Water | 155 | 0 |
| **R.M.S. deviations** |  |  |
| Bond lengths (Å) | 0.004 | 0.09 |
| Bond angles (°) | 0.88 | 2.51 |
| **B-factors (Å<sup>2</sup>)** |  |  |
| Protein | 46.70 | 101.5 |
| Ligand | 68.90 | NA |
| Solvent | 43.60 | NA |
| **Ramachandran Plot** |  |  |
| Favored (%) | 96.0 | 93.09 |
| Allowed (%) | 3.53 | 5.28 |
| Rotamer outliers (%) | 4.30 | 0.14 |
| Clashscore | 6.49 | 8.99 |

Statistics for the highest-resolution shell are shown in parentheses.

**Table S9. Interrogation of macaque FDG lineage precursors prior to HIV vaccination.** Data shown summarizes longitudinal next generation sequencing (NGS) analysis of macaque B cells. The heavy chain (V_H_4) repertoire of B cells present at longitudinal timepoints (weeks) were evaluated using Illumina NGS. cDNA of isotype-specific V_H_4 genes were amplified from PBMC-extracted RNA and split into separate samples for cDNA library preparation and NGS (Run A and B). Sequences were quality filtered and computationally analyzed. DH715 and DH717 clonal lineages were assessed using the Cloanalyst software program. See excel spreadsheet (Table S9).

